# Shell protein composition specified by NEAT1 domains dictates the formation of paraspeckles as distinct membraneless organelles

**DOI:** 10.1101/2023.05.21.541661

**Authors:** Hiro Takakuwa, Tomohiro Yamazaki, Sylvie Souquere, Shungo Adachi, Hyura Yoshino, Naoko Fujiwara, Tetsuya Yamamoto, Tohru Natsume, Shinichi Nakagawa, Gerard Pierron, Tetsuro Hirose

## Abstract

Many membraneless organelles (MLOs) formed through phase separation play crucial roles in various cellular processes. Although these MLOs co-exist in cells, how they maintain their independence without coalescence or engulfment remains largely unknown. Here, we investigated the molecular mechanism by which paraspeckles with core-shell architecture scaffolded by NEAT1_2 lncRNAs exist as distinct MLOs. We identified NEAT1 deletion mutants that assemble paraspeckles that are incorporated into nuclear speckles. Several paraspeckle proteins, including SFPQ, HNRNPF, and BRG1, prevent this incorporation and thus contribute to the segregation of paraspeckles from nuclear speckles. Shell localization of these proteins in the paraspeckles, which is determined by NEAT1_2 lncRNA domains, is required for this segregation process. Conversely, U2-related spliceosomal proteins are involved in internalizing the paraspeckles into nuclear speckles. This study shows that the paraspeckle shell composition dictates the independence of MLOs in the nucleus, providing insights into the importance of the shell in defining features and functions of MLOs.

## Introduction

Membraneless organelles (MLOs), also known as cellular bodies, form through phase separation and play critical roles in various cellular processes by creating intracellular compartments concentrating specific biomolecules^1–5^. Many MLOs are located in the interchromatin spaces in the highly organized nucleus. Nuclear MLOs include nucleoli, Cajal bodies, nuclear speckles (NSs), paraspeckles (PSs), PML nuclear bodies, and histone locus bodies, which are involved in gene expression and the formation of gene expression machinery^1, 4^. Such MLOs typically contain proteins and nucleic acids. RNAs are major constituents of MLOs and contribute to their formation by facilitating molecular interactions^6–8^.

Long noncoding RNAs (lncRNAs) are fundamental regulators of cellular processes, such as gene expression and chromatin organization. A subset of lncRNAs termed architectural RNAs (arcRNAs) function as the scaffolds of MLOs^9, 10^. NEAT1_2 lncRNA serves as an essential architectural scaffold of PSs by concentrating many RNA-binding proteins^11–14^. PSs were initially defined as the nuclear domains found in close proximity to NSs and are enriched with the characteristic DBHS (Drosophila Behavior Human Splicing) family proteins, SFPQ, NONO, and PSPC1^15, 16^. PSs play key roles in physiological and pathological conditions, including the establishment of pregnancy, cancer, and virus infection^17–21^. NEAT1 is transcribed by RNA polymerase II (Pol II) to produce two distinct isoforms, NEAT1_1 (3.7 kb) and NEAT1_2 (22.7 kb), as a consequence of alternative 3’ end processing^22^. NEAT1_2 but not NEAT1_1 is indispensable for PS formation^22, 23^. NEAT1_1 is canonically polyadenylated, whereas NEAT1_2 contains a triple helix structure at the 3’ terminus essential for its stabilization^14, 24^. The PS forms near the NEAT1 gene locus by coupling with NEAT1_2 transcription, and PSs are quickly disintegrated after transcription inhibition by Pol II inhibitors^25, 26^. More than 60 PS proteins (PSPs) are enriched in PSs, and some of these PSPs, such as SFPQ, NONO, FUS, RBM14, and the SWI/SNF complex, are essential for the PS formation processes^22, 27, 28^. SFPQ and NONO proteins are required for the expression of NEAT1_2 lncRNA, and NONO, FUS, RBM14, and the SWI/SNF complex proteins are required for PS assembly^22, 23, 27, 29^. We have analyzed phenotypes of PSs in many cell lines expressing NEAT1 deletion mutants created by using CRISPR/Cas9^23, 30, 31^. We revealed that NEAT1_2 has modular functional RNA domains for RNA stabilization, isoform switching from NEAT1_1 to NEAT1_2, and PS assembly^23^. The middle domain of NEAT1_2 contains multiple binding sites for NONO and SFPQ that are necessary and sufficient for PS assembly through phase separation^23^. The abilities of the proteins for phase separation depend on their self-assembly domains, including the RRM2/NOPS and coiled-coil domains^23^. The PSs possess characteristic core-shell internal organization and show cylindrical as well as spherical shapes^32^. PSPs also show specific patterns of localization within PSs^33^. These characteristic shapes and the internal organization of PSs are distinct from those of the typical MLOs formed by liquid-liquid phase separation (LLPS), which are spherical and have unordered internal structures. A recent study using soft matter physics theories showed that PSs are block copolymer micelles assembled through micellization^31, 34^. Thus, NEAT1_2 lncRNA encodes the form and function of the PSs, but the rules underlying this remain incompletely understood.

Many MLOs co-exist independently in intracellular space, some of which are in close proximity to each other. Although PSs localize in close proximity to NSs, it is unknown how they are immiscible, avoiding coalescence or engulfment. We here investigated the molecular mechanism behind this immiscibility and identified proteins that control the localization of PSs inside and outside of NSs. Our study revealed that specific shell-localizing proteins specified by NEAT1_2 RNA domains determine the independence of PSs from NSs, providing insights into the general molecular mechanisms governing the independence of MLOs.

## Results

### PSs constructed with a NEAT1_2 mutant are incorporated into NSs

NEAT1 deletion analyses have elucidated important mechanisms of PS formation^23, 30, 31^ with deletion of each of the modular NEAT1 domains causing defects in the specific properties of the PSs without affecting the other properties. During detailed analyses of many NEAT1 deletion mutants, we unexpectedly found that the PSs (named mini-PSs) constructed by mini-NEAT1, which contains the NEAT1_2 middle domain, 5’ domain (0– 1 kb), and the triple helix and forms PSs with core-shell internal organization^23^ (Fig. 1a and Extended Data Fig. 1a), were present within NSs. NEAT1 RNA-FISH and immunofluorescence (IF) of SRRM2 as a marker for NS revealed that about 90% of mini-PSs colocalized with NSs, while WT-PSs did not colocalize with NSs as expected (Fig. 1b,c, Extended Data Fig. 1b, c and Supplementary Video 1,2). This colocalization of mini-PS was validated in another mini-NEAT1 cell clone and by using other NS markers, SON and MALAT1 lncRNA (Extended Data Fig. 1d–h). Using a super-resolution microscope (SRM), the internalization of mini-PSs into NSs was observed^23^ (Fig. 1d,e), confirming initial electron microscope (EM) observations indicating that almost all of the immunogold-labeled mini-PSs, which have electron-dense structures similar to WT-PSs and are smaller than WT-PSs as previously reported^23^, were found within the ultrastructurally defined NSs (Fig. 1f and Extended Data Fig. 1i). We did not observe any colocalization of mini-PSs with nuclear bodies visible under EM. In addition, no colocalization of mini-PSs with several nuclear bodies was found by confocal microscopy (Extended Data Fig. 1j), suggesting that mini-PSs are specifically incorporated into NSs, but not into the other nuclear bodies. These findings together show that, although the mini-PSs have a core-shell organization and are immiscible, they are occluded by NSs and thus lack the ability to exist as distinct MLOs.

**Fig. 1.**
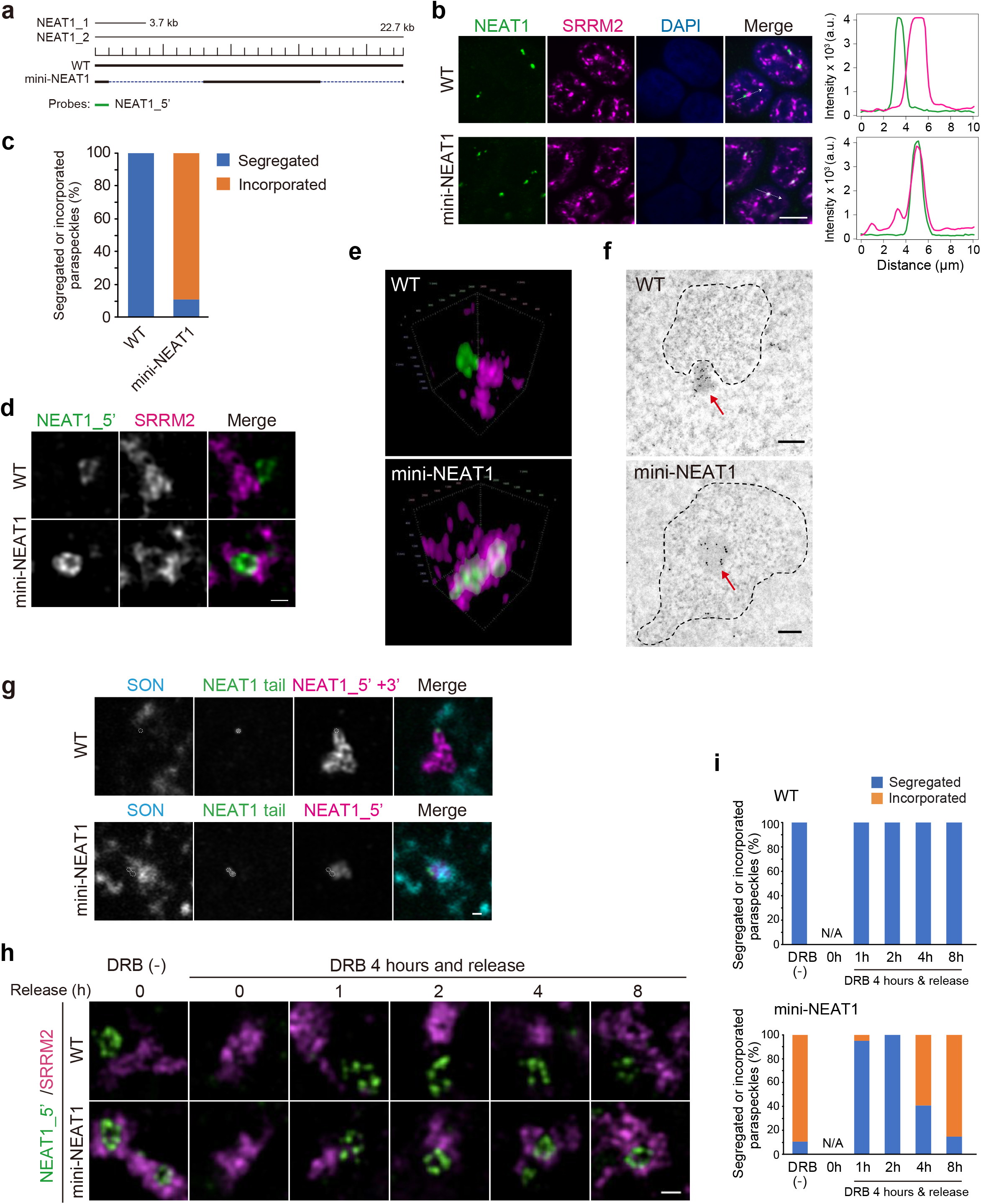
Mini-PSs are incorporated into NSs. **a**, Schematics of the WT and mini-NEAT1 mutant are shown with a scale. The position of a NEAT1_5’ probe used in SRM (green) is shown below. **b**, PSs and NSs in HAP1 WT and mini-NEAT1 cells were detected by NEAT1 RNA-FISH (green) and immunofluorescence (IF) of SRRM2 (magenta) in the presence of MG132 (5 μM for 6 h), which induces NEAT1_2 expression^17^. Nuclei were stained with DAPI. Line profiles of the PSs and NSs are shown (right). Scale bar, 10 μm. **c**, Graph showing the proportion of segregated or incorporated PSs in WT and mini-NEAT1 cells treated with MG132 (5 μM for 6 h) (WT: *n* = 370, mini-NEAT1: *n* = 409). **d**, SIM images of PSs (RNA-FISH with NEAT1_5’ probe) and NSs (SRRM2 IF) in HAP1 WT and mini-NEAT1 cells treated with MG132 (5 μM for 6 h). Scale bar, 500 nm. **e**, 3D SIM images of PSs and NSs in HAP1 WT and mini-NEAT1 cells treated with MG132 (5 μM for 6 h) (corresponding to Supplementary Videos 1, 2). **f**, EM observation of PSs and NSs in HAP1 WT and mini-NEAT1. Localization of NONO was detected as gold particles. Red arrows indicate the position of the PSs. Dashed black circles indicate the position of the NS. Scale bar, 200 nm. **g,** SRM images of NEAT1 transcription site (RNA-FISH with NEAT1_tail probe)^26, 33^, PSs (RNA-FISH with NEAT1_5’ probe), and NSs (SRRM2 IF) in HAP1 WT and mini-NEAT1 cells treated with MG132 (5 μM for 6 h). Scale bar, 500 nm. **h**, SIM images of de novo synthesized PSs in HAP1 WT and mini-NEAT1 cells. The cells were treated with DRB (30 μM) for 4 h and then released for 1, 2, 4, and 8 h. The PSs were detected by NEAT1_5’ RNA-FISH and the NSs were detected by SRRM2 IF. Scale bar, 500 nm. **i**, Graph showing the proportion of segregated or incorporated PSs in the cells in **h.** Data were collected from three independent experiments: WT: *n* = 64 (DRB, 0 h), *n* = 60 (Release, 1 h), *n* = 60 (Release, 2 h), *n* = 60 (Release, 4 h), *n* = 62 (Release, 8 h); mini-NEAT1: *n* = 67 (DRB, 0 h), *n* = 60 (Release, 1 h), *n* = 59 (Release, 2 h), *n* = 62 (Release, 4 h), *n* = 59 (Release, 8 h). Numerical data are available in the Source Data.

Although mini-PSs were clearly present within NSs, it was unclear whether de novo synthesized mini-PSs formed inside or outside of NSs. To investigate this, we first examined whether the transcription sites of NEAT1, where PSs are formed, are inside or outside of NSs. We visualized the transcription sites by detecting nascent NEAT1 transcripts using NEAT1 tail probes, which detect unprocessed NEAT1 downstream regions, as used in a previous study^33^ (Fig. 1g). The NEAT1 transcription sites were located outside and adjacent to NSs in both HAP1 WT and mini-NEAT1 cells (Fig. 1g), suggesting that mini-PSs are formed outside NSs. Furthermore, we observed de novo PS formation over time after release from a reversible Pol II inhibitor, 5,6-dichloro-1-β-D-ribofuranosylbenzimidazole (DRB), which inhibits PS formation^23, 25, 35^. After 4 h of DRB treatment, PSs were completely diminished. At 1–2 h after release, PSs started to re-form in both WT and mini-NEAT1 cell lines, and most of them were present adjacent to NSs (Fig. 1h,i). At 4–8 h after release, the localization of mini-PSs within NSs gradually increased (Fig. 1 h,i). Thus, even when we created a starting point where mini-PSs are present outside the NSs, mini-PSs eventually enter them. These findings suggest that mini-PSs form outside NSs and are then incorporated into them.

### Identification of NEAT1_2 RNA domains required for segregation of PSs from NSs

mini-NEAT1 lacks large 5’ and 3’ regions (1–8 kb and 16.6–22.6 kb). To precisely identify the RNA domains of NEAT1 required for PS segregation from NSs, we established a series of NEAT1 deletion mutant cell lines (Extended Data Fig. 2a,b). The τι1–8 kb/16.6–20.2 kb mutant showed the incorporation of PSs into NSs, while deletion of neither the 1–8 kb region nor the 16.6–20.2 kb region affected the localization of the PSs near NSs (Extended Data Fig. 2c,d). These findings suggest that the 1–8 kb and 16.6–20.2 kb regions are functionally redundant subdomains for PS segregation.

**Fig. 2.**
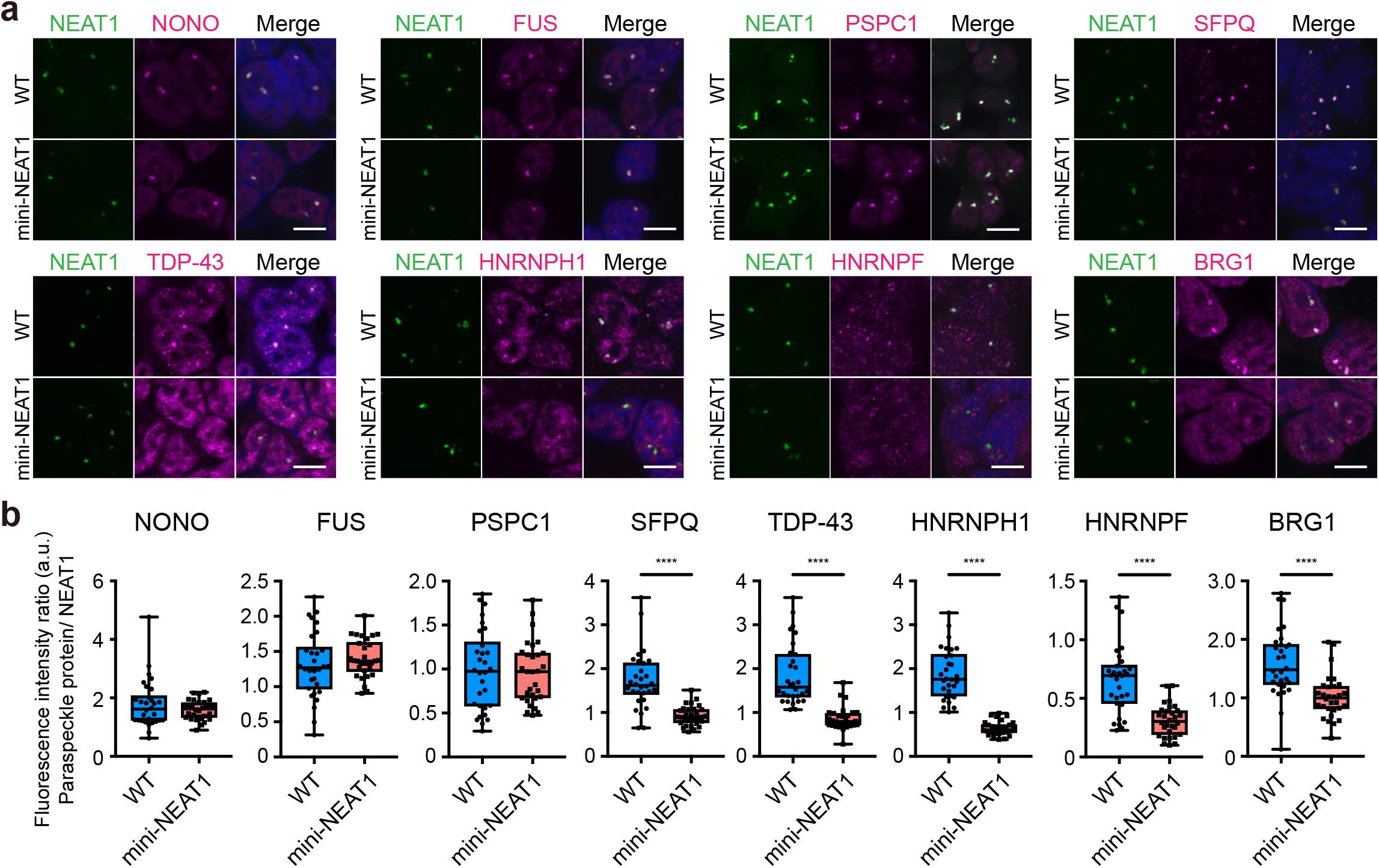
A subset of PSPs are mislocalized in mini-PSs **a**, Confocal observation of the PSs and various PSPs in HAP1 WT and mini-NEAT1 mutant cells treated with MG132 (5 μM for 6 h). PSs and NSs were visualized by RNA-FISH using NEAT1_5’ probes (green) and IF (magenta). Nuclei were stained with DAPI. Scale bar, 10 μm. **b**, Quantification of fluorescence intensity ratio (PSP/NEAT1) in the HAP1 WT (blue) and mini-NEAT1 (pink) cells as observed in **a** (all samples: *n* = 30). Each box plot shows the median (inside line), 25th-75th percentiles (box bottom to top), and minimum-maximum values (whisker bottom to top). \*\*\*\**P* < 0.0001. Data were compared using Mann-Whitney U-test. If the statistical test showed no significant difference (*P* > 0.05), it is not specified in the figure. Numerical data are available in the Source Data.

### A subset of PSPs are mislocalized in mini-PSs

Our previous study showed that multiple major PSPs such as NONO, FUS, and RBM14 are recruited to mini-PS, but SFPQ was not efficiently recruited to mini-PS compared with that to WT-PS^23^. We thus hypothesized that PSPs mislocalized in mini-PS were involved in the PS segregation defect from NSs. By further searching for mislocalized proteins, we found strong reductions in the recruitment of TDP-43, HNRNPH1, HNRNPF, and BRG1 as well as SFPQ to mini-PS (Fig. 2a,b), although the expression levels of these proteins were comparable in the WT and mini-NEAT1 cell lines (Extended Data Fig. 3a). Meanwhile, NONO, PSPC1, FUS, RBM14, HNRNPA1, and HNRNPH3 were normally recruited to both mini-PS and WT-PS (Fig. 2a,b and Extended Data Fig. 3b,c). These findings suggest that SFPQ, TDP-43, HNRNPH1, HNRNPF, and BRG1 are strong candidate proteins to segregate PSs from NSs.

**Fig. 3.**
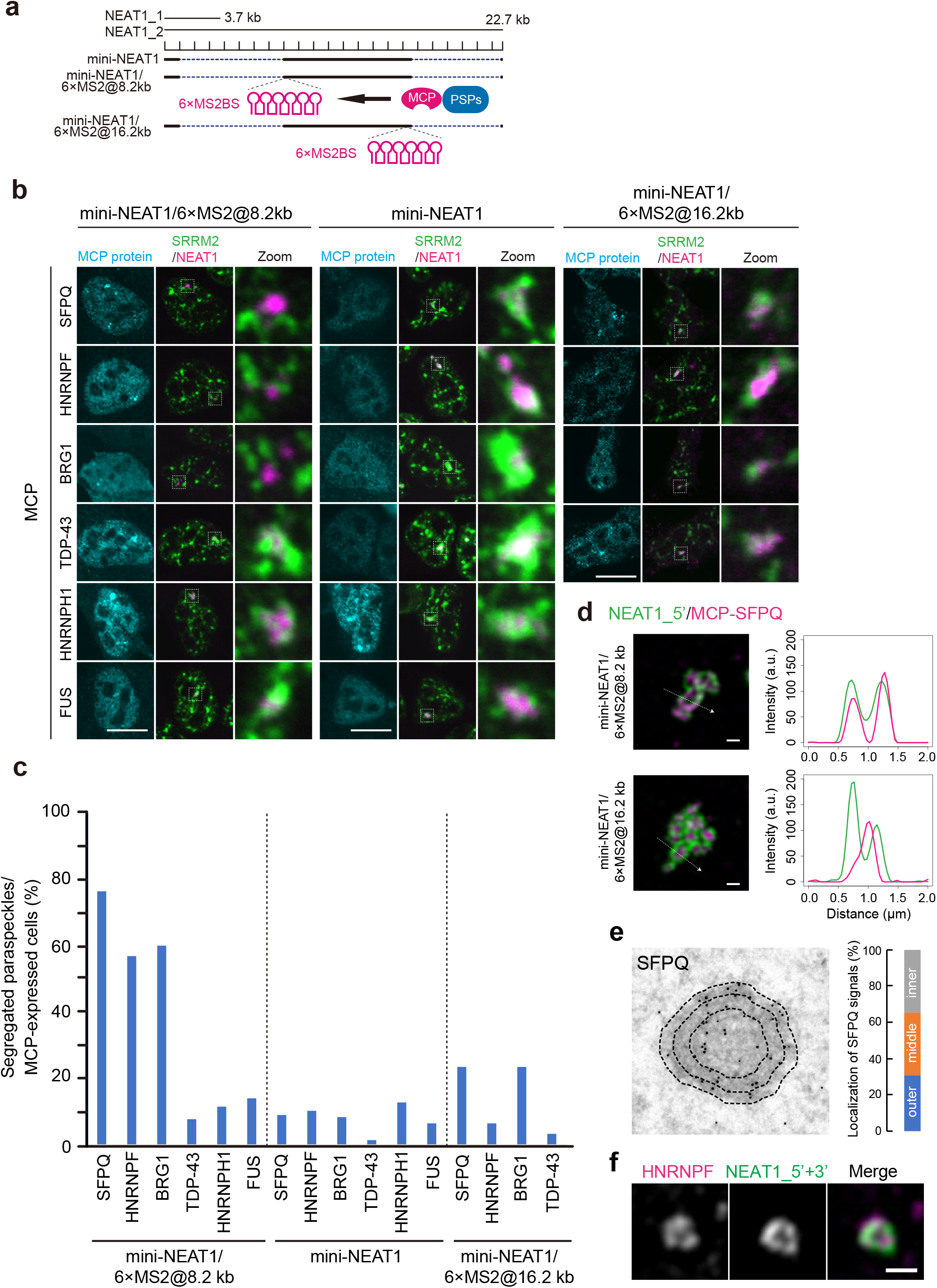
Artificial tethering of SFPQ, HNRNPF, and BRG1 to the shells induces segregation of mini-PS from NSs. **a**, Schematics of the mini-NEAT1 cell lines with 6× MS2BS for MCP-protein tethering experiments to the shell or core of PSs. **b**, Confocal observation of PSs and NSs with transfection of MCP-PSPs into mini-NEAT1/6×MS2@8.2 kb (left), mini-NEAT1 (middle), or mini-NEAT1/6×MS2@16.2 kb (right) in the MG132 treatment conditions (5 μM for 6 h). White boxes indicate the areas shown at a higher magnification. **c**, Proportion (%) of the cells with segregated PSs from NSs in **b**. Data were collected from three independent experiments. mini-NEAT1/6×MS2@8.2 kb: *n* = 68 (MCP-SFPQ), *n* = 65 (MCP-HNRNPF), *n* = 60 (MCP-BRG1), *n* = 62 (HNRNPH1-MCP), *n* = 63 (MCP-TDP-43), *n* = 63 (MCP-FUS), *n* = 63 (MCP-NONO), *n* = 61 (MCP-PSPC1); mini-NEAT1: *n* = 64 (MCP-SFPQ), *n* = 68 (MCP-HNRNPF), *n* = 60 (MCP-BRG1), *n* = 62 (HNRNPH1-MCP), *n* = 60 (MCP-TDP-43), *n* = 60 (MCP-FUS). Mini-NEAT1/6×MS2@16.2 kb: *n* = 60 (MCP-SFPQ), *n* = 60 (MCP-HNRNPF), *n* = 60 (MCP-BRG1), *n* = 60 (MCP-TDP-43). **d**, SRM images of the PSs with transfection of MCP-SFPQ into mini-NEAT1/6×MS2@8.2 kb (upper) or mini-NEAT1/6×MS2@16.2 kb (lower) corresponding to the results in **b**. Line profiles of the PSs are shown (right). Scale bar, 500 nm. **e**, EM observation of the PS and SFPQ in HAP1 WT (left). Localization of SFPQ was detected as gold particles. Dashed black circles indicate outer, middle, and inner regions of the PS. Graph showing the proportion of localization of SFPQ (492 gold particles) in the outer, middle, and inner regions (right). **f**, SIM images of localization of HNRNPF within PSs in HAP1 WT cells treated with MG132 (5 μM for 6 h). PS substructures were visualized by NEAT1 FISH with the mixture of NEAT1_5’ and NEAT1_3’ probes to detect the shell (green) and HNRNPF IF (magenta). Scale bar, 500 nm. Numerical data are available in the Source Data.

### Artificial tethering of SFPQ, HNRNPF, or BRG1 to the shells induces the segregation of mini-PS from NSs

We next examined whether the recruitment of these candidate proteins to mini-PS can compensate for the segregation defect. Since deleted regions (1–8 kb and 16.6–22.6 kb) in the mini-NEAT1 are localized in the shell of the PSs, we artificially tethered these candidate proteins fused with MCP onto mini-NEAT1 carrying 6 × MS2-binding sites (6 × MS2BS) at the 5’ shell region of the mini-NEAT1 (mini-NEAT1/6×MS2@8.2 kb)^23^ (Fig. 3a and Extended Data Fig. 4a). Strikingly, the shell tethering of SFPQ, HNRNPF, or BRG1 rescued the segregation defects of the mini-PS, and most of the mini-PS localized adjacent to NSs, although the other candidates, TDP-43 and HNRNPH1, did not (Fig. 3b–d and Extended Data Fig. 4a, b). Overexpression of these MCP-fused proteins in parental mini-NEAT1 cells did not rescue the segregation defects, suggesting that the interactions of SFPQ, HNRNPF, and BRG1 with NEAT1 are required for the complementation (Fig. 3b,c). Furthermore, we performed similar tethering to the core of mini-PS in the mini-NEAT1 cell line carrying 6 × MS2BS at the 3’ end (mini-NEAT1/6×MS2@16.2 kb), which localizes in the core of PSs (Fig. 3a,d and Extended Data Fig. 4a). The tethering of SFPQ, HNRNPF, or BRG1 to the core did not efficiently rescue the segregation defects, although these proteins were expressed and recruited to PSs in mini-NEAT1/6×MS2@16.2 kb at a level comparable to that in mini-NEAT1/6×MS2@8.2 kb (Fig. 3b,c and Extended Data Fig. 4c,d). These findings suggest that the tethering of SFPQ, HNRNPF, and BRG1 on the shell of the mini-PS facilitates the segregation of mini-PS from NSs.

**Fig. 4.**
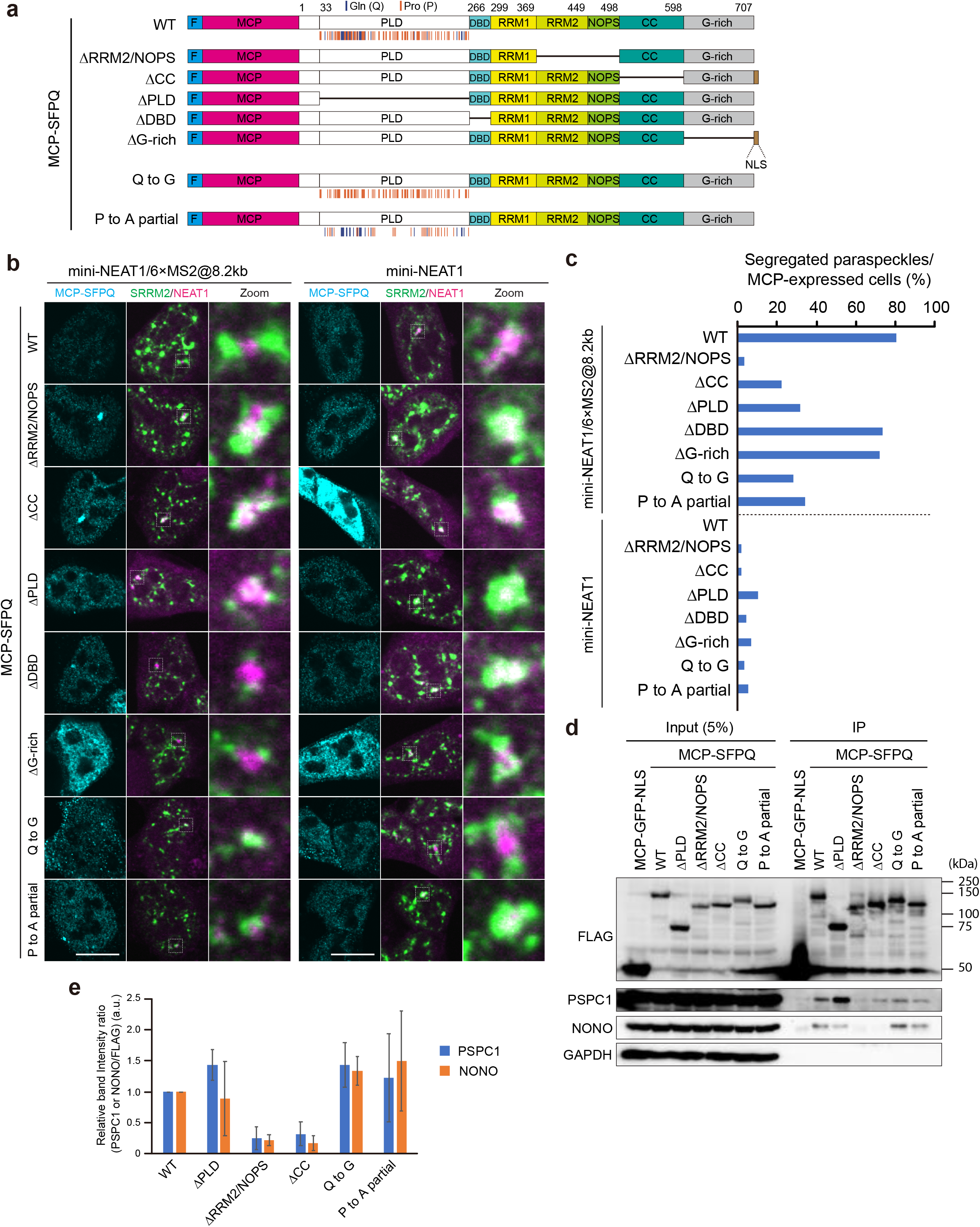
The dimerization, oligomerization, and prion-like domains of SFPQ are required for the mini-PS segregation from NSs. **a**, Schematics of MCP-fused SFPQ WT and mutants used for tethering experiments. The positions of glutamine and glycine residues in the PLD of SFPQ are indicated with blue and orange lines, respectively. **b**, Confocal observation of PSs and NSs with transfection of MCP-SFPQ WT and mutants into mini-NEAT1/6×MS2@8.2 kb (left) and mini-NEAT1 (right) in the MG132 treatment conditions (5 μM for 6 h). White boxes indicate the areas shown at a higher magnification. Scale bars, 10 μm. **c**, Proportion (%) of the cells with PSs segregated from NSs in **b**. Data were collected from three independent experiments. Mini-NEAT1/6×MS2@8.2 kb: *n* = 61 (WT), *n* = 62 (βRRM2/NOPS), *n* = 67 (βCC), *n* = 66 (βPLD), *n* = 76 (βDBD), *n* = 61 (βG-rich), *n* = 60 (Q to G), *n* = 64 (P to A partial); mini-NEAT1: *n* = 59 (WT), *n* = 63 (βRRM2/NOPS), *n* = 62 (βCC), *n* = 67 (βPLD), *n* = 66 (βDBD), *n* = 60 (βG-rich), *n* = 61 (Q to G), *n* = 58 (P to A partial). **d**, Coimmunoprecipitation (co-IP) of MCP-SFPQ WT and mutants. Immunoblotting was performed to detect DBHS proteins in the co-IP samples. GAPDH served as a negative control. **e**, Quantification of co-IP efficiency of the data shown in **d**. Values (PSPC1/FLAG or NONO/FLAG) are mean ± SD of three independent experiments. Data were compared using Kruskal-Wallis ANOVA and post hoc Dunn’s multiple comparison test. Kruskal-Wallis test showed statistically significant differences (PSPC1: *P* = 0.0345, NONO: *P* = 0.0489), but the post hoc Dunn’s multiple comparison test did not. Numerical data and unprocessed blots are available in the Source Data.

We next examined the localization of these proteins within the PSs. SFPQ was localized in the outer as well as the inner layers of the PSs (Fig. 3e). HNRNPF exclusively localized on the shell of the PS (Fig. 3f). A previous work showed that BRG1 shows patchy localization within the PS and some fractions of BRG1 localize in the shell of the PS^23, 27^. These findings support the idea that SFPQ, HNRNPF, and BRG1 in the shell of the PS contribute to the segregation of WT-PSs from NSs.

### The mechanism to rescue mini-PS segregation defect by SFPQ and HNRNPF differs from that by BRG1

To clarify the relationship of the SFPQ, HNRNPF, and BRG1 proteins, we examined the reciprocal recruitment to mini-PS by their tethering (Extended Data Fig. 4e–g). The tethering of HNRNPF facilitated SFPQ recruitment to mini-PS, which is similar to the tethering of NONO and PSPC1 likely through recruitment of SFPQ proteins by heterodimerization (Extended Data Fig. 4b,e,h). Meanwhile, the tethering of SFPQ did not facilitate HNRNPF recruitment (Extended Data Fig. 4f), suggesting a hierarchical relationship between SFPQ and HNRNPF in the recruitment. In addition, the tethering of either SFPQ or HNRNPF did not facilitate BRG1 recruitment (Extended Data Fig. 4g), although the tethering of BRG1 marginally facilitated HNRNPF recruitment and showed a trend toward increased SFPQ recruitment (Extended Data Fig. 4e,f). Collectively, these findings suggest that the mechanism by which mini-PS segregation defects are rescued by SFPQ or HNRNPF tethering appears to differ from that by BRG1, and that the tethering of HNRNPF likely rescues the segregation defect at least partly through SFPQ recruitment.

To gain insight into this recruitment mechanism, we examined how these proteins are recruited to PSs by NEAT1 RNA regions. We used NEAT1 Δ4–8 kb/16.6–22.6 kb and Δ1–4 kb/16.6–22.6 kb cell lines, which have additional NEAT1 regions compared to mini-NEAT1 and in which paraspeckles are segregated from nuclear speckles (Extended Data Fig. 2a,c,d). Recruitment of SFPQ and HNRNPF to PSs is restored in these cell lines compared with that in the mini-NEAT1 cell line, although the recruitment of BRG1 to PSs is not restored at all in these cell lines (Extended Data Fig. 4i,j). These findings are consistent with the above mechanism and suggest that the recruitment of SFPQ/HNRNPF and BRG1 is determined by different NEAT1 RNA domains.

### The dimerization, oligomerization, and prion-like domains of SFPQ are required for the segregation of the PSs from NSs

As SFPQ appears to be a key factor to understand this segregation process, we next investigated SFPQ domains promoting the segregation of mini-PS from NS. SFPQ contains multiple motifs, including RRM2/NOPS, which is required for dimerization with DBHS proteins; coiled-coil (CC), which is necessary for the oligomerization of DBHS proteins; putative DNA binding domain (DBD); and PLD (Fig. 4a and Extended Data Fig. 5a–d). The deletion of RRM2/NOPS (ϕλRRM2/NOPS) completely abolished the ability to rescue the segregation defects, and the deletion of CC or PLD significantly reduced this ability (Fig. 4b, c and Extended Data Fig. 5e). Meanwhile, MCP-SFPQ ϕλDBD and ϕλG-rich mutants rescued the defects similarly to the tethering of MCP-SFPQ WT (Fig. 4b, c). In these MS2 tethering experiments, the tethering might increase the local protein concentration to non-physiological levels in PSs. To test whether the recruitment of physiological levels of SFPQ to PSs is sufficient for the mini-PS segregation, we selected and analyzed the cells in which comparable amounts of SFPQ were localized to mini-PSs as in WT-PSs (Extended Data Fig. 5f). Even under these conditions, MCP-SFPQ WT rescued segregation defects, but MCP-SFPQ ΔRRM2/NOPS did not (Extended Data Fig. 5g,h). These results suggest that physiological concentrations of SFPQ are sufficient to induce PS segregation from NSs. To understand the molecular mechanism behind the rescue activities, we performed immunoprecipitation (IP) using these SFPQ mutants. As expected, SFPQ βRRM2/NOPS and ΔCC completely lost the interactions with SFPQ, NONO, and PSPC1^36, 37^ (Fig. 4d,e and Extended Data Fig. 5i,j) ^36, 37^. In contrast, SFPQ ΔPLD interacted with SFPQ, NONO, and PSPC1 (Fig. 4d,e and Extended Data Fig. 5i,j). These results suggest that SFPQ βRRM2/NOPS and βCC probably cannot rescue the segregation defects due to the lack of interactions with the DBHS proteins, whereas SFPQ ΔPLD likely lost the rescue activity through a different mechanism.

**Fig. 5.**
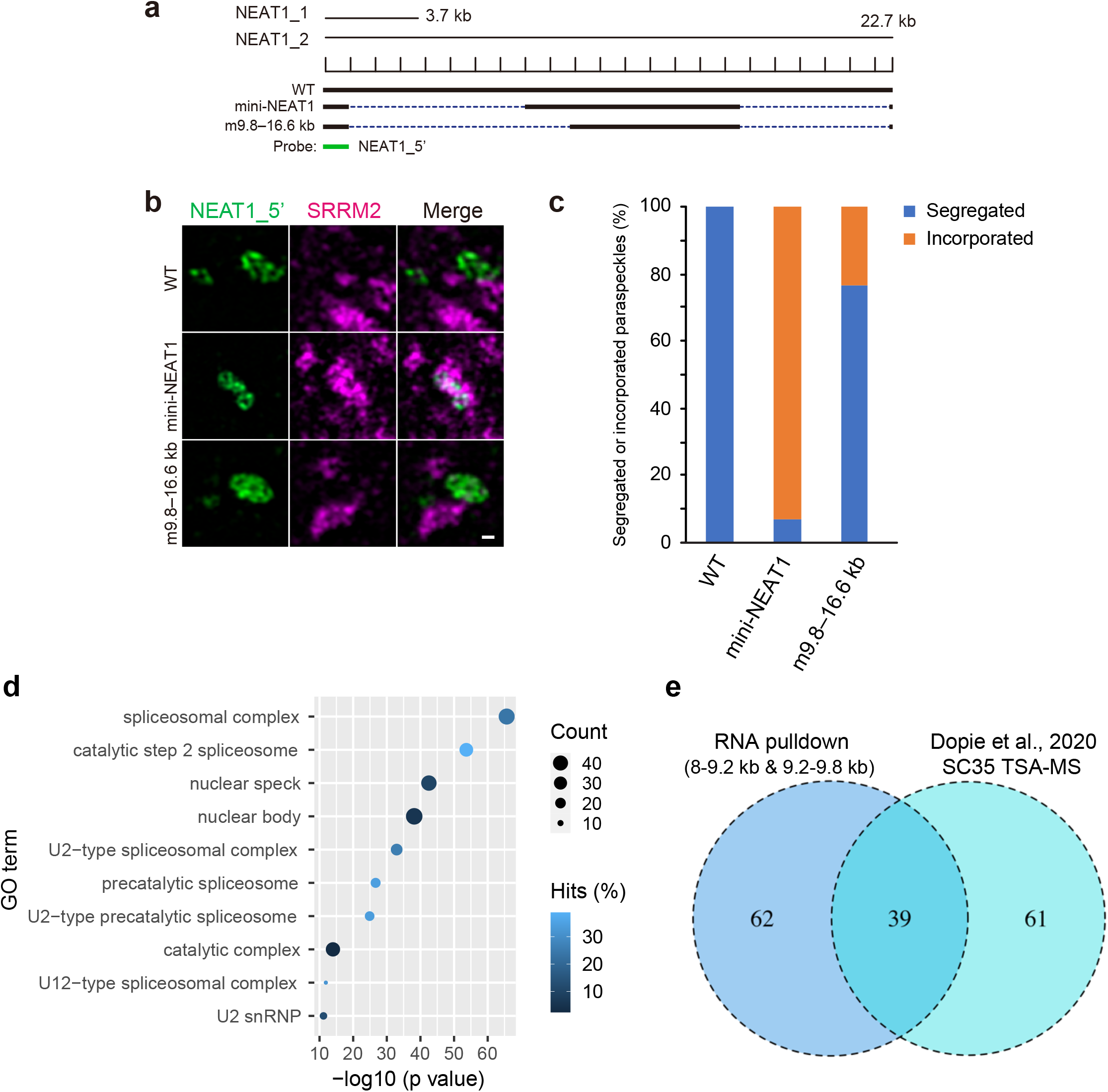
U2 snRNP-related proteins are major interacting proteins of the mini-NEAT1 RNA domain responsible for incorporation into NSs. **a**, Schematics of the WT, mini-NEAT1, and m9.8–16.6 kb mutants are shown with a scale. The position of a NEAT1_5’ probe used in SRM (green) is shown below. **b**, SRM images of PSs (RNA-FISH with NEAT1_5’ probe) and NSs (SRRM2 IF) in HAP1 WT and mini-NEAT1, and m9.8–16.6 kb mutant cells treated with MG132 (5 μM for 6 h). **c**, Proportion (%) of the cells with segregated PSs from NSs in **b**. Data were collected from two independent experiments. WT: *n* = 104, mini-NETA1: *n* = 100, m9.8–16.6kb: *n* = 110. **d**, Top 10 Gene Ontology cellular compartment enrichment for RNA pulldown and MS spectrometry analysis using hNEAT1 8–9.2 kb and 9.2–9.8 kb probes. **e**, Venn diagram showing the overlap between RNA pulldown (8–9.2 kb and 9.2–9.8 kb) and SC35 TSA-MS^38^. Numerical data are available in the Source Data.

SFPQ-PLD (33–265 aa) is required to rescue the segregation defects efficiently. Proline (P, 37.8% [88/233]), glutamine (Q, 10.7% [25/233]), glycine (G, 19.3% [45/233]), and histidine (H, 4.3% [10/233]) are highly enriched in the PLD (Fig. 4a and Extended Data Fig. 5a–d). To investigate their functional importance, we made two MCP-SFPQ mutants carrying Q to G or partial P to A mutations in the PLD (Fig. 4a). These Q to G or P to A partial mutants did not efficiently rescue the segregation defect of mini-PS similarly to SFPQ ΔPLD (Fig. 4b,c). These PLD mutants interacted with the DBHS proteins comparably to SFPQ WT (Fig. 4d,e and Extended Data Fig. 5i,j). Thus, glutamine and proline residues in SFPQ-PLD are required for the segregation process, and the interactions of SFPQ with DBHS proteins are not sufficient for the rescue activity.

We next investigated whether this rescue activity is related to the ability to induce PS assembly by SFPQ^23^. To examine the ability to perform assembly, we used an SFPQ tethering experiment in NEAT1 m13-16.6 kb mutant cells, in which PS assembly is defective^23^. Although most of the SFPQ mutants exhibited a compromised ability to achieve PS assembly, the SFPQ Q to G mutant, which cannot rescue the segregation defect, retained the PS assembly ability (Extended Data Fig. 5k,l). Thus, these two abilities are separable, and the ability to induce the assembly is not sufficient to rescue the segregation defect.

### U2 snRNP-related proteins are major interacting proteins of the mini-NEAT1 domain responsible for incorporation into NSs

The mini-PSs are incorporated into NSs, but not into the other nuclear bodies. Thus, we reasoned that certain factors determined this specific incorporation. To explore this mechanism, we first searched for RNA subdomains of mini-NEAT1 required to incorporate into NSs. We analyzed the m9.8–16.6 kb cell line used in our previous study (Fig. 5a)^23^. PSs in this cell line are segregated from NSs (Fig. 5b,c). In addition, the recruitment of SFPQ and BRG1 was not recovered (Extended Data Fig. 6a,b). These findings suggest that the RNA subdomain (8–9.8 kb region) of mini-NEAT1 actively promotes the internalization of mini-PS into NSs.

**Fig. 6.**
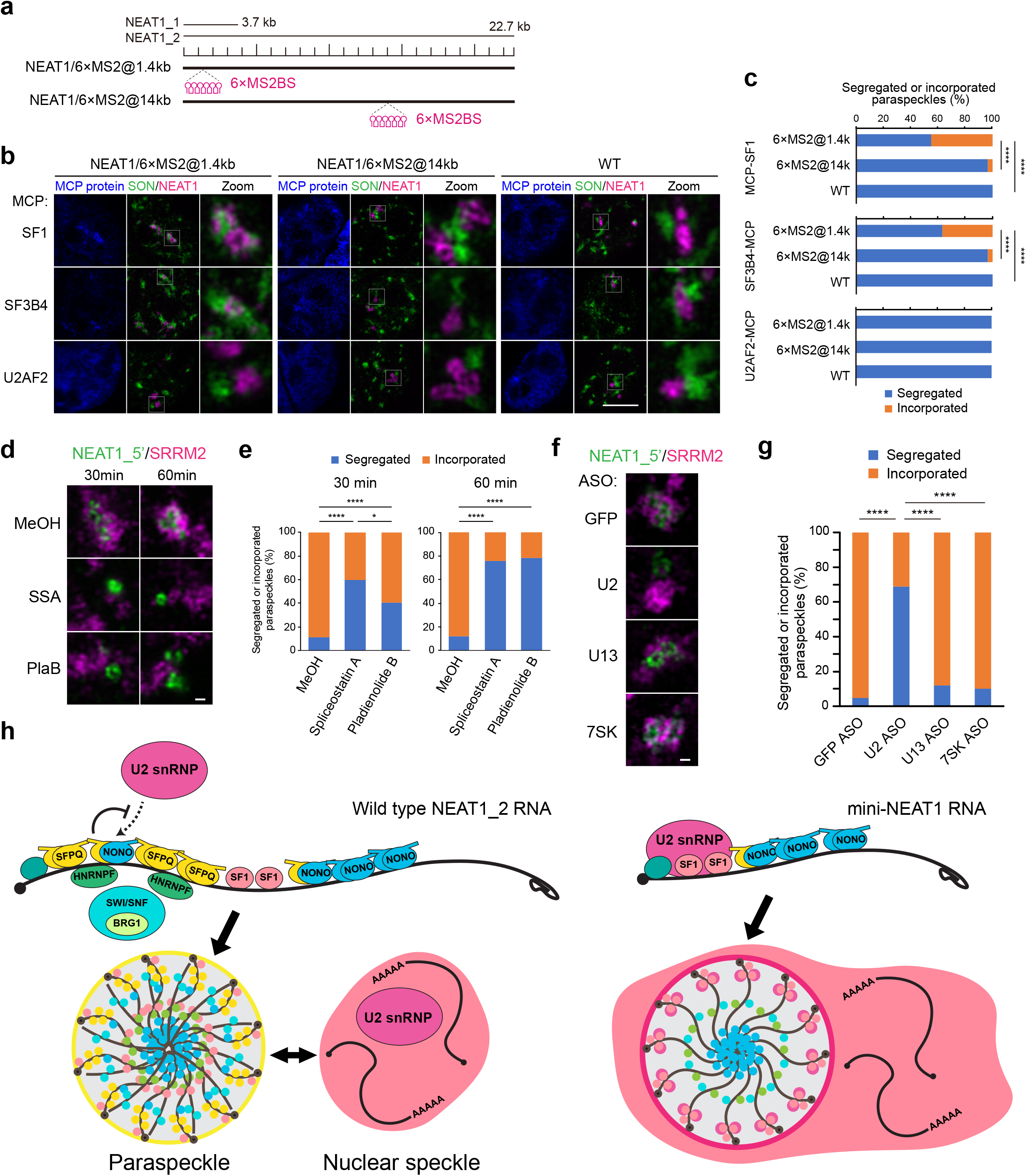
Artificial shell tethering of U2 snRNP components induces the internalization of PSs into NSs. **a**, Schematics of cell lines with 6 × MS2BS for MCP-protein tethering experiments to the shell (5’ or 3’) or core of PSs. **b**, SRM maximum projection images of PSs and NSs with transfection of MCP proteins into NEAT1/6×MS2@1.4 kb (left), NEAT1/6×MS2@14 kb (middle), and WT (right) in the MG132 treatment conditions (5 μM for 6 h). White boxes indicate the areas shown at a higher magnification. Scale bar, 5 μm. **c**, Proportion (%) of the cells with segregated PSs from NSs in **b**. Data were collected from three independent experiments. All samples (*n* = 60). Scale bar, 10 μm. **d**, SRM images of PSs and NSs in mini-NEAT1 cells treated with 100 ng/ml SSA or 1 μM PlaB for 30 or 60 min. Scale bar, 500 nm. **e**, Proportion (%) of the cells with PSs segregated from NSs in **d**. Data were collected from two independent experiments. 30 min: *n* = 208 (MeOH), *n* = 131 (SSA), *n* = 125 (PlaB). 60 min: *n* = 196 (MeOH), *n* = 115 (SSA), *n* = 133 (PlaB). **f**, SRM images of the PSs and NSs in mini-NEAT1 cells transfected with indicated ASOs in the MG132 treatment conditions (5 μM for 6 h). Scale bar, 500 nm. **g**, Ratio (%) of the cells with segregated PSs from NSs in **f**. Data were collected from at least two independent experiments. GFP ASO: *n* = 103, U2 ASO: *n* = 77, U13 ASO: *n* = 185, 7SK ASO: *n* = 109. **h**, Model diagram illustrating how proteins localized to the shell of a PS determine its independence in the nucleus. **c**,**e**,**g**, \*\*\*\**P* < 0.0001, \*\*\**P* < 0.001, \**P* < 0.05. Data were compared using Fisher’s exact test and Bonferroni correction. If the statistical test showed no significant difference (*P* > 0.05), it is not specified in the figure. Numerical data are available in the Source Data.

We then hypothesized that proteins interacting with the domain contribute to the internalization. To identify such proteins, we performed in vitro RNA pulldown using two NEAT1_2 RNA fragments covering the 8–9.8 kb region, followed by mass spectrometry analyses (MS). This analysis identified more than 100 proteins, including spliceosomal U2 snRNP proteins such as the SF3B complex components, SF1, as well as known PSPs as major constituents (Fig. 5d, Extended Data Fig. 6c, and Supplementary Table 1). We found a significant overlap between the NEAT1-associated proteins and NS components identified by the TSA-MS analysis^38^ (Fig. 5e). Similar overlap was also prominent between a recent in vivo identification of PSPs by the HyPro-MS and the NS components identified by the TSA-MS analysis^38, 39^ (Extended Data Fig. 6d,e). As overlapping protein interaction networks are proposed to be important for interactions between distinct MLOs^40^, these findings suggest the possibility that U2 snRNP-related proteins are involved in the interplay between PSs and NSs.

### Artificial shell tethering of U2 snRNP-related proteins induces the internalization of PSs into NSs

To examine whether the identified U2-related spliceosomal proteins are involved in the PS internalization, we tethered several proteins (SF1, SF3B4, SF3A3, U2AF2, PTBP1) identified by our MS in the NEAT1 WT cell lines carrying 6 × MS2BS (Fig. 6a). We tethered the proteins to the shell of the PSs in the NEAT1_2 WT cell line carrying 6 × MS2BS at the 5’ region (NEAT1/6×MS2@1.4 kb). The shell tethering of SF1, SF3B4, and SF3A3 induced internalization of the PSs, whereas U2AF2 and PTBP1 did not (Fig. 6b,c and Extended Data Fig. 7b–d). Furthermore, we tethered these proteins to the core of the PSs in the NEAT1_2 WT cell line carrying 6 × MS2BS at the NEAT1_2 middle region (NEAT1/6×MS2@14 kb). The core tethering of SF1, SF3B4, SF3A3, U2AF2, and PTBP1 did not induce internalization of the PSs (Fig. 6b,c and Extended Data Fig. 7b–d). Consistent with the importance of the U2 snRNP-related proteins in the PS shell, SF3B1, a major component of the SF3B complex, is localized in the PS shell, especially in mini-PS (Extended Data Fig. 7e), showing inverse correlations between the recruitment of SFPQ, HNRNPF, and BRG1 and the recruitment of U2 snRNP-related proteins to PSs. These findings suggest that U2 snRNP-related proteins residing in the PS shells induce the internalization of PSs.

Finally, we attempted to obtain insights into the involvement of the whole U2 snRNP complex in the internalization of mini-PS into NSs. We used chemical inhibitors for U2 snRNP assembly, spliceostatin A (SSA) and pladienolide B (PlaB)^41, 42^. Treatment with these drugs led to significant segregation of mini-PS from NSs in the mini-NEAT1 cell line (Fig. 6d,e). We also performed knockdown (KD) of U2 snRNA using antisense oligonucleotides (ASOs). Consistently, U2 KD resulted in segregation of the mini-PS from NSs, whereas several control ASOs (ASOs against GFP, U13 snoRNA, 7SK snRNA) did not (Fig. 6f,g and Extended Data Fig. 7f). These findings support the idea that the U2 snRNP complex is involved in the internalization of mini-PS into NSs.

## Discussion

We previously identified functional NEAT1 RNA domains involved in PS formation and function^23, 30, 31^. Here, we revealed new functional NEAT1 RNA domains (1–8 kb and 16.6–20.2 kb domains) indispensable for constructing PSs as independent MLOs. These two domains are localized in the paraspeckle shell^23, 31, 32^. These NEAT1 domains facilitate the recruitment of several specific PSPs (SFPQ, HNRNPF, and BRG1) required for PS segregation from NSs. Consistent with these findings, tethering of these proteins to the shell, not to the core, of PS micelles repositions mini-PS in close proximity to NSs. Thus, we could modulate the properties of specific regions within the mesoscopic scale MLO by tethering proteins to different parts of the arcRNA.

We identified SFPQ as a key protein contributing to the segregation process. We previously reported that SFPQ is essential for NEAT1_2 expression and PS assembly^22, 23^. From our mutational analyses, the PLD, dimerization, and oligomerization domains of SFPQ were shown to be required for both the segregation of mini-PS from NSs and PS assembly. Among the SFPQ mutants we tested, the tethering of SFPQ carrying Q to G mutations in the PLD cannot induce mini-PS segregation from NS, but it can induce PS assembly. This mutant can interact with NONO, an essential protein for PS assembly^23^. Thus, SFPQ has separable functions in the segregation and assembly of PSs. SFPQ is localized in both core and shell (Fig. 3e) and exerts its function in PS assembly likely through binding to PS cores^23^. The NEAT1_2 RNA domains for the segregation are located in the PS shell. Thus, shell-localizing SFPQ likely plays a role in the segregation of PSs from NSs. A bioinformatics study showed that HNRNPF binding sequences are enriched around the SFPQ binding sites and that HNRNPF plays a role in SFPQ recruitment to RNAs^43^, which is consistent with our data (Extended Data Fig. 4e,i,j). Further studies on this multifunctional SFPQ and its recruitment mechanism will contribute to the understanding of the molecular basis of the role of the PS shell.

In this study, we identified the proteins involved in the atypical PS incorporation into NSs (particularly U2 snRNP-related proteins) as well as the proteins involved in the ordinary PS segregation from NSs (Extended Data Fig. 8a). These findings provide insights into PS biogenesis as a distinct MLO. NEAT1_2 lncRNAs are transcribed by Pol II, as mRNAs are. In general, mRNAs are localized to NSs and exported from the nucleus to the cytoplasm for translation^44^. In contrast, NEAT1_2 lncRNA is retained in the nucleus, and the 3’ end has a triple helix RNA structure, not a poly(A) tail, for its stabilization^24, 45^. Thus, NEAT1_2 undergoes an exceptional pathway to achieve maturation by escaping from the canonical mRNA maturation pathway including splicing, polyadenylation, and nuclear export. Although NEAT1_2 is not spliced, the NEAT1_2 sequence contains multiple potential branch point sequences (Extended Data Fig. 7g). Thus, we speculate that NEAT1_2 as an architectural RNA dedicates itself to assembling PS by preventing the binding of factors related to mRNA maturation pathways. This mechanism may be important for the proper maturation of NEAT1_2 followed by PS assembly with a proper shell structure that maintains it as a distinct nuclear MLO. Moreover, previous studies suggested that arcRNAs are generally not spliced^13, 35, 46–49^. Roles of SFPQ, HNRNPF, and BRG1 in the inhibition of splicing have been reported^50–53^, which is consistent with our observation of the inverse correlation of SFPQ/HNRNPF/BRG1 recruitment and SF3B1 recruitment (Fig. 2a,b and Extended Data Fig. 7e,g). From these findings, we propose that the co-transcriptional formation of NEAT1_2 RNPs antagonizes the association of splicing factors (e.g., U2 snRNP) with NEAT1 lncRNAs to construct PSs as distinct nuclear MLOs (Fig. 6h).

Our study also provides insights into the importance of the core-shell architecture of the PS, which is one of the distinct characteristics of the PS constructed by micellization^31–33^. De novo-formed mini-PSs are incorporated into NSs while retaining their internal core-shell structure after forming outside of NSs (Fig. 1d,e,g,h,i). We show that artificial tethering of U2 snRNP-related proteins to the 5’ end of WT NEAT1_2 induced significant internalization of the PSs into NSs. Thus, the shell localization of the proteins identified in this study controls their NS localization or nucleoplasmic localization, although we cannot completely rule out the possibility that the NEAT1 5’ terminal domain independent of its shell localization contributes to the PS segregation. It has recently been reported that interactions of specific MLO components regulate the close proximity of the MLOs formed by LLPS (e.g., stress granule and P-body, Cajal body and Gem)^40, 54^. Because U2 snRNPs are recruited to nascent RNAs^55, 56^, it is conceivable that these U2 snRNP proteins are co-transcriptionally recruited to mini-NEAT1 lncRNAs and that the mini-NEAT1 forms mini-PSs containing U2 snRNPs (Extended Data Fig. 7e). Because U2 snRNP components interact with other NS proteins and RNAs, these protein-protein and/or protein-RNA interactions occurring at the interface between the mini-PS and the NS would contribute to the incorporation of mini-PSs into NSs. Furthermore, U2 snRNAs show some degree of peripheral localization in the NSs^54^. These reports and our observations raise the intriguing possibility that some degree of interactions of U2 snRNP-related proteins, which localize in the periphery of the NSs, determines the close proximity of the PS to the NSs. This notion is supported by the eCLIP data showing that U2 snRNP-related proteins preferentially interact with the 5’ terminal region of the NEAT1, which localizes to the shell of the PS (Extended Data Fig. 7e,g). Therefore, the mechanism identified in this study may capture the molecular link between PSs and NSs, contributing to the understanding of how and why these bodies localize in close proximity.

It remains to be determined how SFPQ, HNRNPF, and BRG1 contribute to the formation of WT-PSs as distinct MLOs, although other unidentified proteins might also be involved in this process^22, 57^. SFPQ, HNRNPF, and BRG1, which are mislocalized in NEAT1_2 mutants lacking shell-localizing RNA domains, exclusively or partly localize in the shell of the intact PS. Against this background, one important question is whether these proteins act directly and/or indirectly through factors interacting with them to control the composition of the PS shell. The composition of the shell is likely crucial for communication with other MLOs as well as for PS features/functions^34^. It is still unclear how MLOs scaffolded by RNAs, especially RNP block copolymer micelles, determine their composition, properties, and functions. Accordingly, our study provides new insights into the functional relationships between MLOs such as PSs and NSs and how arcRNAs encode the properties and functions of the MLOs they form.

## Materials and Methods

### Cell lines and cell culture

HAP1 cells were maintained in IMDM (Gibco) supplemented with 10% FBS (Sigma) and 1% penicillin-streptomycin (Nacalai Tesque). The cells were grown at 37°C in a humidified incubator with 5% CO_2_. Flp-In T-REx293 cell lines were maintained in DMEM containing high glucose and pyruvate (Nacalai Tesque) supplemented with 10% FBS (Sigma) and 1% penicillin-streptomycin (Nacalai Tesque). The cells were grown at 37°C in a humidified incubator with 5% CO_2_. To induce NEAT1 expression, the cells were treated with 5 μM MG132 (Sigma-Aldrich) for 6 h. For de novo PS formation experiments, cells were treated with 30 μM DRB (Sigma-Aldrich) for 4 h, and then the medium was replaced three times. To inhibit U2 snRNP assembly, cells were treated with 100 ng/ml spliceostatin A (kindly shared by Dr. Minoru Yoshida [RIKEN]) or 1 μM pladienolide B (Cayman Chemical) for 30 min or 1 h. The chemical compounds used in this study are listed in Supplementary Table 2.

### Genome editing using CRISPR/Cas9

CRISPR/Cas9-mediated deletions of the *NEAT1* gene were performed as previously described^23^. To delete the portions of NEAT1, PX330-B/B plasmids (2 μg) containing two sgRNA sequences were co-transfected with pcDNA6/TR plasmids (0.2 μg) containing the blasticidin resistance gene (Thermo Fisher Scientific) into HAP1 cells (1.5 × 10^6^ cells) by Nucleofector Kit V (Lonza) with a Nucleofector device (Lonza) using program “X-005,” in accordance with the manufacturer’s instructions. The sequences of sgRNA and HAP1 mutant cell lines used in this study are listed in Supplementary Table 2. PX330-B/B was used for deletion of the *NEAT1* gene. The sgRNAs were cloned into PX330-B/B vector^23^. For knock-in of 6 × MS2BS into the NEAT1 locus, three kinds of plasmids were transfected similarly using a Nucleofector device^58^: 1) PX330-B/B (2 μg) expressing two sgRNAs (a knock-in sgRNA to cut both sides of 6 × MS2BS or a knock-in vector to facilitate genomic integration, and an sgRNA targeting the NEAT1 locus), 2) a knock-in vector for 6 × MS2BS (2 μg), and 3) pcDNA6/TR plasmids (0.2 μg). The deletions and insertions were confirmed by genotyping PCR followed by Sanger sequencing analysis. The primers used in this study are listed in Supplementary Table 2.

### Generation of Flp-In T-REx 293 cell lines expressing MCP-SFPQ WT and mutants

To establish doxycycline-inducible Flp-In T-REx293 cells, 1.25 μg of each pcDNA5/FRT/TO/i/FLAG/MCP-based plasmid and 1.25 μg of pOG44 plasmid encoding recombinase were transfected using Lipofectamine LTX (Invitrogen). Cell lines with stably integrated constructs were selected with 150 μg/ml hygromycin (Wako). Transgene expression was induced by the addition of 1 μg/ml doxycycline for 48 h.

### Plasmid construction

To construct plasmids expressing N-terminal FLAG-MCP fusion proteins, TDP-43 and BRG1 ORF were inserted using the EcoRⅤ and XhoI sites of pcDNA5/FRT/TO/i/FLAG/MCP^23^; HNRNPF ORF was inserted using the BamHI and XhoI sites of pcDNA5/FRT/TO/i/FLAG/MCP; PSPC1 beta isoform ORF was inserted using the BamHI site of pcDNA5/FRT/TO/i/FLAG/MCP; and SF1, U2AF2, and PTBP1 ORF were inserted using the BamHⅠ and NotⅠ sites of pcDNA5/FRT/TO/i/FLAG/MCP. For the proteins that could not be recruited to PSs by the MS2 tethering system when MCP was fused to the N-terminal side, MCP was fused to the C-terminal side. To construct FLAG-HNRNPH1-MCP expression vector, HNRNPH1 ORF was inserted using the EcoRⅤ and XhoI sites of pcDNA5/FRT/TO/i/FLAG vector^59^, and MCP ORF was inserted using its XhoI site. To construct FLAG-SF3B4-MCP- and FLAG-SF3A3-MCP-expressing vectors, MCP ORF was inserted using the NotⅠ and XhoI sites of pcDNA5/FRT/TO/i/FLAG vector, and then SF3B4 and SF3A3 ORF were inserted using the BamHⅠ and NotⅠ sites of pcDNA5/FRT/TO/i/FLAG/C-terminal MCP vector. Since FLAG-SF3B4-MCP and FLAG-SF3A3-MCP were mainly localized in the cytoplasm, the SV40NLS sequence was inserted into the C-terminal side of it by site-directed mutagenesis. To construct plasmids expressing FLAG-MCP-SFPQ mutants including SFPQϕλRRM2/NOPS (ϕλ370-498aa), SFPQϕλCC (ϕλ499-598aa fused with SV40-NLS), SFPQϕλPLD (ϕλ33-265aa), SFPQϕλDBD (Δ266-298aa), and SFPQϕλG-rich (ϕλ599-707 fused with SV40 NLS), the deletions were introduced by site-directed mutagenesis. To construct plasmids expressing FLAG-MCP-SFPQ PLD mutants (Q to G and P to A partial), we used a custom cloning and synthesis service (AZENTA). The amino acid sequences of SFPQ PLD mutants are listed in Supplementary Table 2. To construct an HA-SFPQ-expressing vector, 3 × HA ORF was inserted using the HindⅢ site of pcDNA5/FRT/TO/i vector, and SFPQ ORF was inserted using its BamHⅠ and XhoI sites. The plasmids used in this study are listed in Supplementary Table 2.

### Plasmid transfection

For MS2 tethering experiments, HAP1 cells (5.0 × 10^4^ cells/well) were cultured on coverslips (Matsunami, micro-cover glass, 18 mm round; thickness, 0.16–0.19 mm) in a 12-well plate overnight and plasmids (1.5 μg) were transfected using 3 μl of TransIT-LT1 (Mirus). For IP experiments, HAP1 cells (9.0 × 10^5^ cells/plate) were cultured in a 100 mm plate overnight and plasmids (15 μg) were transfected using 45 μl of TransIT-LT1 (Mirus). The cells were cultured for 48 h, including the last 6 h of MG132 treatment. Then, the cells were subjected to each experiment.

### RNA-FISH and immunofluorescence

RNA-FISH and immunofluorescence were performed as previously described^23^. Confocal microscopy and structured illumination microscopy (referred to as SIM) analyses were performed as previously described^23^. Super-resolution microscopy analyses were performed using LSM900 with Airyscan2 with a 63× objective lens (referred to as SRM). The NEAT1 FISH probes for RNA-FISH analyses are listed in Supplementary Table 2.

For single-molecule FISH (smFISH) analyses, the cells were grown on coverslips (Matsunami; micro-cover glass; 18 mm round; thickness, 0.16–0.19 mm) and fixed with 4% paraformaldehyde/PBS at room temperature for 10 min. The cells were then washed with PBS, permeabilized with 0.5% Triton X-100/PBS for 5 min, and washed three times with PBS. Then, the coverslips were incubated with 1× blocking solution (Blocking Reagent [Roche] and TBST [TBS and 0.1% Tween 20]) at room temperature for 1 h. The coverslips were next incubated with primary antibodies in 1× blocking solution at 4℃ overnight, washed three times with TBST for 5 min, incubated with secondary antibodies in 1× blocking solution at room temperature for 1 h, washed two times with TBST for 5 min, and washed with PBS for 5 min. After removing the PBS, smFISH wash buffer (2× SSC and 10% formamide) was added and coverslips were incubated for 5 min. Then, 50 μl of hybridization solution (2× SSC, 100 mg/ml dextran sulfate, and 10% formamide) containing Stellaris NEAT1 probes was dropped onto the coverslips in a humidified chamber and incubated for 16 h in the dark. Next, the coverslips were washed with prewarmed smFISH wash buffer at 37℃ for 30 min, washed with 2× SSC at RT for 5 min, and washed with PBS for 5 min. The coverslips were mounted with VECTASHIELD Hard Set Mounting Medium with DAPI (Vector) or ProLong Gold Antifade reagent (Thermo Fisher Scientific). The antibodies and probes used are listed in Supplementary Table 2. We used SC35 antibody (Sigma) as a marker for NSs, but it mainly recognizes SRRM2 proteins^60^. For this reason, it is referred to as SRRM2 in the text.

All images of this study were processed for publication using ImageJ (NIH) and figures were prepared using Adobe illustrator 2019. ImageJ software (NIH) was used for quantifying the fluorescence intensity of RNA-FISH and IF signals with an intensity the area of PSs. The ‘Plot profile’ module was used for line profiles. ZEN software (Zeiss) was used for the construction of maximum-intensity projection images, 3D images, and movies. For the quantification of segregated or incorporated PSs in Figs. 1c,i, 3c, 4c, 5c, 6c,e,g and Extended data Figs. 1c, 2d, 5h, 7c, we manually classified PSs as segregated or incorporated. We show several typical examples of segregated or incorporated PSs in Extended Data Fig. 1b.

### Reverse-transcription quantitative PCR (RT-qPCR)

Total RNAs were purified with TRI reagent (Molecular Research Center), as previously described^35^. Briefly, homogenates of cells with TRI reagent were heated at 55°C for 20 min with 1000 rpm agitation. Total RNAs were purified in accordance with the manufacturer’s instructions. The total RNAs (0.5 μg) were reverse-transcribed using a High Capacity cDNA Reverse Transcription Kit (Thermo Fisher Scientific) with random hexamer primers. Aliquots of cDNA were amplified by qPCR using KAPA SYBR Fast qPCR Kit (NIPPON Genetics), in accordance with the manufacturer’s instructions. The primers used in this study are listed below. For NEAT1_2 detection, the primer set (forward primer: 5’-CAGTTAGTTTATCAGTTCTCCCATCCA-3’, reverse primer: 5’-GTTGTTGTCGTCACCTTTCAACTCT-3’) was used. For U2 snRNA detection, the primer set (forward primer: 5’-GCCTTTTGGCTAAGATCAAGTGTAGT-3’, reverse primer: 5’-CTATTCCATCTCCCTGCTCCAAA-3’) was used. For U13 snoRNA detection, the primer set (forward primer: 5’-AGTTCATGAGCGTGATGATTGG-3’, reverse primer: 5’-TGTGCCCACGTCGTAACAAG-3’) was used. For 7SK snRNA detection, the primer set (forward primer: 5’-CGGTCTTCGGTCAAGGGTAT-3’, reverse primer: 5’-TGGAGTCTTGGAAGCTTGAC-3’) was used. 18S (forward primer: 5’-TTTAAGTTTCAGCTTTGCAACCATACT-3’, reverse primer: 5’-ATTAACAAGAACGAAAGTCGGAGGT-3’) or GAPDH (forward primer: 5’-ATGAGAAGTATGACAACAGCCTCAAGAT-3’ reverse primer: 5’-ATGAGTCCTTCCACGATACCAAAGTT-3’) primer sets were used as loading controls.

### Immunoblotting

Cells were lysed in immunoprecipitation (IP) buffer (1× PBS, 0.1% Triton X-100, and cOmplete EDTA-free protease inhibitor cocktail [Roche]) and then disrupted by three pulses of sonication for 5 s. The cell lysates were cleared by centrifugation and the protein concentration was determined using the Bradford method. After adding SDS sample buffer, the samples were heated at 95°C for 5 min and separated by SDS-PAGE. After electrophoresis, the proteins were transferred to FluoroTrans W membrane (PALL) by electroblotting. The antibodies used are listed in Supplementary Table 2.

### Electron microscopy

Immunogold electron microscopy was performed on thin sections of cells embedded in Lowicryl K4M (Polysciences Inc.). HAP1 WT or mini-NEAT1 cells fixed in situ for 1 h at 4°C with 4% formaldehyde in 0.1 M Sörensen phosphate buffer pH 7.3 were scraped off and centrifuged. After rinsing in phosphate buffer, cell pellets were equilibrated in 30% methanol and deposited in a Leica EM AFS2/FSP automatic reagent handling apparatus (Leica Microsystems). Lowicryl polymerization under UV was performed for 48 h at – 20°C, followed by 40 h at 20°C. Ultra-thin sections were incubated at room temperature for 1 h with the primary antibody (e.g., mouse anti-NONO [BD Biosciences-N88520], diluted 1/20 in PBS; mouse anti-SFPQ antibody [Sigma-P2260], diluted 1/50 in PBS) and for 30 min with the secondary anti-mouse antibody coupled to 10 nm gold particles (BBI International, Cardiff, UK). To better visualize gold particles on electron-dense structures such as paraspeckles, thin sections were briefly contrasted with uranyl acetate, but lead citrate staining was omitted. Thin sections were analyzed with a Tecnai Spirit (FEI/Thermofisher) and digital images were taken with an SIS MegaviewIII charge-coupled device camera (Olympus). To analyze the distribution of gold particles within PS, three layers were delineated by progressive down-scaling to 81.2% and 57.4% of the 100% external contour. This generated three regions from the periphery to the interior with equal surface areas. Gold particles were counted visually and expressed as the percentage of gold particles in each of the three regions.

### Immunoprecipitation

Flp-In T-REx293 cells were treated with 1 μg/ml doxycycline for 48 h. The nuclear extracts (NEs) of Flp-In T-REx293 cells were prepared as previously described^61^. The NEs were frozen in liquid nitrogen and stored at −80°C until use. The NEs were cleared by centrifugation and the protein concentration was determined by the Bradford method. The NEs were diluted with IP buffer (1× PBS, 0.1% Triton X-100, 0.6 mM PMSF, and cOmplete EDTA-free protease inhibitor cocktail [Roche]) (protein concentration: ∼2.5 mg/ml), mixed with anti-FLAG M2 magnetic beads (Sigma-Aldrich), and rotated at 4°C overnight. The beads were washed five times with wash buffer (1× PBS, 0.1% Triton X-100, 0.6 mM PMSF). The IP samples were recovered by adding SDS sample buffer.

For analysis of the interaction between SFPQ WT and SFPQ mutant proteins, HAP1 cells (mini-NEAT1/6×MS2@8.2kb) transfected with pcDNA5/FRT/TO/i/FLAG/MCP-SFPQ WT or mutants and pcDNA5/FRT/TO/i/HA/SFPQ were cultured for 48 h, including the last 6 h with MG132 treatment, washed with cold PBS, and harvested using a cell scraper. The collected cells were lysed in IP buffer by five pulses of sonication for 5 s. The cell lysates were cleared by centrifugation and the protein concentration was determined using the Bradford method. The lysates were diluted with IP buffer (protein concentration: ∼1 mg/ml) and treated with RNase A (1 μg/ml) at 4°C for 1 h. Then, the lysates were mixed with anti-FLAG M2 magnetic beads (Sigma-Aldrich) and rotated at 4°C overnight. The beads were washed five times with wash buffer. The IP samples were recovered by adding SDS sample buffer.

### RNA pulldown

RNA pulldown was performed as previously described^23^. Biotinylated RNAs were synthesized with T7 or SP6 RNA polymerase (Roche) and Biotin RNA Labeling Mix (Roche), template DNAs were degraded by DNase I (Thermo Fisher Scientific) treatment, and the biotinylated RNAs were purified using a gel filtration column (CENTRI SEP Spin Column; Princeton Separations). The NEAT1 biotinylated RNA probes for RNA pulldown used in this study are listed in Supplementary Table 2. The HeLa NE (25 μl) (CILBIOTECH) spun at 20,000 × g for 5 min was mixed with 75 μl of RNA pulldown buffer (1× PBS, 0.1% Triton X-100, 0.6 mM PMSF, and complete EDTA-free protease inhibitor cocktail [Roche]) (final protein concentration: ∼2 mg/ml). The NE (100 μl) was precleared by mixing 10 μl of Tamavidin2-REV Magnetic Beads (WAKO), washed with RNA pulldown buffer, and then rotated at 4°C for 1 h. In vitro transcribed RNAs (0.5 μg in 10 μl of UltraPure Distilled Water [Thermo Scientific]) were heated at 90°C for 2 min and then cooled on ice for 2 min. An equal volume (10 μl) of 2× RNA structure buffer (20 mM Tris-HCl pH 7.4, 0.2 M KCl, and 20 mM MgCl_2_) was added and kept at room temperature for 20 min to allow proper secondary structure formation. The folded RNAs were mixed with washed Tamavidin2-REV Magnetic Beads (10 μl) and rotated at 4°C for 1 h. The unbound RNAs were washed with cold RNA pulldown buffer and then the RNA-bound Tamavidin2-REV Magnetic Beads (10 μl) were mixed with the precleared NE (100 μl). They were rotated at 4°C for 3 h and washed five times with cold wash buffer (1× PBS, 0.1% Triton X-100, and 0.6 mM PMSF), after which the bound proteins were eluted at 95°C for 5 min in SDS sample buffer. The eluates were subjected to SDS-PAGE, followed by immunoblotting.

### Mass spectrometry and Gene Ontology (GO) analyses

Mass spectrometric analyses with duplicated RNA pulldown samples were performed as described previously^62^. The number of unique peptides in the negative control was subtracted from the number of unique peptides obtained from each RNA pulldown sample. Keratin and likely contaminants (e.g., actin, tubulin, and myosin) are excluded. For GO analyses in Extended Data Fig. 6c, proteins with more than four peptides are used in the analysis, as shown in Supplementary Table 1. For GO analyses in Fig. 5d and e, proteins with a total number of peptides obtained from RNA pulldown using two different RNA probes (8–9.2 kb and 9.2–9.8 kb) of more than nine are used in the analysis, as shown in Supplementary Table 1. Gene Ontology cellular component (GO:CC) analysis was performed using g:Profiler (https://biit.cs.ut.ee/gprofiler/gost). The significance threshold was set to 0.05 and the term size was set to 4–2000.

### ASO-mediated knockdown

The ASO-mediated knockdown experiment was performed as previously described^63^. To deplete U2 snRNA, control GFP ASO (5’-mU*mC*mA*mC*mC*T*T*C*A*C*C*C*T*C*T*mC*mC*mA*mC*mU*-3’), U2 snRNA ASO (5’-mA*mG*mA*mA*mC*A*G*A*T*A*C*T*A*C*A*mC*mU*mU*mG*mA*-3’), U13 snoRNA ASO (5’-mC*mG*mU*mC*mG*T*A*A*C*A*A*G*G*T*T*mC*mA*mA*mG*mG*-3’), or 7SK snRNA ASO (5’-mC*mU*mU*mG*mA*G*A*G*C*U*U*G*U*U*U*mG*mG*mA*mG*mG*-3’) (asterisks represent the phosphothioate-modified backbone, and mN designates the 2’-O-methylribonucleotides) was transfected into HAP1 cells (1.0 × 10^6^ cells) by Nucleofector Kit V (Lonza) with a Nucleofector device (Lonza) using the program “X-005,” in accordance with the manufacturer’s instructions. After electroporation, cells were cultured for 8 h, including the last 6 h with MG132 treatment in 60 mm plates.

### Statistics and reproducibility

All experiments were performed at least twice independently, with similar results obtained. No statistical method was used to predetermine the sample size. No data were excluded from the analyses. Prism7 software (GraphPad) and R (version 4.2.2) were used for the statistical analyses. Kruskal-Wallis test with Dunn’s multiple comparison test was used for Fig. 4e and Extended Data Figs. 4d–g,j, 5g,j,l, 6b, and 7e. Mann-Whitney U-test (two-tailed) was used for Fig. 2b and Extended Data Fig. 3c. One-way ANOVA and post hoc Tukey’s multiple comparison test were used for Extended Data Fig. 7f. Fisher’s exact test and Bonferroni correction were used for Fig. 6c,e,g and Extended Data Figs. 2d, 7c. If the statistical test showed no significant difference (*P* > 0.05), it is not specified in the figure.

## Acknowledgments

The authors thank M. Okamura, C. Fujikawa, and A. Kubota for technical support and A. Marshall, A.H. Fox, C.S. Bond, B. Dyakov, A.C. Gingras, T. Yoda, M. Murakami, and members of the Hirose laboratory for valuable discussions. This research was supported by grants from the Ministry of Education, Culture, Sports, Science, and Technology (MEXT) of Japan (to T. Yamazaki [19K06479, 19H5250, 21H00253, 22H02545], to T.H. [20H00448, 20H05377, 21H05276, 22K19293]), the Mochida Memorial Foundation for Medical and Pharmaceutical Research (to T. Yamazaki), the Naito Foundation (to T. Yamazaki), the Takeda Science Foundation (to T. Yamazaki), JST CREST (JPMJCR20E6) to T.H., and an AMED grant (21479280) to T.H.

## Author contributions

H.T., T. Yamazaki, and T.H. conceived and designed this study. H.T. and T. Yamazaki conducted most of the experiments. S.S. and G.P. performed EM analyses. S.A. and T.N. performed mass spectrometry analyses. N.F. contributed to data analysis and manuscript editing. T. Yamamoto contributed to theoretical interpretations of the experimental data. H.Y. and S.N. contributed to the SRM analyses. H.T., T. Yamazaki, and T.H. wrote the manuscript.

---

**Extended Data Fig. 1.**
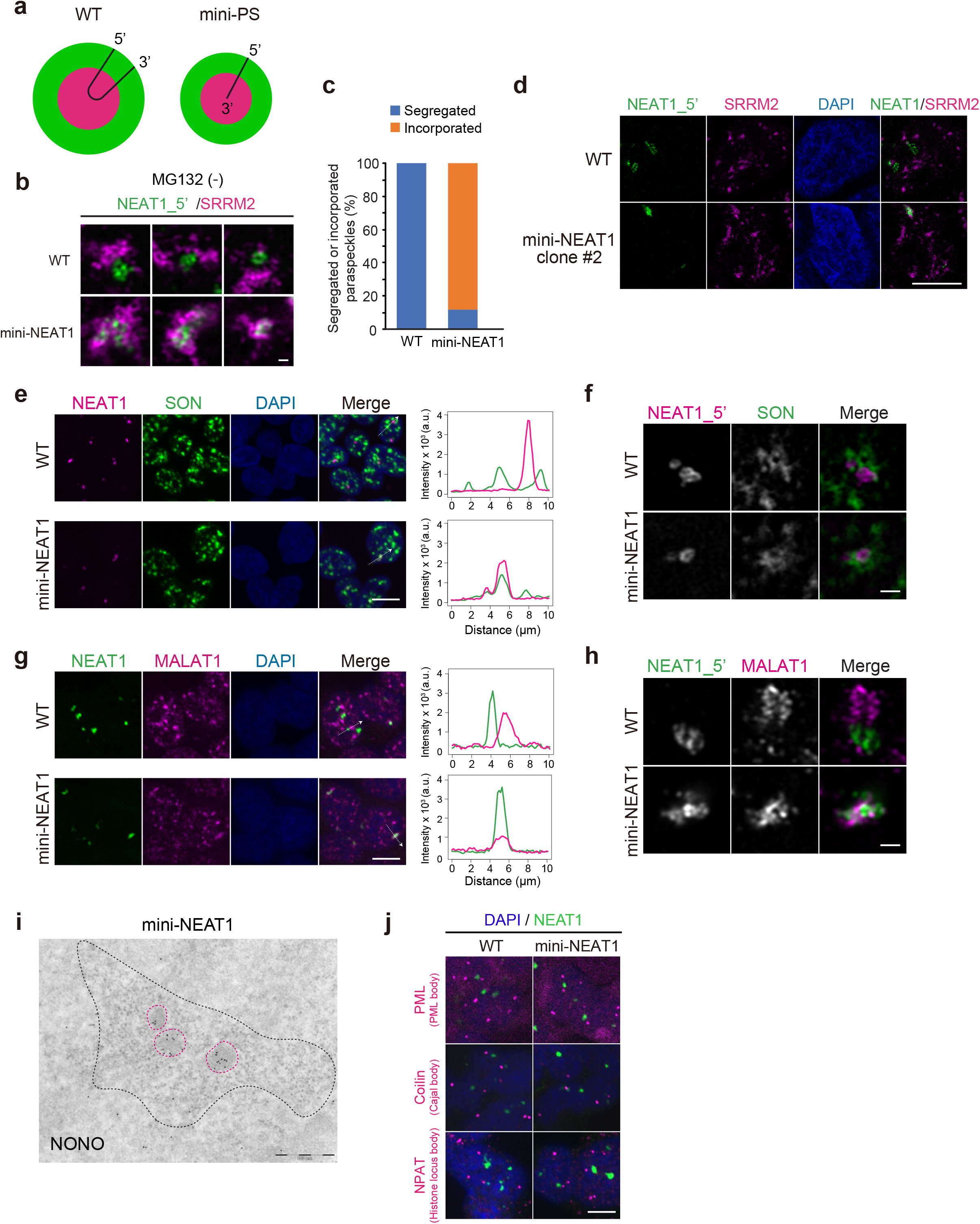
Mini-PSs are present within NSs, although NEAT1 transcription occurs outside of NSs. **a**, Schematics of the NEAT1_2 configuration in the WT-PS and mini-PS. **b**, SRM images of PSs (RNA-FISH with NEAT1_5’ probe) and NSs (SRRM2 IF) in HAP1 WT and mini-NEAT1 mutant cells in the absence of MG132. Scale bar, 500 nm. **c**, Graph showing the proportion of segregated or incorporated PSs in **b**. Data were collected from two independent experiments. WT: *n* = 167, mini-NEAT1: *n* = 210. **d**, SRM images of the PSs (RNA-FISH with NEAT1_5’ probe) and NSs (SRRM2 IF) in HAP1 WT and another mini-NEAT1 mutant cell clone treated with MG132 (5 μM for 6 h). Scale bar, 10 μm. **e**, PSs and NSs were detected by smFISH with NEAT1_5’ probes (magenta) and SON IF (green) in the presence of MG132 (5 μM for 6 h). Nuclei were stained with DAPI. Line profiles of the PSs and NSs are shown (right). **f**, SRM images of PSs (smFISH with NEAT1_5’ probe) and NSs (SON IF) in HAP1 WT and mini-NEAT1 cells treated with MG132 (5 μM for 6 h). Scale bar, 500 nm. **g**, PSs and NSs in HAP1 WT and mini-NEAT1 cells were detected by RNA-FISH with NEAT1_5’ probe and MALAT1 probe (green and magenta, respectively) in the presence of MG132 (5 μM for 6 h). Nuclei were stained with DAPI. Line profiles of the PSs and NSs are shown (right). Scale bar, 10 μm. **h**, SRM images of PSs (RNA-FISH with NEAT1_5’ probe) and NSs (RNA FISH with MALAT1 probe) in HAP1 WT and mini-NEAT1 cells treated with MG132 (5 μM for 6 h). Scale bar, 500 nm. **i**, EM observation of multiple, but unconnected, mini-PSs within an NS. The localization of NONO was detected as gold particles. Dotted magenta circles indicate the position of the mini-PSs. Dotted black circles indicate the position of the NS. Scale bar, 500 nm. **j**, Confocal observation of PSs and other nuclear bodies in HAP1 WT and mini-NEAT1 cells treated with MG132 (5 μM for 6 h). PSs were visualized by NEAT1 RNA-FISH with NEAT1_5’ probe (green). Nuclear bodies were visualized by IF of nuclear body marker proteins (magenta). The names of the nuclear bodies are shown in parentheses. Nuclei were stained with DAPI. Scale bar, 10 μm. Numerical data are available in the Source Data.

**Extended Data Fig. 2.**
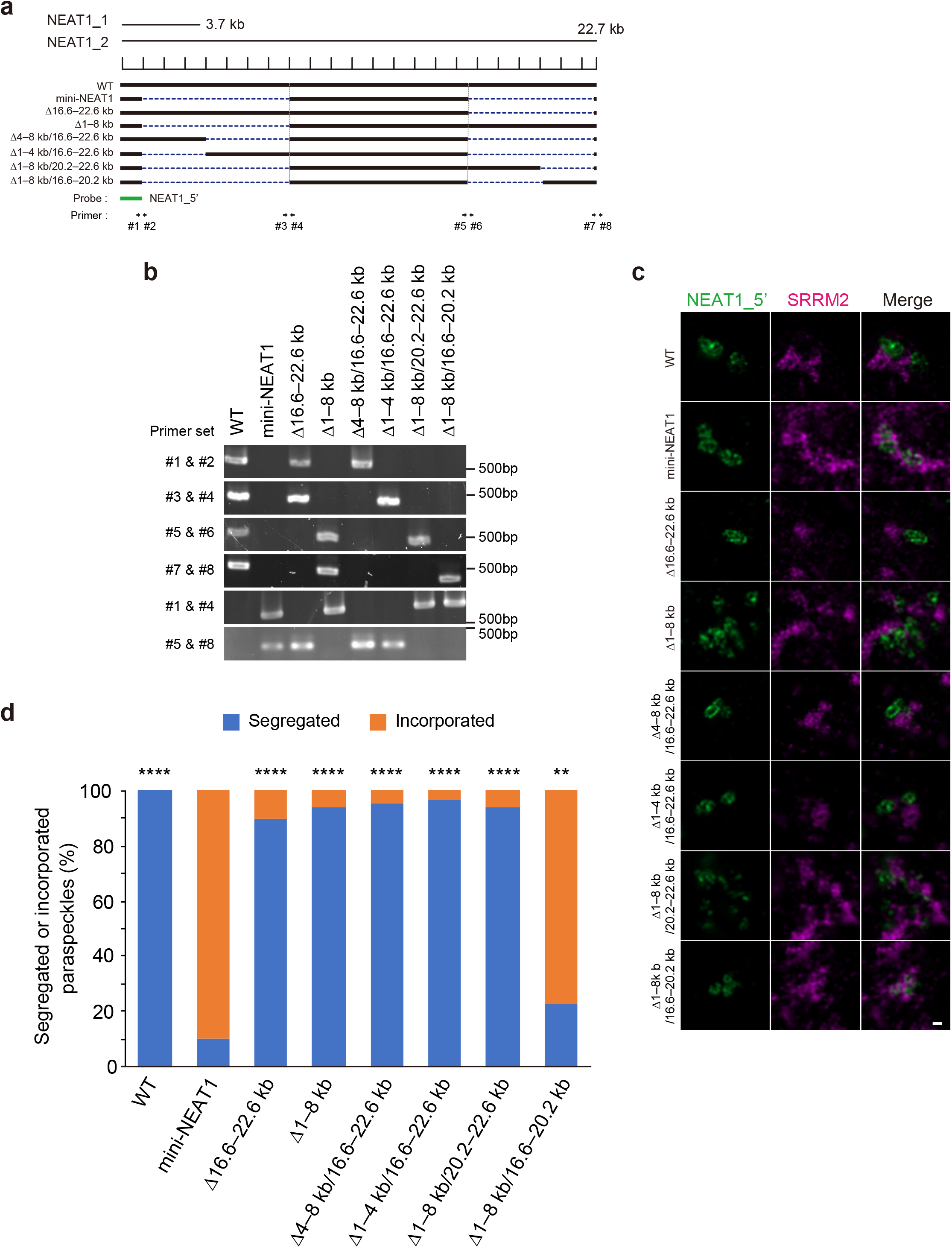
Identification of NEAT1_2 RNA domains required for PS segregation from NSs. **a**, Schematics of the WT, mini-NEAT1, and six NEAT1 deletion mutants are shown with a scale. The position of a NEAT1_5’ probe used in SRM (green) is shown below. **b**, Genotyping PCR to check for the deletions of the *NEAT1* locus. Gel electrophoresis of the PCR products confirmed the specific deletion of the *NEAT1* locus. The primers used for PCR are indicated with arrows in **a** and listed in Supplementary Table 2. **c**, SRM images of the PSs (RNA-FISH with NEAT1_5’ probe) and NSs (SRRM2 IF) in HAP1 WT, mini-NEAT1, and the six NEAT1_2 deletion mutant cells treated with MG132 (5 μM for 6 h). **d**, Graph showing the proportion of segregated or incorporated PSs in the cells observed in **c**. Data were collected from two independent experiments (WT: *n* = 183, mini-NEAT1: *n* = 198, τι16.6–22.6 kb: *n* = 188, τι1–8 kb: *n* = 196, τι4–8 kb/16.6–22.6 kb: *n* = 173, τι1–4 kb/16.6–22.6 kb: *n* = 186, τι1–8 kb/20.2–22.6 kb: *n* = 178, τι1–8 kb/16.6–20.2 kb: *n* = 194). \*\*\*\**P* < 0.0001, \*\**P* < 0.01. Data were compared to the mini-NEAT1 mutant using Fisher’s exact test and Bonferroni correction. Numerical data and unprocessed gels are available in the Source Data.

**Extended Data Fig. 3.**
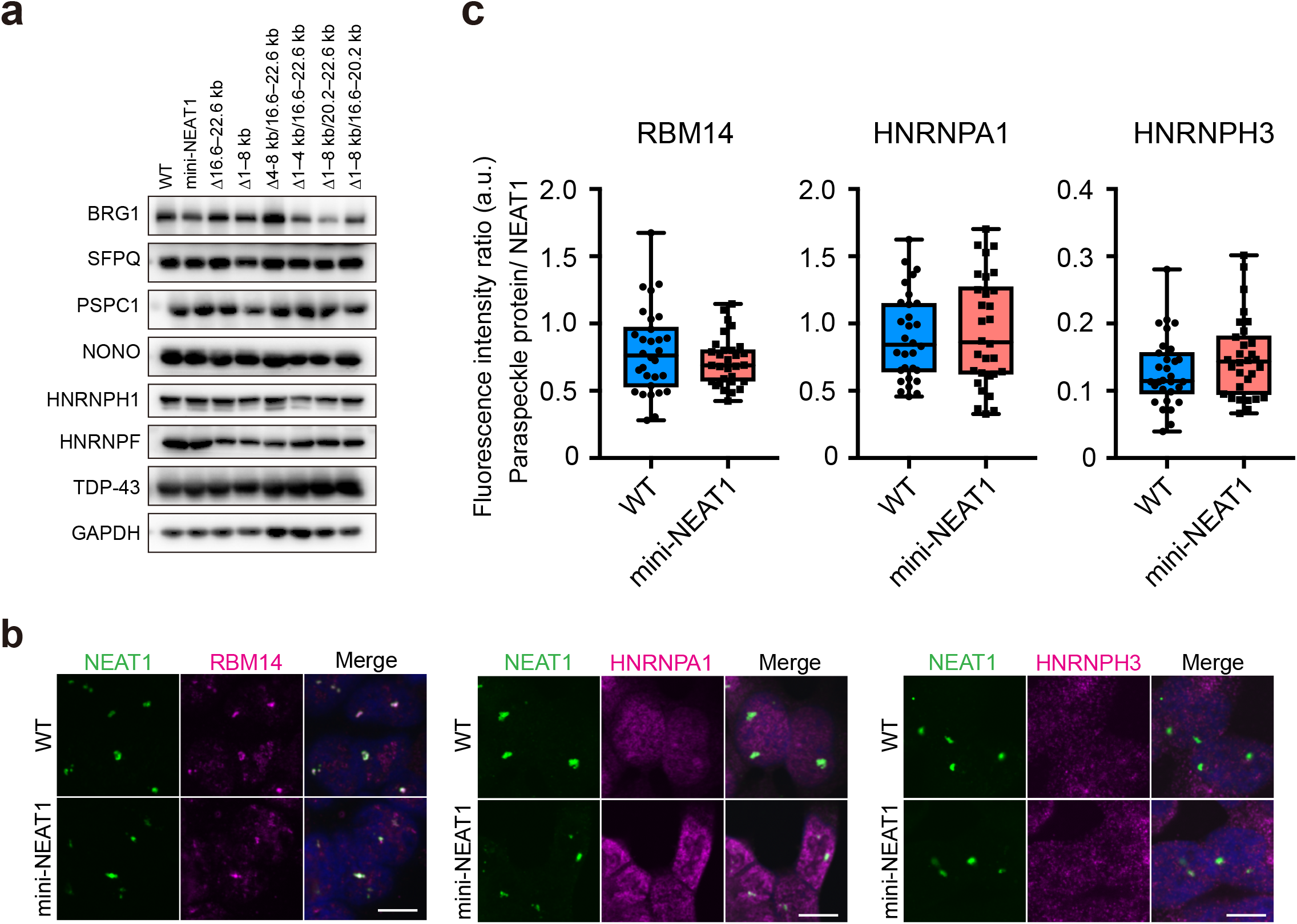
A subset of PSPs are similarly localized in WT-PSs and mini-PSs. **a**, Immunoblotting to examine the expression levels of various PSPs (BRG1, SFPQ, PSPC1, NONO, HNRNPH1, HNRNPF, and TDP-43) in HAP1 WT and the NEAT1 deletion mutant cells in Extended Data Fig. 2a. Cells were treated with MG132 (5 μM for 6 h). GAPDH was used as a control. **b**, Confocal observation of the PSs and various PSPs in HAP1 WT and mini-NEAT1 mutant cells treated with MG132 (5 μM for 6 h). PSs and PSPs were visualized by RNA-FISH using NEAT1_5’ probes (green) and IF (magenta). Nuclei were stained with DAPI. Scale bar, 10 μm. **c**, Quantification of fluorescence intensity ratio (PSPs/NEAT1) in the HAP1 WT (blue) and mini-NEAT1 (pink) cells as observed in **b** (all samples: *n* = 30). Each box plot shows the median (inside line), 25^th^-75^th^ percentiles (box bottom to top), and minimum-maximum values (whisker bottom to top). Data were compared using Mann-Whitney U-test. If the statistical test showed no significant difference (*P* > 0.05), it is not specified in the figure. Numerical data and unprocessed blots are available in the Source Data.

**Extended Data Fig. 4.**
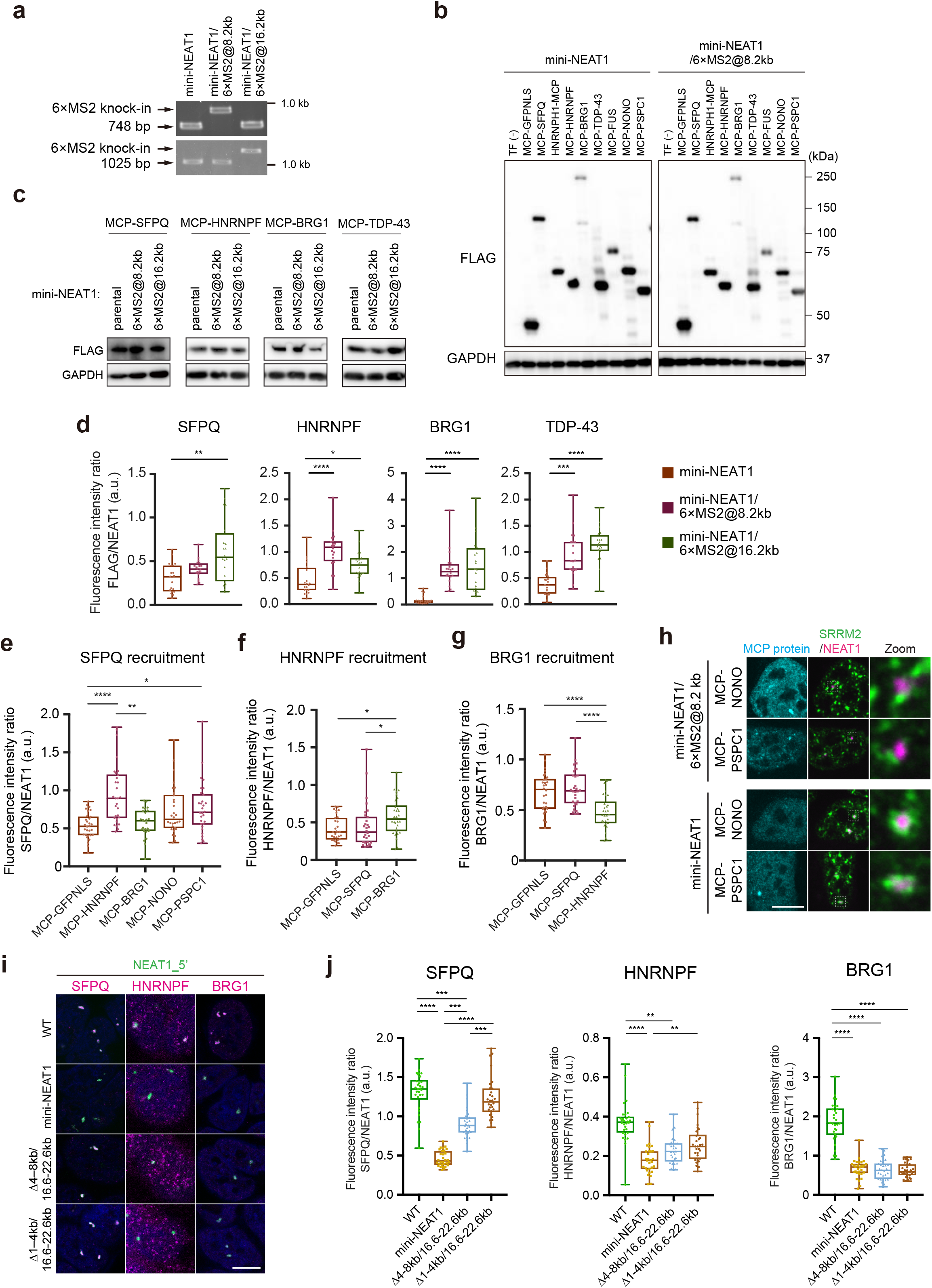
Reciprocal recruitment of SFPQ, HNRNPF, and BRG1 by their tethering. **a**, Genotyping PCR to verify the 6×MS2 knock-in at the *mini-NEAT1* locus. Gel electrophoresis of the PCR products confirmed the 6×MS2 knock-in, represented by upstream band shifts. The primers used in this figure are listed in Supplementary Table S2. **b**, Immunoblotting of Flag-tagged MCP-PSPs in the mini-NEAT1 and mini-NEAT1/6×MS2@8.2 kb cells corresponding to Fig. 3b and 3c. GAPDH was used as a control. Molecular weight markers are shown on the right. **c**, Immunoblotting of Flag-tagged MCP-PSPs in the mini-NEAT1, mini-NEAT1/6×MS2@8.2 kb, and mini-NEAT1/6×MS2@16.2 kb corresponding to Fig. 3b–d. GAPDH was used as a control. Molecular weight markers are shown on the right. **d**, Quantification of fluorescence intensity ratio (FLAG/NEAT1) in the mini-NEAT1, mini-NEAT1/6×MS2@8.2 kb, and mini-NEAT1/6×MS2@16.2 kb cells (all samples: *n* = 20). **e**–**g**, Quantification of fluorescence intensity ratio (PSPs/NEAT1) with transfection of MCP-PSPs into mini-NEAT1/6×MS2@8.2 kb cells. (**e**) SFPQ recruitment: *n* = 30 (MCP-GFPNLS), *n* = 30 (MCP-HNRNPF), *n* = 25 (MCP-BRG1), *n* = 30 (MCP-NONO), *n* = 30 (MCP-PSPC1). (**f**) HNRNPF recruitment: *n* = 30 (MCP-GFPNLS), *n* = 30 (MCP-SFPQ), *n* = 30 (MCP-BRG1). (**g**) BRG1 recruitment: *n* = 30 (MCP-GFPNLS), *n* = 30 (MCP-SFPQ), *n* = 30 (MCP-HNRNPF). **h**, Confocal observation of PSs and NSs with transfection of MCP-NONO or PSPC1 into mini-NEAT1/6×MS2@8.2 kb (upper) and mini-NEAT1 (lower) in the MG132 treatment conditions (5 μM for 6 h). White boxes indicate the areas shown at a higher magnification. Scale bar, 10 μm. **i**, Recruitment of various PSPs to PSs in HAP1 WT and mini-NEAT1, and Δ4–8 kb/16.6–22.6 kb, and Δ1–4 kb/16.6–22.6 kb mutant cells treated with MG132 (5 μM for 6 h). PSs and PSPs were visualized by RNA-FISH using NEAT1_5’ probes (green) and IF (magenta). Nuclei were stained with DAPI. Scale bar, 10 μm. **j**, Quantification of fluorescence intensity ratio (PSPs/NEAT1) in the HAP1 WT, mini-NEAT1, Δ4–8 kb/16.6–22.6 kb, and Δ1–4 kb/16.6–22.6 kb mutant cells as observed in **i** (all samples: *n* = 30). For **d**–**g**,**j**, each box plot shows the median (inside line), 25th-75th percentiles (box bottom to top), and minimum-maximum values (whisker bottom to top). \*\*\*\**P* < 0.0001, \*\*\**P* < 0.001, \*\**P* < 0.01, \**P* < 0.05. Data were compared using Kruskal-Wallis ANOVA and post hoc Dunn’s multiple comparison test. If the statistical test showed no significant difference (*P* > 0.05), it is not specified in the figure. Numerical data and unprocessed blots are available in the Source Data.

**Extended Data Fig. 5.**
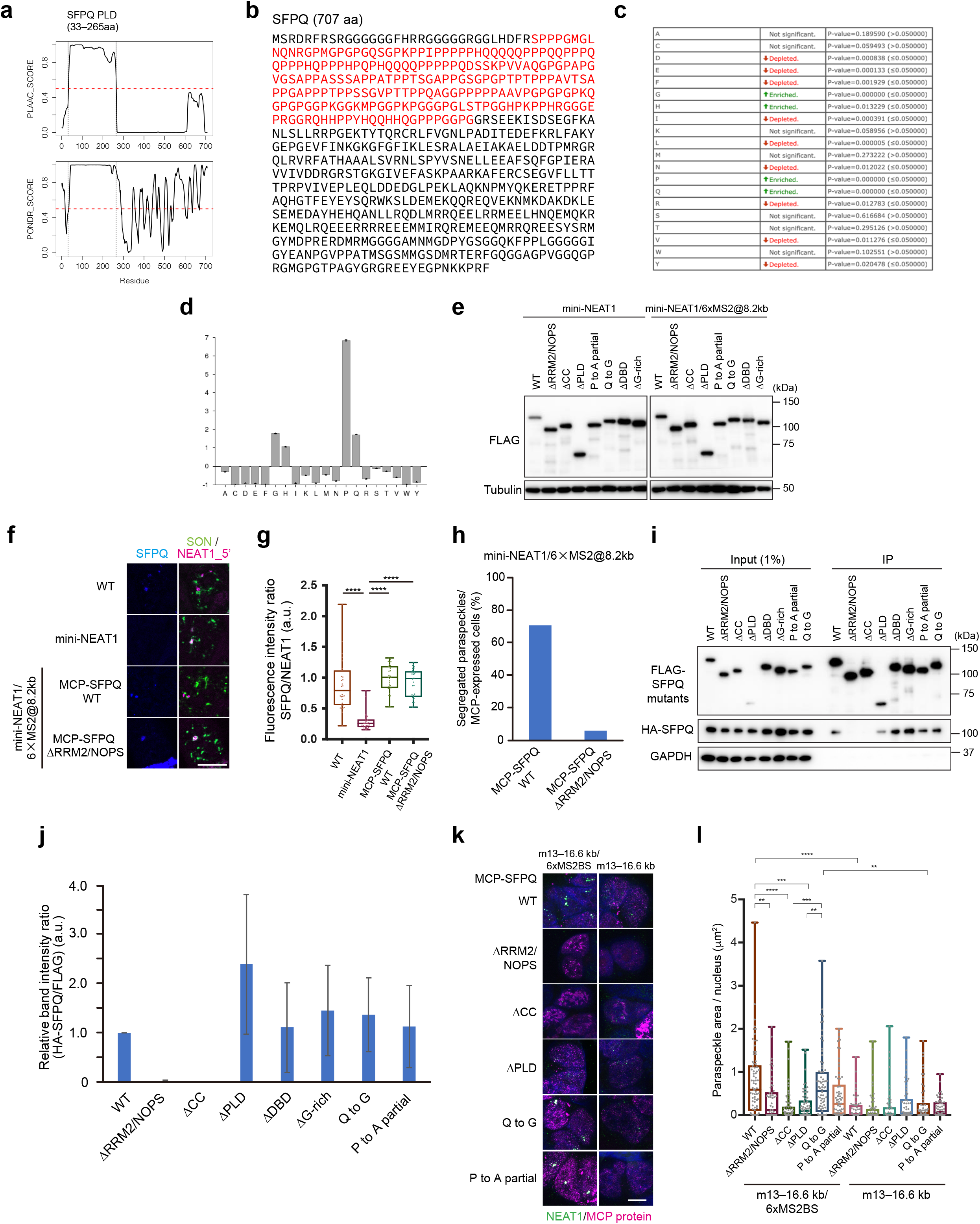
Effects of SFPQ deletion or mutations on interactions between SFPQ and the other DBHS family proteins and PS assembly. **a**, Graph predicting PLD and IDR of SFPQ using PLAAC (upper) and PONDR (lower), respectively. **b**, Amino acid sequence of SFPQ. PLD is highlighted in red. **c**,**d**, Enrichment or depletion patterns of individual amino acids in SFPQ PLD detected by COMPOSITION PROFILER (http://www.cprofiler.org). **e**, Immunoblotting of Flag-tagged MCP-SFPQ WT and mutants in the mini-NEAT1 and mini-NEAT1/6×MS2@8.2 kb cells corresponding to Fig. 4b and c. GAPDH was used as a control. Molecular weight markers are shown on the right. **f**, Confocal observation of the PSs, the NSs, and SFPQ in mini-NEAT1/6×MS2@8.2 kb cells expressing MCP-SFPQ WT or ΔRRM2/NOPS and in HAP1 WT and mini-NEAT1 cells treated with MG132 (5 μM for 6 h). PSs and SFPQ were visualized by NEAT1 RNA-FISH with NEAT1_5’ probe (magenta) and IF of SFPQ (blue). NSs were visualized by IF of SON (green). Scale bar, 10 μm. **g**, Quantification of fluorescence intensity ratio (SFPQ/NEAT1) in mini-NEAT1/6×MS2@8.2 kb cells expressing MCP-SFPQ WT or ΔRRM2/NOPS and in HAP1 WT and mini-NEAT1 cells as observed in **f** (all samples: *n* = 30). **h**, Proportion (%) of the cells with paraspeckles segregated from nuclear speckles in **f**. Data were collected from two independent experiments. mini-NEAT1/6×MS2@8.2 kb: *n* = 58 (MCP-SFPQ WT), *n* = 53 (MCP-SFPQ ΔRRM2/NOPS). **i**, Co-IP of MCP-SFPQ WT and mutant proteins with HA-SFPQ. Immunoblotting was performed to detect HA-tagged SFPQ proteins in the co-IP samples. GAPDH served as a negative control. **j**, Quantification of the data shown in **h**. Values (HA/FLAG) are mean ± SD of three independent experiments. Kruskal-Wallis test showed a statistically significant difference (*P* = 0.03), but post hoc Dunn’s multiple comparison test did not. **k**, Confocal observation of PS formation with transfection of MCP-SFPQ WT and mutant proteins into m13–16.6 kb/6 × MS2BS or m13–16.6 kb cells in the MG132 treatment conditions (5 μM for 6 h). The rescued PSs were visualized by NEAT1 RNA FISH (green) and IF of Flag-tagged MCP-SFPQ WT and mutants (magenta). Scale bar, 10 μm. **l**, Quantification of the PS sizes observed in **k**. m13-16.6 kb/6×MS2BS: *n* = 78 (WT), *n* = 54 (ΔRRM2/NOPS), *n* = 65 (ΔCC), *n* = 62 (ΔPLD), *n* = 78 (Q to G), *n* = 66 (P to A partial); m13-16.6k: *n* = 58 (WT), *n* = 64 (ΔRRM2/NOPS), *n* = 53 (ΔCC), *n* = 62 (ΔPLD), *n* = 60 (Q to G), *n* = 56 (P to A partial). \*\*\*\**P* < 0.0001, \*\*\**P* < 0.001, \*\**P* < 0.01. **g**,**l**, Each box plot shows the median (inside line), 25th-75th percentiles (box bottom to top), and minimum-maximum values (whisker bottom to top). **g**,**j**,**l**, Data were compared using Kruskal-Wallis ANOVA and post hoc Dunn’s multiple comparison test. If the statistical test showed no significant difference (*P* > 0.05), it is not specified in the figure. Numerical data and unprocessed blots are available in the Source Data.

**Extended Data Fig. 6.**
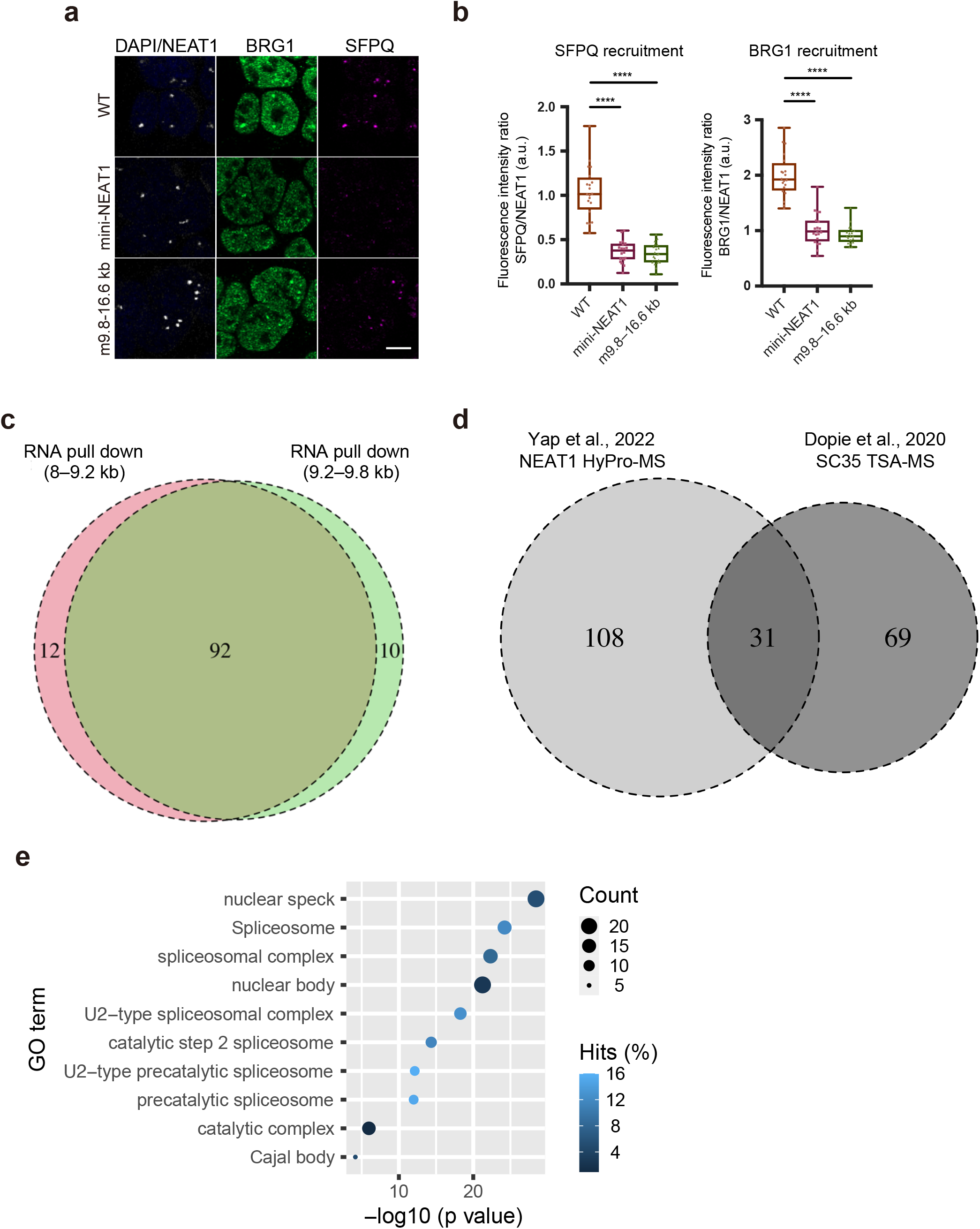
U2 snRNP components are included in both NS and PS proteomes. **a**, Confocal observation of PSs, BRG1, and SFPQ in HAP1 WT, mini-NEAT, and m9.8–16.6 kb mutant cells treated with MG132 (5 μM for 6 h). PSs were visualized by NEAT1 RNA-FISH with NEAT1_5’ probe (white). BRG1 (green) and SFPQ (magenta) were visualized by IF. Nuclei were stained with DAPI. Scale bar, 10 μm. **b**, Quantification of fluorescence intensity ratio (PSPs/NEAT1) observed in **b** (all samples: *n* = 30). Each box plot shows the median (inside line), 25th-75th percentiles (box bottom to top), and minimum-maximum values (whisker bottom to top). \*\*\*\**P* < 0.0001. Data were compared using Kruskal-Wallis ANOVA and post hoc Dunn’s multiple comparison test. If the statistical test showed no significant difference (*P* > 0.05), it is not specified in the figure. **c**,**d**, Venn diagram showing the overlap between 8–9.2 kb and 9.2–9.8 kb of RNA pulldown (**c**), or between NEAT1 HyPro-MS^39^ and SC35 TSA-MS^38^ (**d**). **e**, Top 10 Gene Ontology cellular compartment enrichment for proteins shared between NEAT1 HyPro-MS^39^ and SC35 TSA-MS^38^. Numerical data are available in the Source Data.

**Extended Data Fig. 7.**
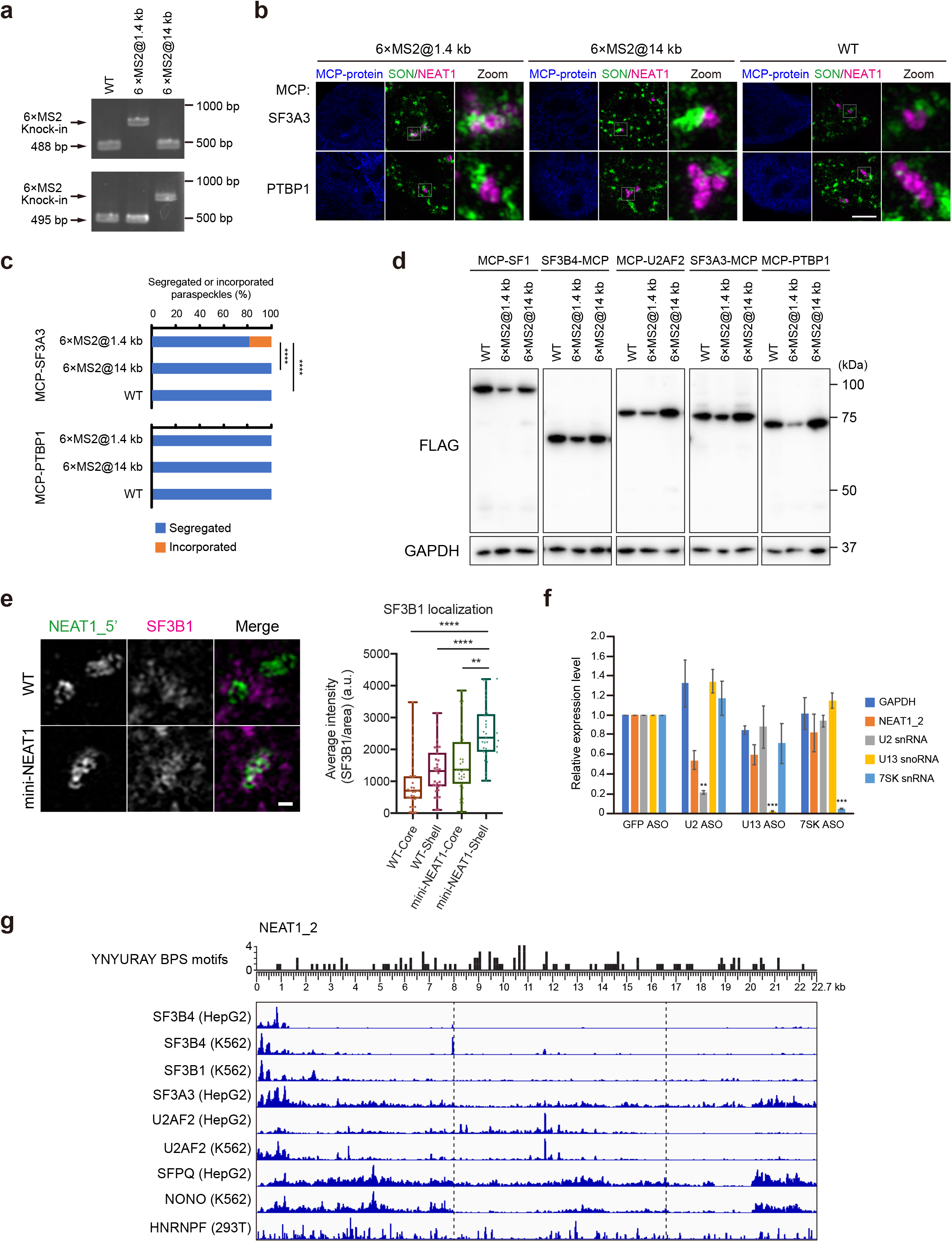
SF3A and SF3B complex components bind to the 5’ terminal shell region of NEAT1, and the NEAT1_2 sequence contains multiple potential branch point sequences. **a**, Genotyping PCR to verify the 6 × MS2 knock-in at the *NEAT1* locus. Gel electrophoresis of the PCR products confirmed the 6 × MS2 knock-in, represented by upstream band shifts. Primers used in this figure are listed in Supplementary Table S1. **b**, SRM maximum projection images of paraspeckles and nuclear speckles with transfection of MCP proteins into NEAT1/6×MS2@1.4 kb (left), NEAT1/6×MS2 @14 kb (middle), and WT (right) in the MG132 treatment conditions (5 μM for 6 h). White boxes indicate the areas shown at a higher magnification. Scale bar, 5 μm. **c**, Proportion (%) of the cells with paraspeckles segregated from nuclear speckles in **b**. Data were collected from three independent experiments. NEAT1/6×MS2@1.4 kb @1.4 kb; *n* = 60 (SF3A3-MCP), *n* = 59 (MCP-PTBP1), NEAT1/6×MS2@1.4 kb @14 kb; *n* = 61 (SF3A3-MCP), *n* = 59 (MCP-PTBP1), WT; *n* = 62 (SF3A3-MCP), *n* = 64 (MCP-PTBP1). \*\*\*\**P* < 0.0001. Data were compared using Fisher’s exact test and Bonferroni correction. If the statistical test showed no significant difference (*P* > 0.05), it is not specified in the figure. **d**, Immunoblotting of Flag-tagged MCP proteins in the experiments in Fig. 6b, c, and Extended Data Fig. 7a, b. GAPDH was used as a control. Molecular weight markers are shown on the right. **e**, SRM images of localization of SF3B1 within paraspeckles in HAP1 WT and mini-NEAT1 cells treated with MG132 (5 μM for 6 h) (upper). Paraspeckles were visualized by NEAT1 RNA FISH with NEAT1_5’ probe (green), and SF3B1 IF (magenta). Scale bar, 500 nm. Graph showing average fluorescence intensity of SF3B1 in the shell or core of paraspeckles (lower) (all samples: *n* = 31). Each box plot shows the median (inside line), 25th-75th percentiles (box bottom to top), and minimum-maximum values (whisker bottom to top). Data were compared using Kruskal-Wallis ANOVA and post hoc Dunn’s multiple comparison test. \*\*\*\**P* < 0.0001, \*\*\**P* < 0.001, \*\**P* < 0.01. If the statistical test showed no significant difference (*P* > 0.05), it is not specified in the figure. **f**, Quantification of U2 snRNA expression levels of the U2 ASO-transfected mini-NEAT1 mutant cells treated with MG132 (5 μM for 6 h) by RT-qPCR in Fig. 6f,g. Data are presented as mean ± SD (*n* = 3). \*\*\**P* < 0.001, \*\**P* < 0.01. Data were compared using one-way ANOVA and post hoc Tukey’s multiple comparison test. **g**, eCLIP data of SF3B4 (HepG2 and K562), SF3B1 (K562), SF3A3 (HepG2), U2AF2 (HepG2 and K562), SFPQ (HepG2), and NONO (K562) mapped on NEAT1_2. HITS-CLIP data of HNRNPF (293T) are also mapped on NEAT1_2^53^. The number of SF1 consensus motifs (YNYURAY; Y = U or C, R = A or G, N = any nucleotide) present in the NEAT1_2 sequence is shown above with a scale^64^. Numerical data and unprocessed blots are available in the Source Data.

**Extended Data Fig. 8.**
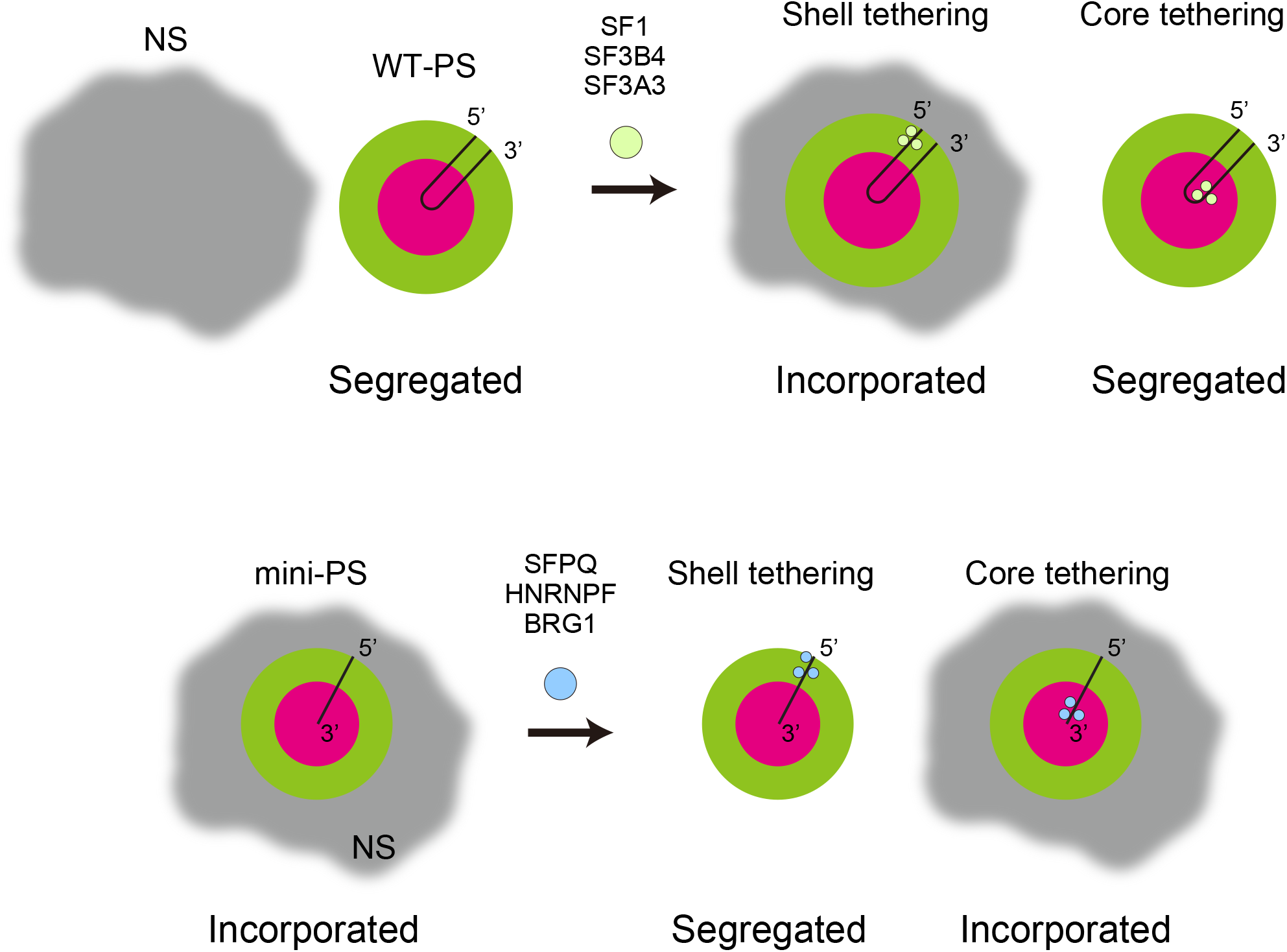
Shell protein composition, but not core protein composition, of the WT-PS or mini-PS dictates their localization to NS. A summary of the shell or core tethering in the WT-PS (upper) and mini-PS (lower).

**Supplementary Video 1.** 3D visualization of PSs and NSs in HAP1 WT cells in the presence of MG132 (5 μM for 6 h), corresponding to Fig. 1e. PSs were visualized by NEAT1 RNA FISH with NEAT1_5’ probe (green), and NSs were visualized by SRRM2 IF (magenta).

**Supplementary Video 2.** 3D visualization of PSs and NSs in mini-NEAT1 mutant cells in the presence of MG132 (5 μM for 6 h), corresponding to Fig 1e. PSs were visualized by NEAT1 RNA FISH with NEAT1_5’ probe (green), and NSs were visualized by SRRM2 IF (magenta).

## References

1. Banani, S. F., Lee, H. O., Hyman, A. A. & Rosen, M. K. Biomolecular condensates: organizers of cellular biochemistry. Nat Rev Mol Cell Biol 18, 285– 298 (2017).

2. Shin, Y. & Brangwynne, C. P. Liquid phase condensation in cell physiology and disease. Science (1979) 357, eaaf4382 (2017).

3. Alberti, S., Gladfelter, A. & Mittag, T. Considerations and Challenges in Studying Liquid-Liquid Phase Separation and Biomolecular Condensates. Cell 176, 419– 434 (2019).

4. Sabari, B. R., Dall’Agnese, A. & Young, R. A. Biomolecular Condensates in the Nucleus. Trends Biochem Sci 45, 961–977 (2020).

5. Lyon, A. S., Peeples, W. B. & Rosen, M. K. A framework for understanding the functions of biomolecular condensates across scales. Nat Rev Mol Cell Biol 22, 215–235 (2021).

6. Yamazaki, T. & Hirose, T. Control of condensates dictates nucleolar architecture. Science (1979) 373, 486–487 (2021).

7. Roden, C. & Gladfelter, A. S. RNA contributions to the form and function of biomolecular condensates. Nat Rev Mol Cell Biol 22, 183–195 (2021).

8. Van Treeck, B. & Parker, R. Emerging Roles for Intermolecular RNA-RNA Interactions in RNP Assemblies. Cell 174, 791–802 (2018).

9. Yamazaki, T., Nakagawa, S. & Hirose, T. Architectural RNAs for Membraneless Nuclear Body Formation. Cold Spring Harb Symp Quant Biol 84, 227–237 (2019).

10. Chujo, T. & Hirose, T. Nuclear Bodies Built on Architectural Long Noncoding RNAs: Unifying Principles of Their Construction and Function. Mol Cells 40, 889– 896 (2017).

11. Chen, L.-L. & Carmichael, G. G. Altered Nuclear Retention of mRNAs Containing Inverted Repeats in Human Embryonic Stem Cells: Functional Role of a Nuclear Noncoding RNA. Mol Cell 35, 467–478 (2009).

12. Clemson, C. M. et al. An Architectural Role for a Nuclear Noncoding RNA: NEAT1 RNA Is Essential for the Structure of Paraspeckles. Mol Cell 33, 717–726 (2009).

13. Sasaki, Y. T. F., Ideue, T., Sano, M., Mituyama, T. & Hirose, T. MENε/β noncoding RNAs are essential for structural integrity of nuclear paraspeckles. Proceedings of the National Academy of Sciences 106, 2525–2530 (2009).

14. Sunwoo, H., et al. *MEN ε/β* nuclear-retained non-coding RNAs are up-regulated upon muscle differentiation and are essential components of paraspeckles. Genome Res 19, 347–359 (2009).

15. Fox, A. H. et al. Paraspeckles: A Novel Nuclear Domain. Current Biology 12, 13– 25 (2002).

16. Visa, N., Puvion-Dutilleul, F., Bachellerie, J. P. & Puvion, E. Intranuclear distribution of U1 and U2 snRNAs visualized by high resolution in situ hybridization: Revelation of a novel compartment containing U1 but not U2 snRNA in HeLa cells. Eur J Cell Biol 60, 308–321 (1993).

17. Hirose, T. et al. NEAT1 long noncoding RNA regulates transcription via protein sequestration within subnuclear bodies. Mol Biol Cell 25, 169–183 (2014).

18. Imamura, K. et al. Long Noncoding RNA NEAT1-Dependent SFPQ Relocation from Promoter Region to Paraspeckle Mediates IL8 Expression upon Immune Stimuli. Mol Cell 53, 393–406 (2014).

19. Nakagawa, S. et al. The lncRNA *Neat1* is required for corpus luteum formation and the establishment of pregnancy in a subpopulation of mice. Development 141, 4618–4627 (2014).

20. Adriaens, C. et al. p53 induces formation of NEAT1 lncRNA-containing paraspeckles that modulate replication stress response and chemosensitivity. Nat Med 22, 861–868 (2016).

21. Mello, S. S., et al. *Neat1* is a p53-inducible lincRNA essential for transformation suppression. Genes Dev 31, 1095–1108 (2017).

22. Naganuma, T. et al. Alternative 3 ′-end processing of long noncoding RNA initiates construction of nuclear paraspeckles. EMBO J 31, 4020–4034 (2012).

23. Yamazaki, T. et al. Functional Domains of NEAT1 Architectural lncRNA Induce Paraspeckle Assembly through Phase Separation. Mol Cell 70, 1038–1053.e7 (2018).

24. Wilusz, J. E. et al. A triple helix stabilizes the 3′ ends of long noncoding RNAs that lack poly(A) tails. Genes Dev 26, 2392–2407 (2012).

25. Fox, A. H., Bond, C. S. & Lamond, A. I. P54nrb Forms a Heterodimer with PSP1 That Localizes to Paraspeckles in an RNA-dependent Manner. Mol Biol Cell 16, 5304–5315 (2005).

26. Mao, Y. S., Sunwoo, H., Zhang, B. & Spector, D. L. Direct visualization of the co-transcriptional assembly of a nuclear body by noncoding RNAs. Nat Cell Biol 13, 95–101 (2011).

27. Kawaguchi, T. et al. SWI/SNF chromatin-remodeling complexes function in noncoding RNA-dependent assembly of nuclear bodies. Proceedings of the National Academy of Sciences 112, 4304–4309 (2015).

28. Yamazaki, T. & Hirose, T. The building process of the functional paraspeckle with long non-coding RNAs. Frontiers in Bioscience 7, 715 (2015).

29. Hennig, S. et al. Prion-like domains in RNA binding proteins are essential for building subnuclear paraspeckles. Journal of Cell Biology 210, 529–539 (2015).

30. Modic, M. et al. Cross-Regulation between TDP-43 and Paraspeckles Promotes Pluripotency-Differentiation Transition. Mol Cell 74, 951–965.e13 (2019).

31. Yamazaki, T. et al. Paraspeckles are constructed as block copolymer micelles. EMBO J 40, e107270 (2021).

32. Souquere, S., Beauclair, G., Harper, F., Fox, A. & Pierron, G. Highly Ordered Spatial Organization of the Structural Long Noncoding NEAT1 RNAs within Paraspeckle Nuclear Bodies. Mol Biol Cell 21, 4020–4027 (2010).

33. West, J. A. et al. Structural, super-resolution microscopy analysis of paraspeckle nuclear body organization. Journal of Cell Biology 214, 817–830 (2016).

34. Yamazaki, T., Yamamoto, T. & Hirose, T. Micellization: A new principle in the formation of biomolecular condensates. Front Mol Biosci 9, (2022).

35. Chujo, T. et al. Unusual semi-extractability as a hallmark of nuclear body-associated architectural noncoding RNAs. EMBO J 36, 1447–1462 (2017).

36. Huang, J. et al. Crystal structure of a SFPQ/PSPC1 heterodimer provides insights into preferential heterodimerization of human DBHS family proteins. Journal of Biological Chemistry 293, 6593–6602 (2018).

37. Lee, M. et al. The structure of human SFPQ reveals a coiled-coil mediated polymer essential for functional aggregation in gene regulation. Nucleic Acids Res 43, 3826–3840 (2015).

38. Dopie, J., Sweredoski, M. J., Moradian, A. & Belmont, A. S. Tyramide signal amplification mass spectrometry (TSA-MS) ratio identifies nuclear speckle proteins. Journal of Cell Biology 219, e201910207 (2020).

39. Yap, K., Chung, T. H. & Makeyev, E. V. Hybridization-proximity labeling reveals spatially ordered interactions of nuclear RNA compartments. Mol Cell 82, 463–478.e11 (2022).

40. Sanders, D. W. et al. Competing Protein-RNA Interaction Networks Control Multiphase Intracellular Organization. Cell 181, 306–324.e28 (2020).

41. Kaida, D. et al. Spliceostatin A targets SF3b and inhibits both splicing and nuclear retention of pre-mRNA. Nat Chem Biol 3, 576–583 (2007).

42. Kotake, Y. et al. Splicing factor SF3b as a target of the antitumor natural product pladienolide. Nat Chem Biol 3, 570–575 (2007).

43. Iida, K., Hagiwara, M. & Takeuchi, A. Multilateral Bioinformatics Analyses Reveal the Function-Oriented Target Specificities and Recognition of the RNA-Binding Protein SFPQ. iScience 23, (2020).

44. Dias, A. P., Dufu, K., Lei, H. & Reed, R. A role for TREX components in the release of spliced mRNA from nuclear speckle domains. Nat Commun 1, 97 (2010).

45. Brown, J. A., Valenstein, M. L., Yario, T. A., Tycowski, K. T. & Steitz, J. A. Formation of triple-helical structures by the 3′-end sequences of MALAT1 and MEN β noncoding RNAs. Proceedings of the National Academy of Sciences 109, 19202– 19207 (2012).

46. Watanabe, Y. & Yamamoto, M. S. pombe mei2+ encodes an RNA-binding protein essential for premeiotic DNA synthesis and meiosis I, which cooperates with a novel RNA species meiRNA. Cell 78, 487–498 (1994).

47. Prasanth, K. V., Rajendra, T. K., Lal, A. K. & Lakhotia, S. C. Omega speckles - a novel class of nuclear speckles containing hnRNPs associated with noncoding hsr-omega RNA in Drosophila. J Cell Sci 113, 3485–3497 (2000).

48. Biamonti, G. & Vourc’h, C. Nuclear Stress Bodies. Cold Spring Harb Perspect Biol 2, a000695–a000695 (2010).

49. Audas, T. E., Jacob, M. D. & Lee, S. Immobilization of Proteins in the Nucleolus by Ribosomal Intergenic Spacer Noncoding RNA. Mol Cell 45, 147–157 (2012).

50. Gordon, P. M., Hamid, F., Makeyev, E. V. & Houart, C. A conserved role for the ALS-linked splicing factor SFPQ in repression of pathogenic cryptic last exons. Nat Commun 12, 1918 (2021).

51. Stagsted, L. V. W., O’Leary, E. T., Ebbesen, K. K. & Hansen, T. B. The RNA-binding protein SFPQ preserves long-intron splicing and regulates circRNA biogenesis in mammals. Elife 10, e63088 (2021).

52. Gañez-Zapater, A. et al. The SWI/SNF subunit BRG1 affects alternative splicing by changing RNA binding factor interactions with nascent RNA. Molecular Genetics and Genomics 297, 463–484 (2022).

53. Huelga, S. C. et al. Integrative Genome-wide Analysis Reveals Cooperative Regulation of Alternative Splicing by hnRNP Proteins. Cell Rep 1, 167–178 (2012).

54. Courchaine, E. M. et al. DMA-tudor interaction modules control the specificity of in vivo condensates. Cell 184, 3612–3625.e17 (2021).

55. Schmidt, U. et al. Real-time imaging of cotranscriptional splicing reveals a kinetic model that reduces noise: Implications for alternative splicing regulation. Journal of Cell Biology 193, (2011).

56. Brody, Y. et al. The in vivo kinetics of RNA polymerase II elongation during co-transcriptional splicing. PLoS Biol 9, (2011).

57. Kawaguchi, T. & Hirose, T. Chromatin remodeling complexes in the assembly of long noncoding RNA-dependent nuclear bodies. Nucleus 6, 462–467 (2015).

58. Yamazaki, T. & Hirose, T. CRISPR-Mediated Mutagenesis of Long Noncoding RNAs. in Methods in Molecular Biology vol. 2254 283–303 (2021).

59. Yamazaki, T. et al. FUS-SMN Protein Interactions Link the Motor Neuron Diseases ALS and SMA. Cell Rep 2, 799–806 (2012).

60. Ilik, İ. A., et al. SON and SRRM2 are essential for nuclear speckle formation. Elife 9, e60579 (2020).

61. Lee, K. A. W. & Green, M. R. Small-scale preparation of extracts from radiolabeled cells efficient in pre-mRNA splicing. in Methods in Enzymology vol. 181 20–30 (1990).

62. Perez, C. A. G. et al. Sense-overlapping lncRNA as a decoy of translational repressor protein for dimorphic gene expression. PLoS Genet 17, e1009683 (2021).

63. Ideue, T., Hino, K., Kitao, S., Yokoi, T. & Hirose, T. Efficient oligonucleotide-mediated degradation of nuclear noncoding RNAs in mammalian cultured cells. RNA 15, 1578–1587 (2009).

64. Gao, K., Masuda, A., Matsuura, T. & Ohno, K. Human branch point consensus sequence is yUnAy. Nucleic Acids Res 36, 2257–2267 (2008).

