## Supplementary Table 1 for "Shell protein composition specified by NEAT1 domains dictates the formation of paraspeckles as distinct membraneless organelles"

hNEAT1 8-9.2 kb & 9.2-9.8 kb RNA pulldown & MS spectrometry analysis

| | initial_alias | converted_alias | name | description | unique peptide # |
|---|---|---|---|---|---|
| 1 | SF3B1 | ENSG00000115524 | SF3B1 | splicing factor 3b subunit 1 [Source:HGNC Symbol;Acc:HGNC:10768] | 215 |
| 2 | DHX15 | ENSG00000109606 | DHX15 | DEAH-box helicase 15 [Source:HGNC Symbol;Acc:HGNC:2738] | 183 |
| 3 | SF3B3 | ENSG00000189091 | SF3B3 | splicing factor 3b subunit 3 [Source:HGNC Symbol;Acc:HGNC:10770] | 162 |
| 4 | CCAR1 | ENSG00000060339 | CCAR1 | cell division cycle and apoptosis regulator 1 [Source:HGNC Symbol;Acc:HGNC:24236] | 128 |
| 5 | U2SURP | ENSG00000163714 | U2SURP | U2 snRNP associated SURP domain containing [Source:HGNC Symbol;Acc:HGNC:30855] | 121 |
| 6 | SF3B2 | ENSG00000087365 | SF3B2 | splicing factor 3b subunit 2 [Source:HGNC Symbol;Acc:HGNC:10769] | 114 |
| 7 | HNRNPL | ENSG00000104824 | HNRNPL | heterogeneous nuclear ribonucleoprotein L [Source:HGNC Symbol;Acc:HGNC:5045] | 100 |
| 8 | HNRNPUL1 | ENSG00000105323 | HNRNPUL1 | heterogeneous nuclear ribonucleoprotein U like 1 [Source:HGNC Symbol;Acc:HGNC:17011] | 99 |
| 9 | PTBP1 | ENSG00000011304 | PTBP1 | polypyrimidine tract binding protein 1 [Source:HGNC Symbol;Acc:HGNC:9583] | 97 |
| 10 | PUF60 | ENSG00000179950 | PUF60 | poly(U) binding splicing factor 60 [Source:HGNC Symbol;Acc:HGNC:17042] | 95 |
| 11 | SF3A3 | ENSG00000183431 | SF3A3 | splicing factor 3a subunit 3 [Source:HGNC Symbol;Acc:HGNC:10767] | 93 |
| 12 | SNRNP200 | ENSG00000144028 | SNRNP200 | small nuclear ribonucleoprotein U5 subunit 200 [Source:HGNC Symbol;Acc:HGNC:30859] | 91 |
| 13 | SF3A1 | ENSG00000099995 | SF3A1 | splicing factor 3a subunit 1 [Source:HGNC Symbol;Acc:HGNC:10765] | 91 |
| 14 | KHSRP | ENSG00000088247 | KHSRP | KH-type splicing regulatory protein [Source:HGNC Symbol;Acc:HGNC:6316] | 90 |
| 15 | HNRNPA2B1 | ENSG00000122566 | HNRNPA2B1 | heterogeneous nuclear ribonucleoprotein A2/B1 [Source:HGNC Symbol;Acc:HGNC:5033] | 87 |
| 16 | U2AF2 | ENSG00000063244 | U2AF2 | U2 small nuclear RNA auxiliary factor 2 [Source:HGNC Symbol;Acc:HGNC:23156] | 82 |
| 17 | DDX5 | ENSG00000108654 | DDX5 | DEAD-box helicase 5 [Source:HGNC Symbol;Acc:HGNC:2746] | 79 |
| 18 | DHX9 | ENSG00000135829 | DHX9 | DExH-box helicase 9 [Source:HGNC Symbol;Acc:HGNC:2750] | 79 |
| 19 | CHERP | ENSG00000085872 | CHERP | calcium homeostasis endoplasmic reticulum protein [Source:HGNC Symbol;Acc:HGNC:16930] | 74 |
| 20 | FUBP1 | ENSG00000162613 | FUBP1 | far upstream element binding protein 1 [Source:HGNC Symbol;Acc:HGNC:4004] | 71 |
| 21 | RBM17 | ENSG00000134453 | RBM17 | RNA binding motif protein 17 [Source:HGNC Symbol;Acc:HGNC:16944] | 70 |
| 22 | PRPF8 | ENSG00000174231 | PRPF8 | pre-mRNA processing factor 8 [Source:HGNC Symbol;Acc:HGNC:17340] | 69 |
| 23 | SF1 | ENSG00000168066 | SF1 | splicing factor 1 [Source:HGNC Symbol;Acc:HGNC:12950] | 69 |
| 24 | ELAVL1 | ENSG00000066044 | ELAVL1 | ELAV like RNA binding protein 1 [Source:HGNC Symbol;Acc:HGNC:3312] | 61 |
| 25 | RBM25 | ENSG00000119707 | RBM25 | RNA binding motif protein 25 [Source:HGNC Symbol;Acc:HGNC:23244] | 60 |
| 26 | DAZAP1 | ENSG00000071626 | DAZAP1 | DAZ associated protein 1 [Source:HGNC Symbol;Acc:HGNC:2683] | 59 |
| 27 | MATR3 | ENSG00000015479 | MATR3 | matrin 3 [Source:HGNC Symbol;Acc:HGNC:6912] | 58 |
| 28 | HNRNPR | ENSG00000125944 | HNRNPR | heterogeneous nuclear ribonucleoprotein R [Source:HGNC Symbol;Acc:HGNC:5047] | 55 |
| 29 | RBM39 | ENSG00000131051 | RBM39 | RNA binding motif protein 39 [Source:HGNC Symbol;Acc:HGNC:15923] | 54 |
| 30 | HNRNPA3 | ENSG00000170144 | HNRNPA3 | heterogeneous nuclear ribonucleoprotein A3 [Source:HGNC Symbol;Acc:HGNC:24941] | 54 |
| 31 | EFTUD2 | ENSG00000108883 | EFTUD2 | elongation factor Tu GTP binding domain containing 2 [Source:HGNC Symbol;Acc:HGNC:30858] | 54 |
| 32 | SRSF1 | ENSG00000136450 | SRSF1 | serine and arginine rich splicing factor 1 [Source:HGNC Symbol;Acc:HGNC:10780] | 53 |
| 33 | RBM10 | ENSG00000182872 | RBM10 | RNA binding motif protein 10 [Source:HGNC Symbol;Acc:HGNC:9896] | 52 |
| 34 | EIF4A3 | ENSG00000141543 | EIF4A3 | eukaryotic translation initiation factor 4A3 [Source:HGNC Symbol;Acc:HGNC:18683] | 51 |
| 35 | CPSF1 | ENSG00000071894 | CPSF1 | cleavage and polyadenylation specific factor 1 [Source:HGNC Symbol;Acc:HGNC:2324] | 50 |
| 36 | HNRNPK | ENSG00000165119 | HNRNPK | heterogeneous nuclear ribonucleoprotein K [Source:HGNC Symbol;Acc:HGNC:5044] | 49 |
| 37 | HNRNPM | ENSG00000099783 | HNRNPM | heterogeneous nuclear ribonucleoprotein M [Source:HGNC Symbol;Acc:HGNC:5046] | 48 |
| 38 | SNRPA1 | ENSG00000131876 | SNRPA1 | small nuclear ribonucleoprotein polypeptide A' [Source:HGNC Symbol;Acc:HGNC:11152] | 48 |
| 39 | SNRNP70 | ENSG00000104852 | SNRNP70 | small nuclear ribonucleoprotein U1 subunit 70 [Source:HGNC Symbol;Acc:HGNC:11150] | 47 |
| 40 | HNRNPH1 | ENSG00000169045 | HNRNPH1 | heterogeneous nuclear ribonucleoprotein H1 [Source:HGNC Symbol;Acc:HGNC:5041] | 44 |
| 41 | PCBP1 | ENSG00000169564 | PCBP1 | poly(rC) binding protein 1 [Source:HGNC Symbol;Acc:HGNC:8647] | 42 |
| 42 | SF3B6 | ENSG00000115128 | SF3B6 | splicing factor 3b subunit 6 [Source:HGNC Symbol;Acc:HGNC:30096] | 37 |
| 43 | CPSF2 | ENSG00000165934 | CPSF2 | cleavage and polyadenylation specific factor 2 [Source:HGNC Symbol;Acc:HGNC:2325] | 36 |
| 44 | HNRNPDL | ENSG00000152795 | HNRNPDL | heterogeneous nuclear ribonucleoprotein D like [Source:HGNC Symbol;Acc:HGNC:5037] | 36 |
| 45 | FUS | ENSG00000089280 | FUS | FUS RNA binding protein [Source:HGNC Symbol;Acc:HGNC:4010] | 35 |
| 46 | PRPF40A | ENSG00000196504 | PRPF40A | pre-mRNA processing factor 40 homolog A [Source:HGNC Symbol;Acc:HGNC:16463] | 34 |
| 47 | SF3A2 | ENSG00000104897 | SF3A2 | splicing factor 3a subunit 2 [Source:HGNC Symbol;Acc:HGNC:10766] | 34 |
| 48 | ACIN1 | ENSG00000100813 | ACIN1 | apoptotic chromatin condensation inducer 1 [Source:HGNC Symbol;Acc:HGNC:17066] | 34 |
| 49 | HNRNPA0 | ENSG00000177733 | HNRNPA0 | heterogeneous nuclear ribonucleoprotein A0 [Source:HGNC Symbol;Acc:HGNC:5030] | 33 |
| 50 | SF3B4 | ENSG00000143368 | SF3B4 | splicing factor 3b subunit 4 [Source:HGNC Symbol;Acc:HGNC:10771] | 32 |
| 51 | SYNCRIP | ENSG00000135316 | SYNCRIP | synaptotagmin binding cytoplasmic RNA interacting protein [Source:HGNC Symbol;Acc:HGNC:16918] | 32 |
| 52 | ALYREF | ENSG00000183684 | ALYREF | Aly/REF export factor [Source:HGNC Symbol;Acc:HGNC:19071] | 31 |
| 53 | DDX3X | ENSG00000215301 | DDX3X | DEAD-box helicase 3 X-linked [Source:HGNC Symbol;Acc:HGNC:2745] | 31 |
| 54 | RBMX | ENSG00000147274 | RBMX | RNA binding motif protein X-linked [Source:HGNC Symbol;Acc:HGNC:9910] | 31 |
| 55 | SNRPD3 | ENSG00000100028 | SNRPD3 | small nuclear ribonucleoprotein D3 polypeptide [Source:HGNC Symbol;Acc:HGNC:11160] | 30 |
| 56 | THRAP3 | ENSG00000054118 | THRAP3 | thyroid hormone receptor associated protein 3 [Source:HGNC Symbol;Acc:HGNC:22964] | 30 |
| 57 | CPSF7 | ENSG00000149532 | CPSF7 | cleavage and polyadenylation specific factor 7 [Source:HGNC Symbol;Acc:HGNC:30098] | 27 |
| 58 | SRSF7 | ENSG00000115875 | SRSF7 | serine and arginine rich splicing factor 7 [Source:HGNC Symbol;Acc:HGNC:10789] | 27 |
| 59 | LUC7L3 | ENSG00000108848 | LUC7L3 | LUC7 like 3 pre-mRNA splicing factor [Source:HGNC Symbol;Acc:HGNC:24309] | 27 |
| 60 | NUDT21 | ENSG00000167005 | NUDT21 | nudix hydrolase 21 [Source:HGNC Symbol;Acc:HGNC:13870] | 25 |
| 61 | HNRNPAB | ENSG00000197451 | HNRNPAB | heterogeneous nuclear ribonucleoprotein A/B [Source:HGNC Symbol;Acc:HGNC:5034] | 25 |
| 62 | HNRNPD | ENSG00000138668 | HNRNPD | heterogeneous nuclear ribonucleoprotein D [Source:HGNC Symbol;Acc:HGNC:5036] | 25 |
| 63 | SRSF11 | ENSG00000116754 | SRSF11 | serine and arginine rich splicing factor 11 [Source:HGNC Symbol;Acc:HGNC:10782] | 25 |
| 64 | RPS7 | ENSG00000171863 | RPS7 | ribosomal protein S7 [Source:HGNC Symbol;Acc:HGNC:10440] | 23 |
| 65 | SNRPB2 | ENSG00000125870 | SNRPB2 | small nuclear ribonucleoprotein polypeptide B2 [Source:HGNC Symbol;Acc:HGNC:11155] | 23 |
| 66 | FIP1L1 | ENSG00000145216 | FIP1L1 | factor interacting with PAPOLA and CPSF1 [Source:HGNC Symbol;Acc:HGNC:19124] | 22 |
| 67 | SRRM2 | ENSG00000167978 | SRRM2 | serine/arginine repetitive matrix 2 [Source:HGNC Symbol;Acc:HGNC:16639] | 22 |
| 68 | EWSR1 | ENSG00000182944 | EWSR1 | EWS RNA binding protein 1 [Source:HGNC Symbol;Acc:HGNC:3508] | 21 |
| 69 | RPS4X | ENSG00000198034 | RPS4X | ribosomal protein S4 X-linked [Source:HGNC Symbol;Acc:HGNC:10424] | 21 |
| 70 | RPL31 | ENSG00000071082 | RPL31 | ribosomal protein L31 [Source:HGNC Symbol;Acc:HGNC:10334] | 20 |
| 71 | DDX17 | ENSG00000100201 | DDX17 | DEAD-box helicase 17 [Source:HGNC Symbol;Acc:HGNC:2740] | 20 |
| 72 | SNRPD1 | ENSG00000167088 | SNRPD1 | small nuclear ribonucleoprotein D1 polypeptide [Source:HGNC Symbol;Acc:HGNC:11158] | 20 |
| 73 | TIAL1 | ENSG00000151923 | TIAL1 | TIA1 cytotoxic granule associated RNA binding protein like 1 [Source:HGNC Symbol;Acc:HGNC:11804] | 20 |
| 74 | KHDRBS1 | ENSG00000121774 | KHDRBS1 | KH RNA binding domain containing, signal transduction associated 1 [Source:HGNC Symbol;Acc:HGNC:18116] | 19 |
| 75 | BUB3 | ENSG00000154473 | BUB3 | BUB3 mitotic checkpoint protein [Source:HGNC Symbol;Acc:HGNC:1151] | 18 |
| 76 | DDX42 | ENSG00000198231 | DDX42 | DEAD-box helicase 42 [Source:HGNC Symbol;Acc:HGNC:18676] | 18 |
| 77 | WDR33 | ENSG00000136709 | WDR33 | WD repeat domain 33 [Source:HGNC Symbol;Acc:HGNC:25651] | 17 |
| 78 | CSTF3 | ENSG00000176102 | CSTF3 | cleavage stimulation factor subunit 3 [Source:HGNC Symbol;Acc:HGNC:2485] | 17 |
| 79 | PNN | ENSG00000100941 | PNN | pinin, desmosome associated protein [Source:HGNC Symbol;Acc:HGNC:9162] | 17 |
| 80 | CPSF6 | ENSG00000111605 | CPSF6 | cleavage and polyadenylation specific factor 6 [Source:HGNC Symbol;Acc:HGNC:13871] | 17 |
| 81 | SAP18 | ENSG00000150459 | SAP18 | Sin3A associated protein 18 [Source:HGNC Symbol;Acc:HGNC:10530] | 17 |
| 82 | SRSF3 | ENSG00000112081 | SRSF3 | serine and arginine rich splicing factor 3 [Source:HGNC Symbol;Acc:HGNC:10785] | 17 |
| 83 | THOC2 | ENSG00000125676 | THOC2 | THO complex 2 [Source:HGNC Symbol;Acc:HGNC:19073] | 17 |
| 84 | DHX36 | ENSG00000174953 | DHX36 | DEAH-box helicase 36 [Source:HGNC Symbol;Acc:HGNC:14410] | 16 |
| 85 | IGF2BP1 | ENSG00000159217 | IGF2BP1 | insulin like growth factor 2 mRNA binding protein 1 [Source:HGNC Symbol;Acc:HGNC:28866] | 16 |
| 86 | HNRNPU | ENSG00000153187 | HNRNPU | heterogeneous nuclear ribonucleoprotein U [Source:HGNC Symbol;Acc:HGNC:5048] | 15 |
| 87 | PRPF4B | ENSG00000112739 | PRPF4B | pre-mRNA processing factor 4B [Source:HGNC Symbol;Acc:HGNC:17346] | 15 |
| 88 | ILF2 | ENSG00000143621 | ILF2 | interleukin enhancer binding factor 2 [Source:HGNC Symbol;Acc:HGNC:6037] | 14 |
| 89 | CIRBP | ENSG00000099622 | CIRBP | cold inducible RNA binding protein [Source:HGNC Symbol;Acc:HGNC:1982] | 14 |
| 90 | NONO | ENSG00000147140 | NONO | non-POU domain containing octamer binding [Source:HGNC Symbol;Acc:HGNC:7871] | 13 |
| 91 | SFPQ | ENSG00000116560 | SFPQ | splicing factor proline and glutamine rich [Source:HGNC Symbol;Acc:HGNC:10774] | 13 |
| 92 | SRRM1 | ENSG00000133226 | SRRM1 | serine and arginine repetitive matrix 1 [Source:HGNC Symbol;Acc:HGNC:16638] | 13 |
| 93 | SNRPD2 | ENSG00000125743 | SNRPD2 | small nuclear ribonucleoprotein D2 polypeptide [Source:HGNC Symbol;Acc:HGNC:11159] | 13 |
| 94 | SMU1 | ENSG00000122692 | SMU1 | SMU1 DNA replication regulator and spliceosomal factor [Source:HGNC Symbol;Acc:HGNC:18247] | 12 |
| 95 | TRA2B | ENSG00000136527 | TRA2B | transformer 2 beta homolog [Source:HGNC Symbol;Acc:HGNC:10781] | 12 |
| 96 | SREK1 | ENSG00000153914 | SREK1 | splicing regulatory glutamic acid and lysine rich protein 1 [Source:HGNC Symbol;Acc:HGNC:17882] | 12 |
| 97 | CPSF3 | ENSG00000119203 | CPSF3 | cleavage and polyadenylation specific factor 3 [Source:HGNC Symbol;Acc:HGNC:2326] | 11 |
| 98 | POLDIP3 | ENSG00000100227 | POLDIP3 | DNA polymerase delta interacting protein 3 [Source:HGNC Symbol;Acc:HGNC:23782] | 11 |
| 99 | HNRNPC | ENSG00000092199 | HNRNPC | heterogeneous nuclear ribonucleoprotein C [Source:HGNC Symbol;Acc:HGNC:5035] | 10 |
| 100 | HNRNPF | ENSG00000169813 | HNRNPF | heterogeneous nuclear ribonucleoprotein F [Source:HGNC Symbol;Acc:HGNC:5039] | 10 |
| 101 | TRIR | ENSG00000123144 | TRIR | telomerase RNA component interacting RNase [Source:HGNC Symbol;Acc:HGNC:28424] | 10 |

hNEAT1 8-9.2 kb RNA pulldown & MS spectrometry analysis

| | initial_alias | converted_alias | name | description | unique peptode # |
|---|---|---|---|---|---|
| 1 | SF3B1 | ENSG00000115524 | SF3B1 | splicing factor 3b subunit 1 [Source:HGNC Symbol;Acc:HGNC:10768] | 102 |
| 2 | DHX15 | ENSG00000109606 | DHX15 | DEAH-box helicase 15 [Source:HGNC Symbol;Acc:HGNC:2738] | 82 |
| 3 | SF3B3 | ENSG00000189091 | SF3B3 | splicing factor 3b subunit 3 [Source:HGNC Symbol;Acc:HGNC:10770] | 80 |
| 4 | SF3B2 | ENSG00000087365 | SF3B2 | splicing factor 3b subunit 2 [Source:HGNC Symbol;Acc:HGNC:10769] | 58 |
| 5 | CCAR1 | ENSG00000060339 | CCAR1 | cell division cycle and apoptosis regulator 1 [Source:HGNC Symbol;Acc:HGNC:24236] | 55 |
| 6 | U2SURP | ENSG00000163714 | U2SURP | U2 snRNP associated SURP domain containing [Source:HGNC Symbol;Acc:HGNC:30855] | 55 |
| 7 | SNRNP200 | ENSG00000144028 | SNRNP200 | small nuclear ribonucleoprotein U5 subunit 200 [Source:HGNC Symbol;Acc:HGNC:30859] | 50 |
| 8 | HNRNPL | ENSG00000104824 | HNRNPL | heterogeneous nuclear ribonucleoprotein L [Source:HGNC Symbol;Acc:HGNC:5045] | 48 |
| 9 | DHX9 | ENSG00000135829 | DHX9 | DExH-box helicase 9 [Source:HGNC Symbol;Acc:HGNC:2750] | 48 |
| 10 | HNRNPUL1 | ENSG00000105323 | HNRNPUL1 | heterogeneous nuclear ribonucleoprotein U like 1 [Source:HGNC Symbol;Acc:HGNC:17011] | 47 |
| 11 | PUF60 | ENSG00000179950 | PUF60 | poly(U) binding splicing factor 60 [Source:HGNC Symbol;Acc:HGNC:17042] | 47 |
| 12 | HNRNPA2B1 | ENSG00000122566 | HNRNPA2B1 | heterogeneous nuclear ribonucleoprotein A2/B1 [Source:HGNC Symbol;Acc:HGNC:5033] | 46 |
| 13 | PTBP1 | ENSG00000011304 | PTBP1 | polypyrimidine tract binding protein 1 [Source:HGNC Symbol;Acc:HGNC:9583] | 46 |
| 14 | SF3A3 | ENSG00000183431 | SF3A3 | splicing factor 3a subunit 3 [Source:HGNC Symbol;Acc:HGNC:10767] | 44 |
| 15 | SF3A1 | ENSG00000099995 | SF3A1 | splicing factor 3a subunit 1 [Source:HGNC Symbol;Acc:HGNC:10765] | 44 |
| 16 | DDX5 | ENSG00000108654 | DDX5 | DEAD-box helicase 5 [Source:HGNC Symbol;Acc:HGNC:2746] | 43 |
| 17 | U2AF2 | ENSG00000063244 | U2AF2 | U2 small nuclear RNA auxiliary factor 2 [Source:HGNC Symbol;Acc:HGNC:23156] | 41 |
| 18 | KHSRP | ENSG00000088247 | KHSRP | KH-type splicing regulatory protein [Source:HGNC Symbol;Acc:HGNC:6316] | 41 |
| 19 | PRPF8 | ENSG00000174231 | PRPF8 | pre-mRNA processing factor 8 [Source:HGNC Symbol;Acc:HGNC:17340] | 37 |
| 20 | RBM17 | ENSG00000134453 | RBM17 | RNA binding motif protein 17 [Source:HGNC Symbol;Acc:HGNC:16944] | 35 |
| 21 | ELAVL1 | ENSG00000066044 | ELAVL1 | ELAV like RNA binding protein 1 [Source:HGNC Symbol;Acc:HGNC:3312] | 35 |
| 22 | CPSF1 | ENSG00000071894 | CPSF1 | cleavage and polyadenylation specific factor 1 [Source:HGNC Symbol;Acc:HGNC:2324] | 35 |
| 23 | HNRNPA3 | ENSG00000170144 | HNRNPA3 | heterogeneous nuclear ribonucleoprotein A3 [Source:HGNC Symbol;Acc:HGNC:24941] | 33 |
| 24 | SF1 | ENSG00000168066 | SF1 | splicing factor 1 [Source:HGNC Symbol;Acc:HGNC:12950] | 33 |
| 25 | FUBP1 | ENSG00000162613 | FUBP1 | far upstream element binding protein 1 [Source:HGNC Symbol;Acc:HGNC:4004] | 32 |
| 26 | CHERP | ENSG00000085872 | CHERP | calcium homeostasis endoplasmic reticulum protein [Source:HGNC Symbol;Acc:HGNC:16930] | 31 |
| 27 | DAZAP1 | ENSG00000071626 | DAZAP1 | DAZ associated protein 1 [Source:HGNC Symbol;Acc:HGNC:2683] | 29 |
| 28 | RBM39 | ENSG00000131051 | RBM39 | RNA binding motif protein 39 [Source:HGNC Symbol;Acc:HGNC:15923] | 28 |
| 29 | MATR3 | ENSG00000015479 | MATR3 | matrin 3 [Source:HGNC Symbol;Acc:HGNC:6912] | 28 |
| 30 | RBM25 | ENSG00000119707 | RBM25 | RNA binding motif protein 25 [Source:HGNC Symbol;Acc:HGNC:23244] | 28 |
| 31 | EIF4A3 | ENSG00000141543 | EIF4A3 | eukaryotic translation initiation factor 4A3 [Source:HGNC Symbol;Acc:HGNC:18683] | 28 |
| 32 | CPSF2 | ENSG00000165934 | CPSF2 | cleavage and polyadenylation specific factor 2 [Source:HGNC Symbol;Acc:HGNC:2325] | 28 |
| 33 | HNRNPK | ENSG00000165119 | HNRNPK | heterogeneous nuclear ribonucleoprotein K [Source:HGNC Symbol;Acc:HGNC:5044] | 27 |
| 34 | RBM10 | ENSG00000182872 | RBM10 | RNA binding motif protein 10 [Source:HGNC Symbol;Acc:HGNC:9896] | 26 |
| 35 | EFTUD2 | ENSG00000108883 | EFTUD2 | elongation factor Tu GTP binding domain containing 2 [Source:HGNC Symbol;Acc:HGNC:30858] | 25 |
| 36 | HNRNPM | ENSG00000099783 | HNRNPM | heterogeneous nuclear ribonucleoprotein M [Source:HGNC Symbol;Acc:HGNC:5046] | 24 |
| 37 | SNRPA1 | ENSG00000131876 | SNRPA1 | small nuclear ribonucleoprotein polypeptide A' [Source:HGNC Symbol;Acc:HGNC:11152] | 24 |
| 38 | HNRNPH1 | ENSG00000169045 | HNRNPH1 | heterogeneous nuclear ribonucleoprotein H1 [Source:HGNC Symbol;Acc:HGNC:5041] | 24 |
| 39 | PCBP1 | ENSG00000169564 | PCBP1 | poly(rC) binding protein 1 [Source:HGNC Symbol;Acc:HGNC:8647] | 24 |
| 40 | SNRNP70 | ENSG00000104852 | SNRNP70 | small nuclear ribonucleoprotein U1 subunit 70 [Source:HGNC Symbol;Acc:HGNC:11150] | 23 |
| 41 | HNRNPR | ENSG00000125944 | HNRNPR | heterogeneous nuclear ribonucleoprotein R [Source:HGNC Symbol;Acc:HGNC:5047] | 21 |
| 42 | SF3B6 | ENSG00000115128 | SF3B6 | splicing factor 3b subunit 6 [Source:HGNC Symbol;Acc:HGNC:30096] | 20 |
| 43 | SRSF1 | ENSG00000136450 | SRSF1 | serine and arginine rich splicing factor 1 [Source:HGNC Symbol;Acc:HGNC:10780] | 19 |
| 44 | FUS | ENSG00000089280 | FUS | FUS RNA binding protein [Source:HGNC Symbol;Acc:HGNC:4010] | 18 |
| 45 | ALYREF | ENSG00000183684 | ALYREF | Aly/REF export factor [Source:HGNC Symbol;Acc:HGNC:19071] | 17 |
| 46 | SNRPD3 | ENSG00000100028 | SNRPD3 | small nuclear ribonucleoprotein D3 polypeptide [Source:HGNC Symbol;Acc:HGNC:11160] | 17 |
| 47 | SNRPD3 | ENSG00000286070 | None | novel protein | 17 |
| 48 | PRPF40A | ENSG00000196504 | PRPF40A | pre-mRNA processing factor 40 homolog A [Source:HGNC Symbol;Acc:HGNC:16463] | 17 |
| 49 | SF3A2 | ENSG00000104897 | SF3A2 | splicing factor 3a subunit 2 [Source:HGNC Symbol;Acc:HGNC:10766] | 17 |
| 50 | THRAP3 | ENSG00000054118 | THRAP3 | thyroid hormone receptor associated protein 3 [Source:HGNC Symbol;Acc:HGNC:22964] | 17 |
| 51 | CPSF7 | ENSG00000149532 | CPSF7 | cleavage and polyadenylation specific factor 7 [Source:HGNC Symbol;Acc:HGNC:30098] | 16 |
| 52 | HNRNPDL | ENSG00000152795 | HNRNPDL | heterogeneous nuclear ribonucleoprotein D like [Source:HGNC Symbol;Acc:HGNC:5037] | 16 |
| 53 | HNRNPA0 | ENSG00000177733 | HNRNPA0 | heterogeneous nuclear ribonucleoprotein A0 [Source:HGNC Symbol;Acc:HGNC:5030] | 16 |
| 54 | DDX3X | ENSG00000215301 | DDX3X | DEAD-box helicase 3 X-linked [Source:HGNC Symbol;Acc:HGNC:2745] | 15 |
| 55 | ACIN1 | ENSG00000100813 | ACIN1 | apoptotic chromatin condensation inducer 1 [Source:HGNC Symbol;Acc:HGNC:17066] | 15 |
| 56 | SF3B4 | ENSG00000143368 | SF3B4 | splicing factor 3b subunit 4 [Source:HGNC Symbol;Acc:HGNC:10771] | 15 |
| 57 | SYNCRIP | ENSG00000135316 | SYNCRIP | synaptotagmin binding cytoplasmic RNA interacting protein [Source:HGNC Symbol;Acc:HGNC:16918] | 15 |
| 58 | RBMX | ENSG00000147274 | RBMX | RNA binding motif protein X-linked [Source:HGNC Symbol;Acc:HGNC:9910] | 15 |
| 59 | SRSF7 | ENSG00000115875 | SRSF7 | serine and arginine rich splicing factor 7 [Source:HGNC Symbol;Acc:HGNC:10789] | 14 |
| 60 | NUDT21 | ENSG00000167005 | NUDT21 | nudix hydrolase 21 [Source:HGNC Symbol;Acc:HGNC:13870] | 14 |
| 61 | WDR33 | ENSG00000136709 | WDR33 | WD repeat domain 33 [Source:HGNC Symbol;Acc:HGNC:25651] | 13 |
| 62 | RPS7 | ENSG00000171863 | RPS7 | ribosomal protein S7 [Source:HGNC Symbol;Acc:HGNC:10440] | 13 |
| 63 | FIP1L1 | ENSG00000145216 | FIP1L1 | factor interacting with PAPOLA and CPSF1 [Source:HGNC Symbol;Acc:HGNC:19124] | 13 |
| 64 | HNRNPAB | ENSG00000197451 | HNRNPAB | heterogeneous nuclear ribonucleoprotein A/B [Source:HGNC Symbol;Acc:HGNC:5034] | 12 |
| 65 | HNRNPD | ENSG00000138668 | HNRNPD | heterogeneous nuclear ribonucleoprotein D [Source:HGNC Symbol;Acc:HGNC:5036] | 12 |
| 66 | SRSF11 | ENSG00000116754 | SRSF11 | serine and arginine rich splicing factor 11 [Source:HGNC Symbol;Acc:HGNC:10782] | 12 |
| 67 | CSTF3 | ENSG00000176102 | CSTF3 | cleavage stimulation factor subunit 3 [Source:HGNC Symbol;Acc:HGNC:2485] | 12 |
| 68 | RPL31 | ENSG00000071082 | RPL31 | ribosomal protein L31 [Source:HGNC Symbol;Acc:HGNC:10334] | 11 |
| 69 | DDX17 | ENSG00000100201 | DDX17 | DEAD-box helicase 17 [Source:HGNC Symbol;Acc:HGNC:2740] | 11 |
| 70 | SNRPB2 | ENSG00000125870 | SNRPB2 | small nuclear ribonucleoprotein polypeptide B2 [Source:HGNC Symbol;Acc:HGNC:11155] | 12 |
| 71 | TIAL1 | ENSG00000151923 | TIAL1 | TIA1 cytotoxic granule associated RNA binding protein like 1 [Source:HGNC Symbol;Acc:HGNC:11804] | 11 |
| 72 | HNRNPU | ENSG00000153187 | HNRNPU | heterogeneous nuclear ribonucleoprotein U [Source:HGNC Symbol;Acc:HGNC:5048] | 11 |
| 73 | SNRPD1 | ENSG00000167088 | SNRPD1 | small nuclear ribonucleoprotein D1 polypeptide [Source:HGNC Symbol;Acc:HGNC:11158] | 11 |
| 74 | RPS4X | ENSG00000198034 | RPS4X | ribosomal protein S4 X-linked [Source:HGNC Symbol;Acc:HGNC:10424] | 10 |
| 75 | DHX36 | ENSG00000174953 | DHX36 | DEAH-box helicase 36 [Source:HGNC Symbol;Acc:HGNC:14410] | 10 |
| 76 | ILF2 | ENSG00000143621 | ILF2 | interleukin enhancer binding factor 2 [Source:HGNC Symbol;Acc:HGNC:6037] | 10 |
| 77 | EWSR1 | ENSG00000182944 | EWSR1 | EWS RNA binding protein 1 [Source:HGNC Symbol;Acc:HGNC:3508] | 9 |
| 78 | LUC7L3 | ENSG00000108848 | LUC7L3 | LUC7 like 3 pre-mRNA splicing factor [Source:HGNC Symbol;Acc:HGNC:24309] | 9 |
| 79 | KHDRBS1 | ENSG00000121774 | KHDRBS1 | KH RNA binding domain containing, signal transduction associated 1 [Source:HGNC Symbol;Acc:HGNC:18116] | 9 |
| 80 | PNN | ENSG00000100941 | PNN | pinin, desmosome associated protein [Source:HGNC Symbol;Acc:HGNC:9162] | 9 |
| 81 | CPSF6 | ENSG00000111605 | CPSF6 | cleavage and polyadenylation specific factor 6 [Source:HGNC Symbol;Acc:HGNC:13871] | 9 |
| 82 | SAP18 | ENSG00000150459 | SAP18 | Sin3A associated protein 18 [Source:HGNC Symbol;Acc:HGNC:10530] | 9 |
| 83 | SRSF3 | ENSG00000112081 | SRSF3 | serine and arginine rich splicing factor 3 [Source:HGNC Symbol;Acc:HGNC:10785] | 9 |
| 84 | THOC2 | ENSG00000125676 | THOC2 | THO complex 2 [Source:HGNC Symbol;Acc:HGNC:19073] | 9 |
| 85 | CPSF3 | ENSG00000119203 | CPSF3 | cleavage and polyadenylation specific factor 3 [Source:HGNC Symbol;Acc:HGNC:2326] | 9 |
| 86 | POLDIP3 | ENSG00000100227 | POLDIP3 | DNA polymerase delta interacting protein 3 [Source:HGNC Symbol;Acc:HGNC:23782] | 9 |
| 87 | SFPQ | ENSG00000116560 | SFPQ | splicing factor proline and glutamine rich [Source:HGNC Symbol;Acc:HGNC:10774] | 8 |
| 88 | SRRM2 | ENSG00000167978 | SRRM2 | serine/arginine repetitive matrix 2 [Source:HGNC Symbol;Acc:HGNC:16639] | 8 |
| 89 | IGF2BP1 | ENSG00000159217 | IGF2BP1 | insulin like growth factor 2 mRNA binding protein 1 [Source:HGNC Symbol;Acc:HGNC:28866] | 8 |
| 90 | PRPF4B | ENSG00000112739 | PRPF4B | pre-mRNA processing factor 4B [Source:HGNC Symbol;Acc:HGNC:17346] | 8 |
| 91 | NONO | ENSG00000147140 | NONO | non-POU domain containing octamer binding [Source:HGNC Symbol;Acc:HGNC:7871] | 7 |
| 92 | CIRBP | ENSG00000099622 | CIRBP | cold inducible RNA binding protein [Source:HGNC Symbol;Acc:HGNC:1982] | 7 |
| 93 | SMU1 | ENSG00000122692 | SMU1 | SMU1 DNA replication regulator and spliceosomal factor [Source:HGNC Symbol;Acc:HGNC:18247] | 7 |
| 94 | SRRM1 | ENSG00000133226 | SRRM1 | serine and arginine repetitive matrix 1 [Source:HGNC Symbol;Acc:HGNC:16638] | 6 |
| 95 | SNRPD2 | ENSG00000125743 | SNRPD2 | small nuclear ribonucleoprotein D2 polypeptide [Source:HGNC Symbol;Acc:HGNC:11159] | 6 |
| 96 | TRA2B | ENSG00000136527 | TRA2B | transformer 2 beta homolog [Source:HGNC Symbol;Acc:HGNC:10781] | 6 |
| 97 | HNRNPC | ENSG00000092199 | HNRNPC | heterogeneous nuclear ribonucleoprotein C [Source:HGNC Symbol;Acc:HGNC:5035] | 5 |
| 98 | BUB3 | ENSG00000154473 | BUB3 | BUB3 mitotic checkpoint protein [Source:HGNC Symbol;Acc:HGNC:1151] | 5 |
| 99 | CPSF4 | ENSG00000160917 | CPSF4 | cleavage and polyadenylation specific factor 4 [Source:HGNC Symbol;Acc:HGNC:2327] | 5 |
| 100 | SNRPF | ENSG00000139343 | SNRPF | small nuclear ribonucleoprotein polypeptide F [Source:HGNC Symbol;Acc:HGNC:11162] | 5 |
| 101 | SNRPA | ENSG00000077312 | SNRPA | small nuclear ribonucleoprotein polypeptide A [Source:HGNC Symbol;Acc:HGNC:11151] | 5 |
| 102 | RPS11 | ENSG00000142534 | RPS11 | ribosomal protein S11 [Source:HGNC Symbol;Acc:HGNC:10384] | 5 |
| 103 | RPS15A | ENSG00000134419 | RPS15A | ribosomal protein S15a [Source:HGNC Symbol;Acc:HGNC:10389] | 5 |
| 104 | CHTOP | ENSG00000160679 | CHTOP | chromatin target of PRMT1 [Source:HGNC Symbol;Acc:HGNC:24511] | 5 |

hNEAT1 9.2-9.8 kb RNA pulldown & MS spectrometry analysis

| | initial_alias | converted_alias | name | description | unique peptide # |
|---|---|---|---|---|---|
| 1 | SF3B1 | ENSG00000115524 | SF3B1 | splicing factor 3b subunit 1 [Source:HGNC Symbol;Acc:HGNC:10768] | 113 |
| 2 | DHX15 | ENSG00000109606 | DHX15 | DEAH-box helicase 15 [Source:HGNC Symbol;Acc:HGNC:2738] | 101 |
| 3 | SF3B3 | ENSG00000189091 | SF3B3 | splicing factor 3b subunit 3 [Source:HGNC Symbol;Acc:HGNC:10770] | 82 |
| 4 | CCAR1 | ENSG00000060339 | CCAR1 | cell division cycle and apoptosis regulator 1 [Source:HGNC Symbol;Acc:HGNC:24236] | 73 |
| 5 | U2SURP | ENSG00000163714 | U2SURP | U2 snRNP associated SURP domain containing [Source:HGNC Symbol;Acc:HGNC:30855] | 66 |
| 6 | SF3B2 | ENSG00000087365 | SF3B2 | splicing factor 3b subunit 2 [Source:HGNC Symbol;Acc:HGNC:10769] | 56 |
| 7 | HNRNPL | ENSG00000104824 | HNRNPL | heterogeneous nuclear ribonucleoprotein L [Source:HGNC Symbol;Acc:HGNC:5045] | 52 |
| 8 | HNRNPUL1 | ENSG00000105323 | HNRNPUL1 | heterogeneous nuclear ribonucleoprotein U like 1 [Source:HGNC Symbol;Acc:HGNC:17011] | 52 |
| 9 | PTBP1 | ENSG00000011304 | PTBP1 | polypyrimidine tract binding protein 1 [Source:HGNC Symbol;Acc:HGNC:9583] | 51 |
| 10 | SF3A3 | ENSG00000183431 | SF3A3 | splicing factor 3a subunit 3 [Source:HGNC Symbol;Acc:HGNC:10767] | 49 |
| 11 | KHSRP | ENSG00000088247 | KHSRP | KH-type splicing regulatory protein [Source:HGNC Symbol;Acc:HGNC:6316] | 49 |
| 12 | PUF60 | ENSG00000179950 | PUF60 | poly(U) binding splicing factor 60 [Source:HGNC Symbol;Acc:HGNC:17042] | 48 |
| 13 | SF3A1 | ENSG00000099995 | SF3A1 | splicing factor 3a subunit 1 [Source:HGNC Symbol;Acc:HGNC:10765] | 47 |
| 14 | CHERP | ENSG00000085872 | CHERP | calcium homeostasis endoplasmic reticulum protein [Source:HGNC Symbol;Acc:HGNC:16930] | 43 |
| 15 | HNRNPA2B1 | ENSG00000122566 | HNRNPA2B1 | heterogeneous nuclear ribonucleoprotein A2/B1 [Source:HGNC Symbol;Acc:HGNC:5033] | 41 |
| 16 | U2AF2 | ENSG00000063244 | U2AF2 | U2 small nuclear RNA auxiliary factor 2 [Source:HGNC Symbol;Acc:HGNC:23156] | 41 |
| 17 | SNRNP200 | ENSG00000144028 | SNRNP200 | small nuclear ribonucleoprotein U5 subunit 200 [Source:HGNC Symbol;Acc:HGNC:30859] | 41 |
| 18 | FUBP1 | ENSG00000162613 | FUBP1 | far upstream element binding protein 1 [Source:HGNC Symbol;Acc:HGNC:4004] | 39 |
| 19 | DDX5 | ENSG00000108654 | DDX5 | DEAD-box helicase 5 [Source:HGNC Symbol;Acc:HGNC:2746] | 36 |
| 20 | SF1 | ENSG00000168066 | SF1 | splicing factor 1 [Source:HGNC Symbol;Acc:HGNC:12950] | 36 |
| 21 | RBM17 | ENSG00000134453 | RBM17 | RNA binding motif protein 17 [Source:HGNC Symbol;Acc:HGNC:16944] | 35 |
| 22 | HNRNPR | ENSG00000125944 | HNRNPR | heterogeneous nuclear ribonucleoprotein R [Source:HGNC Symbol;Acc:HGNC:5047] | 34 |
| 23 | SRSF1 | ENSG00000136450 | SRSF1 | serine and arginine rich splicing factor 1 [Source:HGNC Symbol;Acc:HGNC:10780] | 34 |
| 24 | PRPF8 | ENSG00000174231 | PRPF8 | pre-mRNA processing factor 8 [Source:HGNC Symbol;Acc:HGNC:17340] | 32 |
| 25 | RBM25 | ENSG00000119707 | RBM25 | RNA binding motif protein 25 [Source:HGNC Symbol;Acc:HGNC:23244] | 32 |
| 26 | DHX9 | ENSG00000135829 | DHX9 | DExH-box helicase 9 [Source:HGNC Symbol;Acc:HGNC:2750] | 31 |
| 27 | MATR3 | ENSG00000015479 | MATR3 | matrin 3 [Source:HGNC Symbol;Acc:HGNC:6912] | 30 |
| 28 | DAZAP1 | ENSG00000071626 | DAZAP1 | DAZ associated protein 1 [Source:HGNC Symbol;Acc:HGNC:2683] | 30 |
| 29 | EFTUD2 | ENSG00000108883 | EFTUD2 | elongation factor Tu GTP binding domain containing 2 [Source:HGNC Symbol;Acc:HGNC:30858] | 29 |
| 30 | RBM39 | ENSG00000131051 | RBM39 | RNA binding motif protein 39 [Source:HGNC Symbol;Acc:HGNC:15923] | 26 |
| 31 | ELAVL1 | ENSG00000066044 | ELAVL1 | ELAV like RNA binding protein 1 [Source:HGNC Symbol;Acc:HGNC:3312] | 26 |
| 32 | RBM10 | ENSG00000182872 | RBM10 | RNA binding motif protein 10 [Source:HGNC Symbol;Acc:HGNC:9896] | 26 |
| 33 | HNRNPM | ENSG00000099783 | HNRNPM | heterogeneous nuclear ribonucleoprotein M [Source:HGNC Symbol;Acc:HGNC:5046] | 24 |
| 34 | SNRPA1 | ENSG00000131876 | SNRPA1 | small nuclear ribonucleoprotein polypeptide A' [Source:HGNC Symbol;Acc:HGNC:11152] | 24 |
| 35 | SNRNP70 | ENSG00000104852 | SNRNP70 | small nuclear ribonucleoprotein U1 subunit 70 [Source:HGNC Symbol;Acc:HGNC:11150] | 24 |
| 36 | EIF4A3 | ENSG00000141543 | EIF4A3 | eukaryotic translation initiation factor 4A3 [Source:HGNC Symbol;Acc:HGNC:18683] | 23 |
| 37 | HNRNPK | ENSG00000165119 | HNRNPK | heterogeneous nuclear ribonucleoprotein K [Source:HGNC Symbol;Acc:HGNC:5044] | 22 |
| 38 | HNRNPA3 | ENSG00000170144 | HNRNPA3 | heterogeneous nuclear ribonucleoprotein A3 [Source:HGNC Symbol;Acc:HGNC:24941] | 21 |
| 39 | HNRNPH1 | ENSG00000169045 | HNRNPH1 | heterogeneous nuclear ribonucleoprotein H1 [Source:HGNC Symbol;Acc:HGNC:5041] | 20 |
| 40 | HNRNPDL | ENSG00000152795 | HNRNPDL | heterogeneous nuclear ribonucleoprotein D like [Source:HGNC Symbol;Acc:HGNC:5037] | 20 |
| 41 | ACIN1 | ENSG00000100813 | ACIN1 | apoptotic chromatin condensation inducer 1 [Source:HGNC Symbol;Acc:HGNC:17066] | 19 |
| 42 | PCBP1 | ENSG00000169564 | PCBP1 | poly(rC) binding protein 1 [Source:HGNC Symbol;Acc:HGNC:8647] | 18 |
| 43 | LUC7L3 | ENSG00000108848 | LUC7L3 | LUC7 like 3 pre-mRNA splicing factor [Source:HGNC Symbol;Acc:HGNC:24309] | 18 |
| 44 | FUS | ENSG00000089280 | FUS | FUS RNA binding protein [Source:HGNC Symbol;Acc:HGNC:4010] | 17 |
| 45 | SF3B6 | ENSG00000115128 | SF3B6 | splicing factor 3b subunit 6 [Source:HGNC Symbol;Acc:HGNC:30096] | 17 |
| 46 | PRPF40A | ENSG00000196504 | PRPF40A | pre-mRNA processing factor 40 homolog A [Source:HGNC Symbol;Acc:HGNC:16463] | 17 |
| 47 | SF3A2 | ENSG00000104897 | SF3A2 | splicing factor 3a subunit 2 [Source:HGNC Symbol;Acc:HGNC:10766] | 17 |
| 48 | HNRNPA0 | ENSG00000177733 | HNRNPA0 | heterogeneous nuclear ribonucleoprotein A0 [Source:HGNC Symbol;Acc:HGNC:5030] | 17 |
| 49 | SF3B4 | ENSG00000143368 | SF3B4 | splicing factor 3b subunit 4 [Source:HGNC Symbol;Acc:HGNC:10771] | 17 |
| 50 | SYNCRIP | ENSG00000135316 | SYNCRIP | synaptotagmin binding cytoplasmic RNA interacting protein [Source:HGNC Symbol;Acc:HGNC:16918] | 17 |
| 51 | RBMX | ENSG00000147274 | RBMX | RNA binding motif protein X-linked [Source:HGNC Symbol;Acc:HGNC:9910] | 16 |
| 52 | CPSF1 | ENSG00000071894 | CPSF1 | cleavage and polyadenylation specific factor 1 [Source:HGNC Symbol;Acc:HGNC:2324] | 15 |
| 53 | DDX3X | ENSG00000215301 | DDX3X | DEAD-box helicase 3 X-linked [Source:HGNC Symbol;Acc:HGNC:2745] | 15 |
| 54 | DDX42 | ENSG00000198231 | DDX42 | DEAD-box helicase 42 [Source:HGNC Symbol;Acc:HGNC:18676] | 15 |
| 55 | ALYREF | ENSG00000183684 | ALYREF | Aly/REF export factor [Source:HGNC Symbol;Acc:HGNC:19071] | 14 |
| 56 | SRRM2 | ENSG00000167978 | SRRM2 | serine/arginine repetitive matrix 2 [Source:HGNC Symbol;Acc:HGNC:16639] | 14 |
| 57 | SNRPD3 | ENSG00000100028 | SNRPD3 | small nuclear ribonucleoprotein D3 polypeptide [Source:HGNC Symbol;Acc:HGNC:11160] | 13 |
| 58 | THRAP3 | ENSG00000054118 | THRAP3 | thyroid hormone receptor associated protein 3 [Source:HGNC Symbol;Acc:HGNC:22964] | 13 |
| 59 | HNRNPAB | ENSG00000197451 | HNRNPAB | heterogeneous nuclear ribonucleoprotein A/B [Source:HGNC Symbol;Acc:HGNC:5034] | 13 |
| 60 | HNRNPD | ENSG00000138668 | HNRNPD | heterogeneous nuclear ribonucleoprotein D [Source:HGNC Symbol;Acc:HGNC:5036] | 13 |
| 61 | SRSF11 | ENSG00000116754 | SRSF11 | serine and arginine rich splicing factor 11 [Source:HGNC Symbol;Acc:HGNC:10782] | 13 |
| 62 | BUB3 | ENSG00000154473 | BUB3 | BUB3 mitotic checkpoint protein [Source:HGNC Symbol;Acc:HGNC:1151] | 13 |
| 63 | SRSF7 | ENSG00000115875 | SRSF7 | serine and arginine rich splicing factor 7 [Source:HGNC Symbol;Acc:HGNC:10789] | 12 |
| 64 | EWSR1 | ENSG00000182944 | EWSR1 | EWS RNA binding protein 1 [Source:HGNC Symbol;Acc:HGNC:3508] | 12 |
| 65 | SNRPB2 | ENSG00000125870 | SNRPB2 | small nuclear ribonucleoprotein polypeptide B2 [Source:HGNC Symbol;Acc:HGNC:11155] | 12 |
| 66 | NUDT21 | ENSG00000167005 | NUDT21 | nudix hydrolase 21 [Source:HGNC Symbol;Acc:HGNC:13870] | 11 |
| 67 | RPS4X | ENSG00000198034 | RPS4X | ribosomal protein S4 X-linked [Source:HGNC Symbol;Acc:HGNC:10424] | 11 |
| 68 | CPSF7 | ENSG00000149532 | CPSF7 | cleavage and polyadenylation specific factor 7 [Source:HGNC Symbol;Acc:HGNC:30098] | 10 |
| 69 | RPS7 | ENSG00000171863 | RPS7 | ribosomal protein S7 [Source:HGNC Symbol;Acc:HGNC:10440] | 10 |
| 70 | SNRPD1 | ENSG00000167088 | SNRPD1 | small nuclear ribonucleoprotein D1 polypeptide [Source:HGNC Symbol;Acc:HGNC:11158] | 10 |
| 71 | KHDRBS1 | ENSG00000121774 | KHDRBS1 | KH RNA binding domain containing, signal transduction associated 1 [Source:HGNC Symbol;Acc:HGNC:18116] | 10 |
| 72 | RPL31 | ENSG00000071082 | RPL31 | ribosomal protein L31 [Source:HGNC Symbol;Acc:HGNC:10334] | 9 |
| 73 | FIP1L1 | ENSG00000145216 | FIP1L1 | factor interacting with PAPOLA and CPSF1 [Source:HGNC Symbol;Acc:HGNC:19124] | 9 |
| 74 | DDX17 | ENSG00000100201 | DDX17 | DEAD-box helicase 17 [Source:HGNC Symbol;Acc:HGNC:2740] | 9 |
| 75 | TIAL1 | ENSG00000151923 | TIAL1 | TIA1 cytotoxic granule associated RNA binding protein like 1 [Source:HGNC Symbol;Acc:HGNC:11804] | 9 |
| 76 | CPSF2 | ENSG00000165934 | CPSF2 | cleavage and polyadenylation specific factor 2 [Source:HGNC Symbol;Acc:HGNC:2325] | 8 |
| 77 | PNN | ENSG00000100941 | PNN | pinin, desmosome associated protein [Source:HGNC Symbol;Acc:HGNC:9162] | 8 |
| 78 | CPSF6 | ENSG00000111605 | CPSF6 | cleavage and polyadenylation specific factor 6 [Source:HGNC Symbol;Acc:HGNC:13871] | 8 |
| 79 | SAP18 | ENSG00000150459 | SAP18 | Sin3A associated protein 18 [Source:HGNC Symbol;Acc:HGNC:10530] | 8 |
| 80 | SRSF3 | ENSG00000112081 | SRSF3 | serine and arginine rich splicing factor 3 [Source:HGNC Symbol;Acc:HGNC:10785] | 8 |
| 81 | THOC2 | ENSG00000125676 | THOC2 | THO complex 2 [Source:HGNC Symbol;Acc:HGNC:19073] | 8 |
| 82 | IGF2BP1 | ENSG00000159217 | IGF2BP1 | insulin like growth factor 2 mRNA binding protein 1 [Source:HGNC Symbol;Acc:HGNC:28866] | 8 |
| 83 | SREK1 | ENSG00000153914 | SREK1 | splicing regulatory glutamic acid and lysine rich protein 1 [Source:HGNC Symbol;Acc:HGNC:17882] | 8 |
| 84 | PRPF4B | ENSG00000112739 | PRPF4B | pre-mRNA processing factor 4B [Source:HGNC Symbol;Acc:HGNC:17346] | 7 |
| 85 | CIRBP | ENSG00000099622 | CIRBP | cold inducible RNA binding protein [Source:HGNC Symbol;Acc:HGNC:1982] | 7 |
| 86 | SRRM1 | ENSG00000133226 | SRRM1 | serine and arginine repetitive matrix 1 [Source:HGNC Symbol;Acc:HGNC:16638] | 7 |
| 87 | SNRPD2 | ENSG00000125743 | SNRPD2 | small nuclear ribonucleoprotein D2 polypeptide [Source:HGNC Symbol;Acc:HGNC:11159] | 7 |
| 88 | TRIR | ENSG00000123144 | TRIR | telomerase RNA component interacting RNase [Source:HGNC Symbol;Acc:HGNC:28424] | 7 |
| 89 | CMTR1 | ENSG00000137200 | CMTR1 | cap methyltransferase 1 [Source:HGNC Symbol;Acc:HGNC:21077] | 7 |
| 90 | NONO | ENSG00000147140 | NONO | non-POU domain containing octamer binding [Source:HGNC Symbol;Acc:HGNC:7871] | 6 |
| 91 | DHX36 | ENSG00000174953 | DHX36 | DEAH-box helicase 36 [Source:HGNC Symbol;Acc:HGNC:14410] | 6 |
| 92 | TRA2B | ENSG00000136527 | TRA2B | transformer 2 beta homolog [Source:HGNC Symbol;Acc:HGNC:10781] | 6 |
| 93 | PRPF19 | ENSG00000110107 | PRPF19 | pre-mRNA processing factor 19 [Source:HGNC Symbol;Acc:HGNC:17896] | 6 |
| 94 | HNRNPF | ENSG00000169813 | HNRNPF | heterogeneous nuclear ribonucleoprotein F [Source:HGNC Symbol;Acc:HGNC:5039] | 6 |
| 95 | GTPBP4 | ENSG00000107937 | GTPBP4 | GTP binding protein 4 [Source:HGNC Symbol;Acc:HGNC:21535] | 6 |
| 96 | SFPQ | ENSG00000116560 | SFPQ | splicing factor proline and glutamine rich [Source:HGNC Symbol;Acc:HGNC:10774] | 5 |
| 97 | CSTF3 | ENSG00000176102 | CSTF3 | cleavage stimulation factor subunit 3 [Source:HGNC Symbol;Acc:HGNC:2485] | 5 |
| 98 | ERH | ENSG00000100632 | ERH | ERH mRNA splicing and mitosis factor [Source:HGNC Symbol;Acc:HGNC:3447] | 5 |
| 99 | SMU1 | ENSG00000122692 | SMU1 | SMU1 DNA replication regulator and spliceosomal factor [Source:HGNC Symbol;Acc:HGNC:18247] | 5 |
| 100 | HNRNPC | ENSG00000092199 | HNRNPC | heterogeneous nuclear ribonucleoprotein C [Source:HGNC Symbol;Acc:HGNC:5035] | 5 |
| 101 | DDX23 | ENSG00000174243 | DDX23 | DEAD-box helicase 23 [Source:HGNC Symbol;Acc:HGNC:17347] | 5 |
| 102 | SRSF2 | ENSG00000161547 | SRSF2 | serine and arginine rich splicing factor 2 [Source:HGNC Symbol;Acc:HGNC:10783] | 5 |
