## Supplementary Table 2 for "Shell protein composition specified by NEAT1 domains dictates the formation of paraspeckles as distinct membraneless organelles"

| REAGENT or RESOURCE | SOURCE | IDENTIFIER |
| --- | --- | --- |
| Mouse monoclonal anti-SC35 | Sigma-Aldrich (Merck) | Cat#S4045 |
| Rabbit polyclonal anti-SON/DBP5 | Abcam | Cat#ab121759 |
| Mouse monoclonal anti-NONO (clone 3/p54nrb) | BD Biosciences | Cat#611279 |
| Mouse monoclonal anti-FUS (clone 4H11) | Santa Cruz Biotechnology | Cat#sc-47711 |
| Mouse monoclonal anti-SFPQ (PSF) (clone C23) | MBL | Cat#RN014MW |
| Rabbit polyclonal anti-HNRNPH1 | Bethyl laboratories | Cat#A300-511A |
| Rabbit polyclonal anti-BRG1 | Bethyl laboratories | Cat#A300-813A |
| Rabbit polyclonal anti-HNRNPF (N1N3) | GeneTex | Cat#IHC00087 |
| Rabbit polyclonal anti-PSPC1 | Naganuma et al., 2012 | N/A |
| Rabbit polyclonal anti-HNRNPH3 | Abcam | Cat#ab6663 |
| Rabbit polyclonal anti-RBM14 | Bethyl laboratories | Cat#300-331A |
| Rabbit polyclonal anti-TDP43 | Proteintech | Cat#10782-2-AP |
| Rabbit polyclonal anit-HNRNPA1 | MBL | Cat#RN014PW |
| Rabbit polyclonal anti-DDDDK-tag pAb | MBL | Cat#PM020 |
| Mouse monoclonal anti-FLAG (clone M2) | Sigma-Aldrich (Merck) | Cat#F3165 |
| Rabbit polyclonal anti-MS2 coat protiein | Sigma-Aldrich (Merck) | Cat#ABE76-I |
| Mouse monoclonal anti-HA (clone 16B12) | BioLegend | Cat#901513 |
| Mouse monoclonal anti-SF3B1 (clone B-3) | Santa Cruz Biotechnology | Cat#sc-514655 |
| Mouse monclonal anti-Coilin (Pdelta) | Abcam | Cat#ab11822 |
| Mouse monclonal anti-PML (PG-M3) | Santa Cruz Biotechnology | Cat#sc-966 |
| Mouse monclonal anti-NPAT | Transduction Laboratories | Cat#N91820 |
| Mouse monoclonal anti-GAPDH (clone 6C5) | Abcam | Cat#ab8245 |
| Rabbit polyclonal anti-GAPDH | Abcam | Cat#ab37168 |
| Goat anti-rabbit IgG, Cy2 conjugate | Abcam | Cat#ab6940 |
| Goat anti-mouse IgG, Cy3 conjugate | Merck | Cat#AP124C |
| Goat anti-Rabbit IgG (H+L) Cross-Adsorbed Secondary Antibody, Alexa Fluor 405 | Thermo Fisher Scientific | Cat#A-31556 |
| Goat anti-Mouse IgG (H+L), Superclonal™ Recombinant Secondary Antibody, Alexa Fluor 488 | Thermo Fisher Scientific | Cat#A-28175 |
| Goat anti-Rabbit IgG (H+L) Highly Cross-Adsorbed Secondary Antibody, Alexa Fluor 488 | Thermo Fisher Scientific | Cat#A-11034 |
| Goat anti-Mouse IgG (H+L) Highly Cross-Adsorbed Secondary Antibody, Alexa Fluor 568 | Thermo Fisher Scientific | Cat#A-11031 |
| Goat anti-Rabbit IgG (H+L) Highly Cross-Adsorbed Secondary Antibody, Alexa Fluor 568 | Thermo Fisher Scientific | Cat#A-11036 |
| Streptavidin, Alexa Fluor 568 conjugate | Thermo Fisher Scientific | Cat#S11226 |
| Streptavidin, Alexa Fluor 647 conjugate | Thermo Fisher Scientific | Cat#S32359 |

| Bacterial and Virus Strains | SOURCE | IDENTIFIER |
| --- | --- | --- |
| DH5 $\alpha$ Competent Cells | Thermo Fisher Scientific | Cat#18265017 |
| Stbl3 Competent cells | Thermo Fisher Scientific | Cat#C737303 |

| Chemicals, Peptides, and Recombinant Proteins | SOURCE | IDENTIFIER |
| --- | --- | --- |
| Z-Leu-Leu-Leu-al (MG132) | Sigma-Aldrich | Cat#C2211 |
| 5,6-Dichlorobenzimidazole 1-$\beta$-D-ribofuranoside (DRB) | Sigma-Aldrich | Cat#D1916 |
| Spliceostatin A | Yoshida Laboratory | N/A |
| Pladienolide B | Cayman Chemical | Cat#445493-23-2 |
| Blasticidin | InvivoGen | Cat#Ant-bl-1 |
| Puromycin | Sigma-Aldrich | Cat#P9620 |
| 2,2'-Thiodiethanol | Sigma-Aldrich | Cat#166782 |
| DABCO 33-LV | Sigma-Aldrich | Cat#290734 |
| VECTASHIELD Hard Set Mounting Medium with DAPI | Vector laboratories | Cat#H-1500 |
| ProLong™ Gold Antifade Mountant | Thermo Fisher Scientific | Cat#P36930 |
| TRI Reagent | Molecular Research Center | Cat#TR118 |
| cOmplete, EDTA free Protease Inhibitor Cocktail | Roche | Cat#5056489001 |
| TransIT LT-1 Reagent | Mirus | Cat#MIR2300 |
| Proteinase K | Roche | Cat#3115828001 |
| Blocking reagent | Roche | Cat#11096176001 |

| Critical Commercial Assays | SOURCE | IDENTIFIER |
| --- | --- | --- |
| High Capacity cDNA Reverse Transcription Kit | Thermo Fisher Scientific | Cat#4368813 |
| Stellaris FISH probe (Human NEAT1_5 with Quaser 570 Dye) | LGC Biosearch Technologies | Cat#SMF-2036-1 |
| Stellaris FISH probe (Human NEAT1_m with Quaser 570 Dye) | LGC Biosearch Technologies | Cat#SMF-2037-1 |
| T7 Endonuclease I | NEB | Cat#M0302L |
| Anti-FLAG M2 magnetic beads | Sigma-Aldrich | Cat#M8823 |
| Tamavidin2-REV Magnetic Beads | WAKO | Cat#136-18341 |
| KAPA SYBR Fast qPCR Kit | NIPPON Genetics | KK4610 |

| **HAP1 mutant cell lines** | **Indel regions in hNEAT1 (22743 nt)** | Reference | sgRNA#1 (5' to 3') | sgRNA#2 (5' to 3') | Cell lines modified |
|---|---|---|---|---|---|
| HAP1 mini-NEAT1 (Δ1-8kb/16.6-22.6kb) | 1034-8006/16613-22647 | Yamazaki et al., 2017 | | | |
| NEAT1 Δ16.6-22.6kb | 16613-22647 | Yamazaki et al., 2017 | | | |
| NEAT1 Δ1-8kb | 1034-8006 | Yamazaki et al., 2017 | | | |
| NEAT1 Δ4-8kb/16.6-22.6kb | 4054-8005/16613-22647 | This study | TACCACTTAGAACTCTAACC | GGATATCATATACTCACAAC | HAP1 NEAT1 Δ16.6-22.6k |
| NEAT1 Δ1-4kb/16.6-22.6kb | 1034-4053/16613-22647 | This study | AGAGGCCTTCCCGCTGAGGC | TACCACTTAGAACTCTAACC | HAP1 NEAT1 Δ16.6-22.6k |
| NEAT1 Δ20.2-22.6kb | 20157-22647 | This study | GGGAGGCGCGGAGCCGCCGC | TCAGATAACCATTCTAACTA | HAP1 WT |
| NEAT1 Δ1-8kb/20.2-22.6kb | 1034-8005/20157-22647 | This study | AGAGGCCTTCCCGCTGAGGC | GGATATCATATACTCACAAC | HAP1 NEAT1 Δ20.1-22.6k |
| NEAT1 Δ1-8kb/16.6-20.2kb | 1034-8006/16613-20156 | This study | TGACTTGATTTGGCTGTGCA | TCAGATAACCATTCTAACTA | HAP1 NEAT1 Δ1-8k |
| m9.8-16.6k (Δ1-9.8kb/16.6-22.6kb) | 1033-9830/16613-22647 | Yamazaki et al., 2017 | | | |

| **HAP1 knock-in mutant cell lines** | **Indel regions in hNEAT1 (22743 nt)** | Reference | sgRNA#1 (5' to 3') targeting template vector | sgRNA#2 (5' to 3') | Cell lines modified |
|---|---|---|---|---|---|
| mini-NEAT1/6xMS2BS@8.2kb | 8175-(6xMS2BS insertion)-8176 | This study | GCATCGTACGCGTACGTGTT | GGAGATTGGTTGTACAACAA | HAP1 mini-NEAT1 |
| mini-NEAT1/6xMS2BS@16.2kb | 16204-(6xMS2BS insertion)-16205 | This study | GCATCGTACGCGTACGTGTT | GGAACCCTCTTACTTACTGG | HAP1 mini-NEAT1 |
| NEAT1/6xMS2BS@1.4kb | 1438-(6xMS2BS insertion)-1439 | This study | GCATCGTACGCGTACGTGTT | AGCAGATGCATCCGGCTCGA | HAP1 WT |
| NEAT1/6xMS2BS@14kb | 14020-(6xMS2BS insertion)-14021 | This study | GCATCGTACGCGTACGTGTT | TACCGCATATCTGTGTACAT | HAP1 WT |

Genotyping PCR in Extended Data Fig. 2a

| Primer name | Sequence (5' to 3') |
|---|---|
| hNEAT1_#1 (748F) | CCTAGCATGTTTGACAGGCG |
| hNEAT1_#2 (1276R) | CATCAATTGAGGCGGGCTCT |
| hNEAT1_#3 (7930F) | CCTCGGCAGTGAGCACTTAT |
| hNEAT1_#4 (8396R) | AGACGCATCTCGAGGACACT |
| hNEAT1_#5 (16520F) | TCCCATTCTGTGGGTTGTGT |
| hNEAT1_#6 (17009R) | ACACCTGTAGTCTGATAGGGGA |
| hNEAT1_#7 (22280F) | CCCTCACCTATCCCACCCTA |
| hNEAT1_#8 (+72R) | TGGAAGGAAGCAGCAACACT |

Genotyping PCR in Extended Data Fig. 4a

| Primer name | Sequence (5' to 3') |
|---|---|
| hNEAT1_#1 (748F) | CCTAGCATGTTTGACAGGCG |
| hNEAT1_#9 (8263R) | GGCTGGAGGGAATGGAATGT |
| hNEAT1_#10 (15749F) | CTAGTCTTTGGTAACCTTTG |
| hNEAT1_#11 (+65R) | AAGCAGCAACACTGCGGGGA |

Genotyping PCR in Extended Data Fig. 7a

| Primer name | Sequence (5' to 3') |
|---|---|
| hNEAT1_#12 (1208F) | CCTGGCCAGTGTGAGTCCTA |
| hNEAT1_#13 (1695R) | TGGGTAGGTGAGAGGTCACG |
| hNEAT1_#14 (13685F) | ACCCTGTGGGTGATTCAGTG |
| hNEAT1_#15 (14179R) | GGGTGAAGGTGTGGGGAAAT |

| Synthesized RNAs for RNA pulldown | |
|---|---|
| Name | Sequence (5` to 3') |
| NEAT1_8-9.2kb S | gggcgaattgggccctctagatgcatgctcgagcggccgccagtgtgatggatat<br>ctgcagaattcgcccttGTGAGTATATGATATCCATTTCCCTACAT<br>AGCCACTAACATCAGGTTTTTACAATTTTATTTATTTCTTG<br>CTACTTTAAGAAATTTTTGTGGTGAAATACATATAATAGAA<br>GTTGACTATCTGAATCATTTTTAAGTATACATTCAGTAGTG<br>TTAAGTATGTCGCCATTGTTGTACAACCAATCTCCAGAAC<br>TTTTTCATCTTGCAAAACAAACTCTGTACCCATTAAATAAC<br>ATTAAACATTCCATTCCCTCCAGCCTCAGCAACCCCATT<br>CTACTTTCTGTTTCTGTGAGTTTGACTATTCCAAGCACTTC<br>ATATCAGTTAAATCATGAAGTATTTGTCTGTCTGTGACTGG<br>CTTATTTCTCTGAGCACAGTGTCCTCGAGATGCGTCTATG<br>TTGTAGCATATGTCAGAATTTCCTTCCTTTTTAAAAGATCC<br>AAATAATATTCTTATTTTATATCTTTTTTTTATCCATTCATCC<br>ATTAGTGGACACTTGGGTTGCTTTTGGCTATTGTAAATAAT<br>GGTGCTATGTACAAATATCTATATTATTGTATTTACAAGTAT<br>AATGCTGTAATGTACACACATCTTTTTGAGATCCTACCTTC<br>AGTTCTTTTGAGTATATAGCCAGAAGTGGTATTACTAAATC<br>TTACGATATTTCTATTTTTAATTTATTGAGGAACCACTGTAG<br>TTTTTCATAGCAACTGCACCATTTTACGTTCTCACCAAGA<br>GTGCACAAGGGTTCCGAGGTTCCCACATCCTCCCCAACA<br>CTTGTTATTTTCTGCTTTTTTTAGATTGCAGCCATCATAGT<br>GGGTGTGAGGTGACATTTCATTGTGGTTTTGATTTGCATTT<br>CCCTAATGAGGAGTGATGCTGAGCATCTTTTCATATGCTT<br>ACTGGTCATTTGTATGTTGTCTTTGGAAAAATGTCTATTCA<br>AGTCCTTTGACTATTTTAAAAATTGGGTTATTAGAGTTATC<br>GTTGTTGTTGACTTGTAGGAGTTTCTTTCTATATTCTGGAT |
| NEAT1_9.2-9.8kb S | agaatactcaagctatgcatcaagcttggtaccgagctcggatccactagtaacg<br>gccgccagtgtgctggaattcgcccttTCCAACGTTATAAGGTACTTTT<br>AAGGTATTTTAGTTGTCTTAGTCTATATTTCTGTACTCACC<br>TTTCTTTATCCACTCATCAGTTGATGGGCATGTAGGTTGGT<br>TCCATATCTTTGCAATTCTGAATTGTGCTATGATCAGGTGT<br>CTTTTTAGTATAATGATTTACTCTCCTTTGGGTAGATACCC<br>AGTAGTGGGATTGCTGGATCGAATGGTTTTTATAATTTTCT<br>ATTTTACCACAGTTTCTCTCTGCATTTTTCCTCTTTGACCA<br>CTAACCATGTGAAATTCTCATATTGACCTTTATAATGATCA<br>TGAACTCTTAGTATCATTGGGAAGGCCACATTTGCCACTT<br>ATGATTGTAAACCTTATCCTCCATTTTTCCTGTTATTGTTG<br>GTGCAAAAAGCACCTATTATACCAGGACTTTAAAAATCAG<br>TCTGATAAGTCTTTGATAAGTCTAATAATAATAACTGATAA<br>GTCCATTGAATTTGCTTCTGATTACTTTTTCTTTAGTAGCTA<br>AACATGTATGTACTCCTATGATTACAATGAACACTCCTCT<br>CCATTTAAATTAATTATTTACATTGATGAAATAGCAAAATGT<br>TAATGACTAAATACTGTCTTGGTTTTTTCGTTCCAGGaaggg<br>cgaattctgcagatatccatcacactggc |

* NEAT1 derived sequences are capitalized and linker sequences are shown in lowe

| Synthesized RNAs for RNA-FISH | |
|---|---|
| Name | Sequence (5` to 3') |
| NEAT1_0-1kb-AS | agaatactcaagcttgcatgcctgcaggtcgactctagaggatccCCAACA<br>ATACCGACTCCAACAGCCACTCGGCTTACTGTCCCGG<br>GCTTACCAGATGACCAGGTAATGTTTTAAGTGAATGGAT<br>AAGTTAAAGGGAGGAAGAAGGGGTGGAGTGAGCTCACA<br>AGAAGAGTTTAGCGCCAAACCTAGAGAAAAGTCCAAAA<br>GGAGCACTGCCACCTGGAAAATAAAGCGTTGGTCAATG<br>TTGTCCCCAACACCCGAGCAGCGTGCCTCGCAGTCCC<br>CGCCTGTCAAACATGCTAGGTGCCGCACCAGTCACCG<br>GTGAGGATGGCGCCTTAACTCCACATCACTCCTCAGAC<br>CACCCCTCACCTCCGCCCAGGGGCCTGGCTTCGCAC<br>CCAGACCTGGACGCTCCACCAGGCTGGGCCCGGCCT<br>GGGCCCCGTCCCAGGCCGAGCGAAAATTACATACCAC<br>CCTGGGCCAGAGCGGTCAGCCCCGTCGAGCTAAGTTC<br>AGTTCCACAAGACCAGGCCTCATCCCCAGGCGGGTTC<br>ACAGCCCTTGGTCTGGAAAAAAAGGGGCTGCTGGCATG<br>GACAAGTTGAAGATTAGCCCTCCCGGCCCTCCTGCAG<br>CCCTGCACCCACTGCCTGCCTTCCTGATCATTTCCAGG<br>GCTGCTGCGGCCTATTCCTCCTGACTCCTCCACCCCTT<br>CTACCTCTCCCTGCCTTCCTCCTTCCACACAGACCAGC<br>GCACCCGGGCTCGCTCAGCTATGCAAGAGCGGCGCC<br>CTCCCAGAGGTCAAGTTCCCCCTCCACCAGGCAGTAG<br>GGACAGCCTGGCCTGGAGCGTGGCTGTTCTGCCTGGG<br>GGACCCTGCGGATATTTTCCATGCAGCCTGCCCCACTG<br>TGGTCCCCCTAGACCTAGTCTCCTTGCCAAGCTTCCTT<br>CTCGCACCCCCAGCCCGCCCCCTCGTCTCATCTAACT<br>CAGTCATCTCTCCCTGTCTGTCCCCTGAAGCCCTGAGC |
| NEAT1_21.7-22.6kb-AS | agaatactcaagctatgcatcaagcttggtaccgagctcggatccactagtaac<br>ggccgccagtgtgctggaattcgcccttCATAGCCAGGGGACATTGT<br>CCCCCAGCTTCACATCCACATGTCTCCTAGCATGGCAC<br>ATGCATATCCTGCCCAAGGAGCATGAAGTCAGACCAGC<br>AAACTGAACACGAGGCACAGTCCTGCATGCTCTGGGA<br>CCGAGCACTGAGGGGGCCCACAGGTGAGAAGGTGCTG<br>GCCCTTCAGCTCTCCCTCCCTCTGCCTTCACGAGCAG<br>AGGCACAAGCTTCTGTGTGTATACAGAAATTTCCAGAGT<br>GAGTCTAGGAAACTCGAGAGGACATCTCTGTGTAGTAG<br>GGTGGGATAGGTGAGGGCAGGGATTTTACTTGCATTTTA<br>TACCCATGGCTTGAATTTTTGACTCTATGCAGATCACTT<br>TTATCATTAAAGGCACAAATCTTTTCATCTGAACAGGGA<br>ATCTAAGACCTGTGGACACTTCCGGCACCCTGGCCTAG<br>TGGAAATGGTTCTCTGGGTCTGCTGGGCCCCAAGCACC<br>TGAGGTCACGCAGACGCCCGCCTGCCTCGCCTTCCAT<br>TCTCTGATCTGTCAGCCACCGTCAGCTGTGGATTCCTT<br>CCTGGGAACTGCCATCACTCGAAATGAAGTCGGAAGCA<br>CAGGGCCCCACCAGCCTTTCTCCTGTCACTTGACTCTA<br>ACCTGGAGTCCAAGTCACCAGGGCCCCAGGACCCAAA<br>GGTACACAAAGCCCACCTGGACCCTCAGCCGGTACTT<br>GCCCCAGGCCACCCTCAGGATCCACGGTCCACAGCA<br>CCACTGCTGTTGGCTCAGGCTCTGCCTCCCCTGGGCC<br>CACGGCGGGTCCGTGGGAGCCCCCCACGAGCTCTGC<br>TGTCCTCATTTGAGAGTCCTGTCCTGGCaagggcgaattctg<br>cagatatccatcacactggc |
| NEAT1_DS1-600-AS | agaatactcaagctatgcatcaagcttggtaccgagctcggatccactagtaac<br>ggccgccagtgtgctggaattcgcccttAGCACAGGCTGACTTCCCT<br>GTCCCACCCGGCACCTTGTGGTCCCAGCTGTTGGCCT<br>TCCTGCACAAGGGCCGGTTCGTTGGGCCCTGTCCGCC<br>CACCCATCTCCAGCCCAGGCCTCAGGATTTCTGGCCT<br>CTCCCCTTGGACCTCATCTCCCCAAGCCCTCCCTGGA<br>AGAGCCACGCTGGTCCTCAGATGTGGAGAGGAGGGTA<br>GCGGTGAGTTGATCCTGGGAAGAAGAGGGCAGTGGGG<br>AGTATGCCCTGAGGGGTGGGGACACTCCAGGTAGGCC<br>CCACAGCTTTGGGTGCAGAGCCGCCAAGCTGTCCCCC<br>TCCCGCTCCCACAGGAGTGGCTGAGGGGGCTAAGTTA<br>TCAGCTGCCTGTGGCCCCACGTCCCCTGCTGTATTCC<br>CAGGGCCCCAGAGGATGAGGGAGGGGATAGCAGGGC<br>AGGAGGGCTTCTGGGAGAGGCCTGGGGTGCCCCACCC<br>AATGCTACCCCTCTAGGAGACAAGATATCCCTGGTGGG<br>GTGGGAGGAGAGGCCTGGAGGAGGGAGCTGGAAGGAA<br>GCAGCAACACTGCGGGGACTCGAACCCCGGCCCAGC<br>CGTGCTGGACGTGCCACCACCAGCGCCaagggcgaattctg<br>cagatatccatcacactggc |
| MALAT1_4601-5219-AS | agaatactcaagctatgcatcaagcttggtaccgagctcggatccactagtaac<br>ggccgccagtgtgctggaattcgcccttaAGATGGACATTGCCTCTT<br>CATTGTATTTCTCATCAATTCATTATTTTTGTGGTTATAGC<br>TTGACAAGCAATTAACTTTAAAATGGTAGATTCCGTAAC<br>TTTAAATTGGTAGCTTTCATTTGCTTAAAATTTTTTGGCAT<br>ATGCAGATAATGTTCTCATCAGTAGTAAGAATCTCAGGG<br>TTATGCTTATTCCCCAATGGAGGTATGACATATAATCTTT<br>TCTGCCTTTACTTATCAATTCACCAAGGAGCTGTTTTCT<br>CTGCATCTAGGCCATCATACTGCCAGGCTGGTTATGAC<br>TCAGAAGATGTTATCTGAAAAAAGTCTATAGAAAAAAAA<br>AAAGTTTCCCCTCCCTCATCAACAAAAGCCCACCCTCT<br>AAGAGACATTCAAGCTGAACTATCACAATTCTTAATCAG<br>TTACAATTTACAAACAGATAAGTTTAAAATAAACAATTTA<br>CAAAATTTTTGAAGCATACCTTAACATCTTGTTTTGCAGT<br>TAAACAATGGAAAAGTATTTCTCCTACACTAAAAAAAAA<br>CTTGCTTACACACAACTGAAAATAGAATCTTACTTGATA<br>ATACAAAAGCTACCATCAGAAGAAATCCCTTCAGGATC<br>ATTAAGCCACTcggtgaagggcgaattctgcagat |

* NEAT1 derived sequences are capitalized and linker sequences are shown in lowe

| Name | The amino acid sequence |
|---|---|
| SFPQ Q to G mutant | MSRDRFRSRGGGGGGFHRRGGGGGRGGLHDFRSPPPGMGLN<br>GNRGPMGPGPGGSGPKPPIPPPPPHGGGGGPPPGGPPPGGPP<br>PHGPPPHPGPHGGGGPPPPPGDSSKPVVAGGPGPAPGVGSAP<br>PASSSAPPATPPTSGAPPGSGPGPTPTPPPAVTSAPPGAPPPTP<br>PSSGVPTTPPGAGGPPPPPAAVPGPGPGPKGGPGPGGPKGGK<br>MPGGPKPGGGPGLSTPGGHPKPPHRGGGEPRGGRGHHPPYHG<br>GHHGGPPPGGPGGRSEEKISDSEGFKANLSLLRRPGEKTYTQRC<br>RLFVGNLPADITEDEFKRLFAKYGEPGEVFINKGKGFGFIKLESRA<br>LAEIAKAELDDTPMRGRQLRVRFATHAAALSVRNLSPYVSNELLE<br>EAFSQFGPIERAVVIVDDRGRSTGKGIVEFASKPAARKAFERCSEG<br>VFLLTTTPRPVIVEPLEQLDDEDGLPEKLAQKNPMYQKERETPPR<br>FAQHGTFEYEYSQRWKSLDEMEKQQREQVEKNMKDAKDKLESE<br>MEDAYHEHQANLLRQDLMRRQEELRRMEELHNQEMQKRKEMQ<br>LRQEEERRRREEEMMIRQREMEEQMRRQREESYSRMGYMDPRE<br>RDMRMGGGGAMNMGDPYGSGGQKFPPLGGGGGIGYEANPGV<br>PPATMSGSMMGSDMRTERFGQGGAGPVGGQGPRGMGPGTPA<br>GYGRGREEYEGPNKKPRF |
| SFPQ P to A partial mutant | MSRDRFRSRGGGGGGFHRRGGGGGRGGLHDFRSAAAGMGLN<br>QNRGPMGPGPGQSGPKAAIAAAAAHQQQQQAAAQQAAAQQAA<br>AHQAAAHPQPHQQQQAAAAAQDSSKPVVAQGPGPAPGVGSAA<br>AASSSAAAATAATSGAAAGSGPGPTPTAAAAVTSAAAGAAAATA<br>ASSGVPTTAAQAGGAAAAAAAVPGPGPGPKQGPGPGGPKGGK<br>MPGGPKPGGGPGLSTPGGHPKAAHRGGGEPRGGRQHHAAYHQ<br>QHHQGAAAGGPGGRSEEKISDSEGFKANLSLLRRPGEKTYTQRC<br>RLFVGNLPADITEDEFKRLFAKYGEPGEVFINKGKGFGFIKLESRA<br>LAEIAKAELDDTPMRGRQLRVRFATHAAALSVRNLSPYVSNELLE<br>EAFSQFGPIERAVVIVDDRGRSTGKGIVEFASKPAARKAFERCSEG<br>VFLLTTTPRPVIVEPLEQLDDEDGLPEKLAQKNPMYQKERETPPR<br>FAQHGTFEYEYSQRWKSLDEMEKQQREQVEKNMKDAKDKLESE<br>MEDAYHEHQANLLRQDLMRRQEELRRMEELHNQEMQKRKEMQ<br>LRQEEERRRREEEMMIRQREMEEQMRRQREESYSRMGYMDPRE<br>RDMRMGGGGAMNMGDPYGSGGQKFPPLGGGGGIGYEANPGV<br>PPATMSGSMMGSDMRTERFGQGGAGPVGGQGPRGMGPGTPA<br>GYGRGREEYEGPNKKPRF |
